# Electrical Brain Activity during Human Walking with Parametric Variations in Terrain Unevenness and Walking Speed

**DOI:** 10.1101/2023.07.31.551289

**Authors:** Chang Liu, Ryan J. Downey, Jacob S. Salminen, Sofia Arvelo Rojas, Natalie Richer, Erika M. Pliner, Jungyun Hwang, Yenisel Cruz-Almeida, Todd M. Manini, Chris J. Hass, Rachael D. Seidler, David J. Clark, Daniel P. Ferris

## Abstract

Mobile brain imaging with high-density electroencephalography (EEG) can provide insight into the cortical processes involved in complex human walking tasks. While uneven terrain is common in the natural environment and poses challenges to human balance control, there is limited understanding of the supraspinal processes involved with traversing uneven terrain. The primary objective of this study was to quantify electrocortical activity related to parametric variations in terrain unevenness for neurotypical young adults. We used high-density EEG to measure brain activity when thirty-two young adults walked on a novel custom-made uneven terrain treadmill surface with four levels of difficulty at a walking speed tailored to each participant. We identified multiple brain regions associated with uneven terrain walking. Alpha (8 - 13 Hz) and beta (13 - 30 Hz) spectral power decreased in the sensorimotor and posterior parietal areas with increasing terrain unevenness while theta (4 - 8 Hz) power increased in the mid/posterior cingulate area with terrain unevenness. We also found that within stride spectral power fluctuations increased with terrain unevenness. Our secondary goal was to investigate the effect of parametric changes in walking speed (0.25 m/s, 0.5m/s, 0.75 m/s, 1.0 m/s) to differentiate the effects of walking speed from uneven terrain. Our results revealed that electrocortical activities only changed substantially with speed within the sensorimotor area but not in other brain areas. Together, these results indicate there are distinct cortical processes contributing to the control of walking over uneven terrain versus modulation of walking speed on smooth, flat terrain. Our findings increase our understanding of cortical involvement in an ecologically valid walking task and could serve as a benchmark for identifying deficits in cortical dynamics that occur in people with mobility deficits.

## Introduction

Human bipedal locomotion is inherently unstable. Even when walking on a smooth, level surface, the body’s center of mass moves outside of the base of support and thus poses a challenge to maintain dynamic balance. The natural environment surrounding us is rarely flat. Walking over uneven terrain poses additional challenges to bipedal locomotion as people need to be aware of the external environment (e.g., irregular rocks, grass, slippery surfaces) and actively control their balance to prevent falls. Prior studies have found changes in biomechanical measures when walking over uneven terrain compared to flat terrain, including an increase in kinematic variability, a decrease in gait stability, and an increase in energetic cost [1–5].

While walking on a flat surface is usually considered automatic and requires little contribution from the brain [6,7], performing complex walking tasks requires top-down cortical control at higher-order brain centers (reviewed by [8]). Cognitive processes such as motor planning, motor execution, and error detection are necessary for walking over uneven terrain and avoiding obstacles [9–11]. For example, when walking over some types of uneven terrain, people often plan foot placement based on visual information [12,13]. People actively monitor their motor performance using multi-sensory feedback to generate corrective responses to control their balance. Traditionally, there has been a lack of brain imaging techniques enabling high-quality brain data during human locomotion.

The recent development of mobile imaging techniques using high-density electroencephalography (EEG) has enabled direct, non-invasive measurement of cortical activity during whole-body movement with millisecond temporal resolution. For example, there is building evidence for consistent changes in electrocortical activity with challenges to balance control [14–19]. Compared to normal walking on a smooth, level surface, alpha (8 - 13 Hz) and beta (13 - 30 Hz) spectral power at the sensorimotor area was lower when walking on a ramp [16], walking on a narrow beam [17], and walking on uneven grass terrain [15]. Beta spectral power at the premotor area was lower when participants were not given external stability support during walking [14]. Additionally, theta (4 - 8 Hz) spectral power was greater during narrow beam walking compared with treadmill walking [17]. Theta power increased sharply when participants stepped off the balance beam [17,19,20]. Each of these EEG spectral power bands (theta, alpha, beta) provide unique insight into brain function during motor tasks (reviewed by [21]). Taken together, these results suggest that an increase in theta power and a decrease in alpha and beta power are associated with increasing demand for balance control during gait.

However, few studies have investigated how electrocortical activities change during walking with parametrically varied complexity, which could lead to a better understanding of neural compensation in populations with mobility deficits for future studies [8].

In many studies examining balance control of walking, an important confounding factor across studies is walking speed [2,22]. Faster walking speeds require greater muscle activation, mechanical energy, and metabolic energy expenditure [23,24]. There is mixed evidence as to whether increasing walking speed increases, decreases, or does not alter gait stability [25–27]. Two previous studies found that alpha and beta band spectral power at sensorimotor and posterior parietal areas was lower for faster walking speeds compared to slower walking speeds [28,29]. Yet, both studies had relatively small sample sizes and did not include slow walking speeds (i.e., < 0.5 m/s).

The primary goal of this study was to determine how electrocortical activity measured by EEG changed with parametric changes in terrain unevenness for neurotypical young adults. We used a high-density EEG system to measure brain activities when young adults walked on a novel uneven terrain treadmill surface with four levels of difficulty at a walking speed tailored to each participant. We hypothesized that alpha and beta spectral power would be lower with greater terrain unevenness at the sensorimotor and posterior parietal area due to the increasing demand for balance control over uneven terrain. We also hypothesized that theta spectral power would be greater with greater terrain unevenness in the anterior cingulate area due to the increasing demand to monitor motor performance. Related to our hypotheses, we expected to find greater intra-stride spectral power fluctuations on more uneven terrains due to more cortical processing of sensorimotor information and motor adjustments within the gait cycle. The secondary goal was to determine how electrocortical activity measured by EEG changed with walking speed.

We hypothesized that alpha and beta spectral power would be lower at the sensorimotor and posterior parietal areas. Intra-stride spectral power fluctuations would be greater in the alpha and beta power band at slower walking speeds because of the need to consciously adapt to the required slow walking speed. In addition to testing our main hypotheses, we performed an exploratory analysis on other brain areas such as supplementary motor area, occipital area for uneven terrain and speed effects. The results from this study will increase our understanding of brain activity during an ecologically valid uneven terrain walking task and serve as a benchmark for future studies on population with mobility deficits.

## Results

We recorded brain activity using high-density EEG in n = 32 young adults. Participants completed four treadmill walking trials on different levels of uneven terrain (flat, low, medium, high), one seated resting trial, and four walking trials at different speeds (0.25, 0.5, 0.75, 1 m/s) (see Methods; Fig. 1A). Simultaneously, we recorded ground reaction force at each foot with capacitive shoe insole sensors to detect gait events, and we recorded sacrum kinematics with an inertial measurement unit (IMU). We also acquired a T_1_ – weighted magnetic resonance image (MRI) on a separate visit to create individual-specific head models for EEG source localization.

**Fig. 1:**
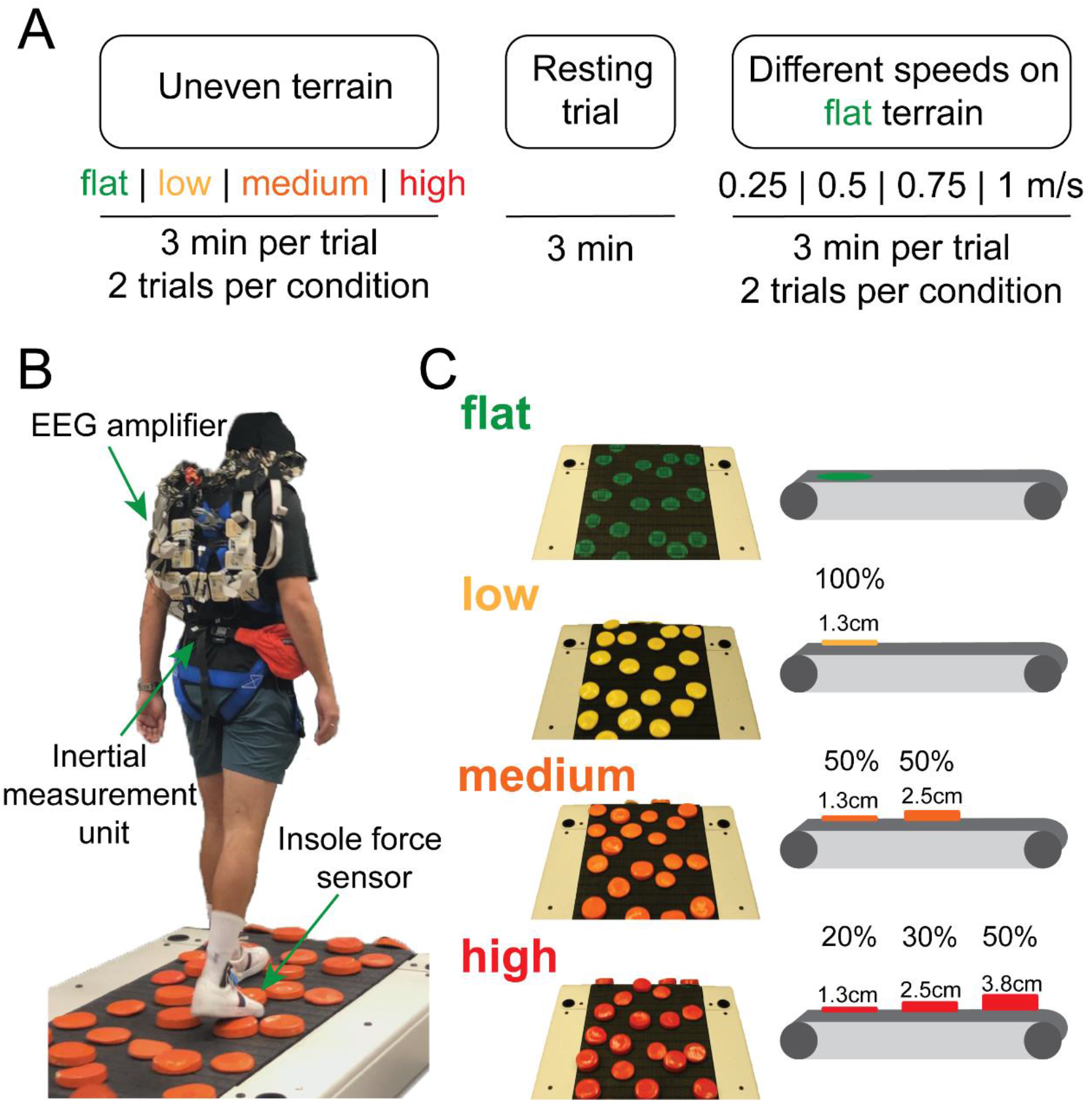
Experimental protocol and setup. (**A**) Participants completed treadmill walking trials on four different levels of uneven terrain (flat, low, medium, high), one seated resting trial, and walking trials at four different speeds (0.25, 0.50, 0.75, 1.0 m/s) performed on the flat terrain. Participants completed a block of two treadmill walking trials per condition. Each trial was 3 minutes. (**B**) Experimental setup. (**C**) Design of uneven terrain treadmill. Low, medium, and high terrain unevenness was achieved by altering the height of the rigid disks on the treadmill. Percentage refers to the proportion of disks that were the specified height.

We used EEGLAB [30] to preprocess EEG data with built-in and custom functions to remove motion artifacts and neck muscle contamination (Fig. 2A - B). After source localization, we clustered brain components based on the estimated dipole source location. Lastly, we computed the averaged power spectral density (PSD) and event-related spectral perturbations (ERSPs) tied to gait events for each component within each cluster.

**Fig. 2:**
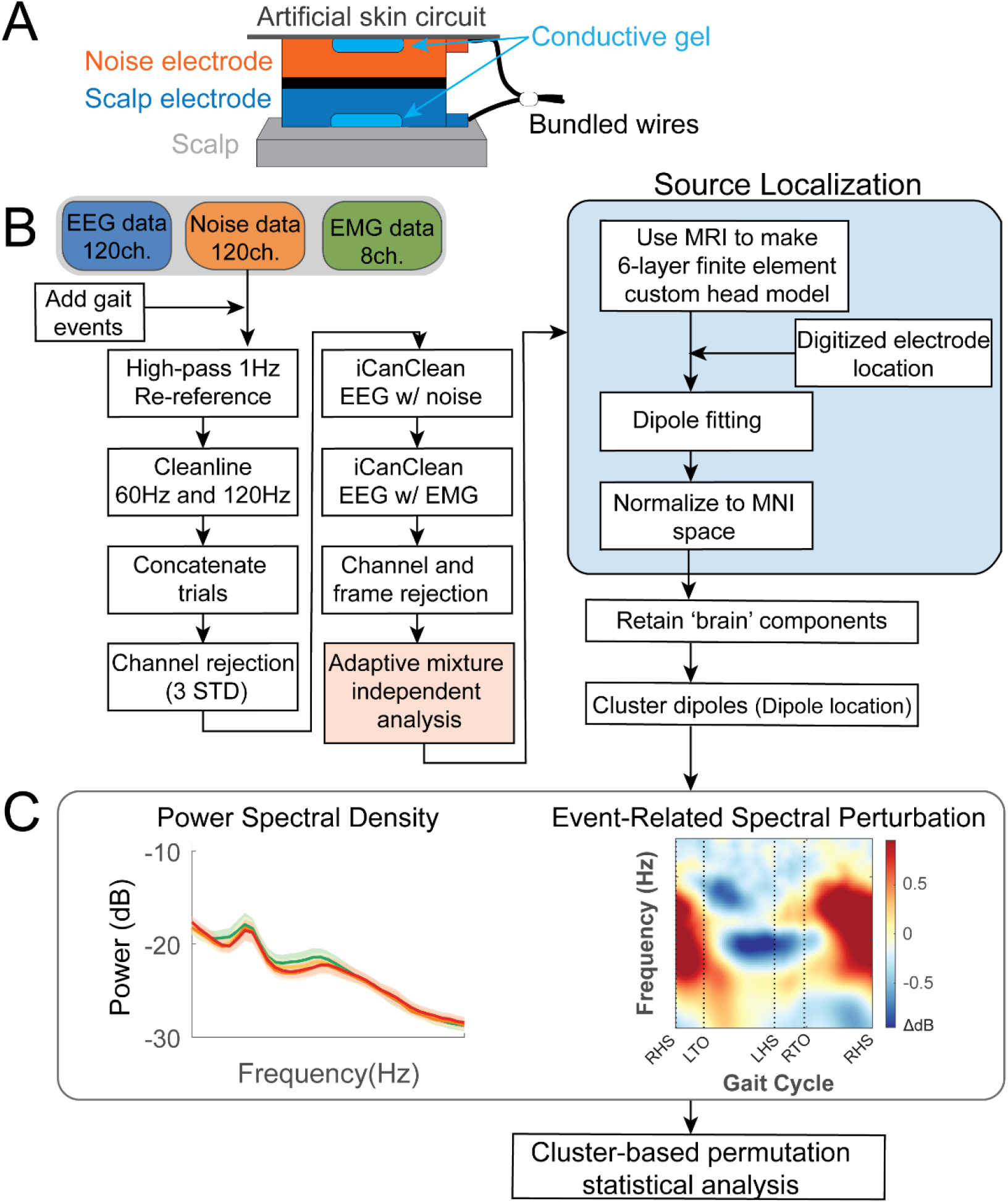
EEG processing pipeline. (**A**) Dual-electrode EEG setup. Scalp electrodes and noise electrodes were mechanically coupled. We used a conductive fabric as an artificial skin circuit to bridge the noise electrodes. Figure is by courtesy of Amanda Studnicki. (**B**) Data processing flowchart with steps for EEG pre-processing, source localization, clustering brain components. (**C**) We performed the analysis in the frequency domain after clustering brain components to investigate how electrocortical activity changes with terrain unevenness and speed. We averaged power spectral density (PSD) and event-related spectral perturbations (ERSPs) tied to gait events (RHS: right heel strike; LTO: left toe off; LHS: left heel strike; RTO: right toe off) within each brain cluster.

### Behavioral analysis

Behavioral results demonstrated that the novel, custom-made uneven terrain treadmill successfully increased gait kinematics variability and decreased gait stability. We assessed the effect of terrain unevenness on behavioral measures, including step duration, step duration variability, and sacral excursion variability in the anteroposterior and mediolateral direction in young adults (n = 32; Fig. 3) using a linear mixed effect model for each outcome measure. The mixed-effect models included Terrain (flat, low, medium, high) as an independent variable and walking speed as a covariate. We found a significant main effect on step duration (F(3, 123) = 6.2, p = 0.0006, Cohen’s f^2^ = 0.20; Fig. 3A) and step duration coefficient of variation (F(3, 123) = 213.6, p < 0.001, Cohen’s f^2^ = 8.7; Fig. 3B). Post-hoc pairwise comparison corrected by false discovery rate (FDR) indicated that longer step duration and greater step duration coefficient of variation was associated with greater terrain unevenness (Table 1).

**Fig. 3:**
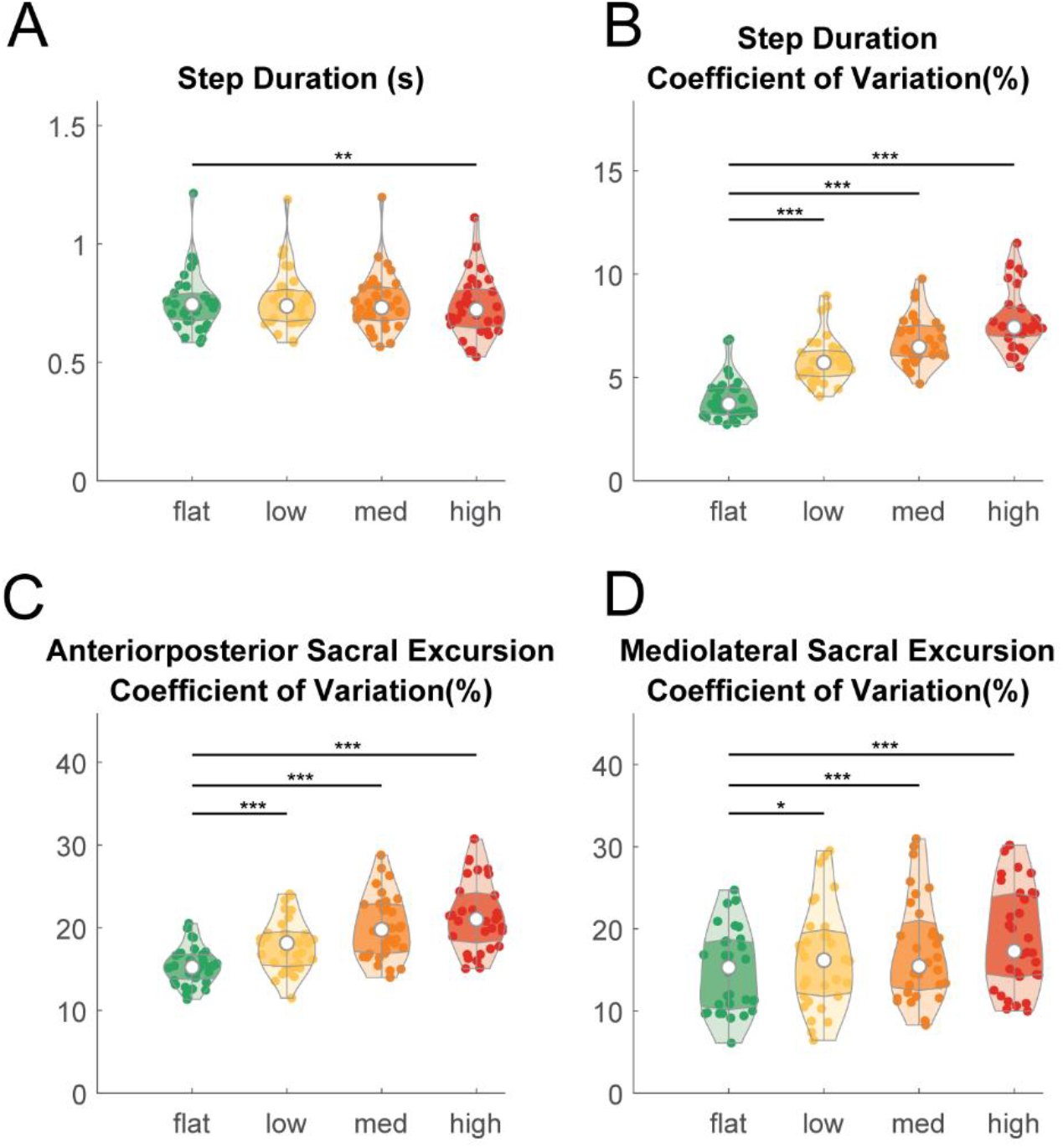
Violin plot shows the behavioral measures during walking on different levels of terrain unevenness and at different speeds. (**A**) Step duration time at flat, low, medium, and high terrain conditions. (**B**) Step duration coefficient of variation at different terrains. Sacral excursion coefficient of variation in the anteroposterior (**C**) and mediolateral (**D**) direction. The shaded regions represent data distribution across participants by estimating the probability density function. The white dots represent the median of data. The darker shaded region represents the 25 to 75 percentiles of the data. Individual data points are plotted as small dots. We only showed pairwise comparison statistics relative to the flat terrain. Refer to S1 Table for full pairwise comparison results. (* pFDR_adjusted_ < 0.05, ** pFDR_adjusted_ < 0.01, *** pFDR_adjusted_ < 0.001).

**Table 1:** Statistical results from the pairwise comparison examining the effects of terrain unevenness on each dependent variable.

| Dependent Variable | Condition 1 | Condition 2 | DF | t value | P value | P<sub>FDR-adjusted</sub> Value |
| --- | --- | --- | --- | --- | --- | --- |
| **Step Duration (s)** | flat | low | 123 | 1.08 | 0.28 | 0.34 |
|  | flat | med | 123 | -0.39 | 0.7 | 0.7 |
|  | flat | high | 123 | -3.05 | 0.0027 | **0.0082** |
|  | low | med | 123 | -1.5 | 0.14 | 0.22 |
|  | low | high | 123 | -4.1 | <0.001 | **<0.001** |
|  | med | high | 123 | -2.7 | 0.0087 | **0.017** |
| **Step Duration CoV (%)** | flat | low | 123 | 11.7 | <0.001 | **<0.001** |
|  | flat | med | 123 | 17.8 | <0.001 | **<0.001** |
|  | flat | high | 123 | 24.3 | <0.001 | **<0.001** |
|  | low | med | 123 | 6.09 | <0.001 | **<0.001** |
|  | low | high | 123 | 12.6 | <0.001 | **<0.001** |
|  | med | high | 123 | 6.5 | <0.001 | **<0.001** |
| **Anteroposterior excursion CoV (%)** | flat | low | 123 | 4.1 | <0.001 | **<0.001** |
|  | flat | med | 123 | 7.6 | <0.001 | **<0.001** |
|  | flat | high | 123 | 9.9 | <0.001 | **<0.001** |
|  | low | med | 123 | 3.5 | <0.001 | **<0.001** |
|  | low | high | 123 | 5.8 | <0.001 | **<0.001** |
|  | med | high | 123 | 2.3 | 0.026 | **0.026** |
| **Mediolateral excursion CoV (%)** | flat | low | 123 | 2.6 | 0.01 | **0.015** |
|  | flat | med | 123 | 4.1 | <0.001 | **<0.001** |
|  | flat | high | 123 | 6.3 | <0.001 | **<0.001** |
|  | low | med | 123 | 1.5 | 0.15 | 0.15 |
|  | low | high | 123 | 3.7 | <0.001 | **<0.001** |
|  | med | high | 123 | 2.2 | 0.026 | **0.031** |

We also found a significant main effect of terrain on sacral excursion coefficient of variation in both the anteroposterior direction (F(3, 123) = 37.2, p < 0.001, Cohen’s f^2^ = 1.51; Fig. 3C) and mediolateral direction (F(3, 123) = 14.1, p < 0.001, Cohen’s f^2^ = 0.51; Fig. 3D). Greater sacral excursion variability was associated with greater terrain unevenness in both anteroposterior and mediolateral directions (Table 1).

### EEG source analysis

We identified multiple neural sources that contained dipoles from more than half of the participants (n > 16). These dipole clusters were located at right sensorimotor (n = 21), left sensorimotor (n = 28), right premotor (n = 24), left pre-supplementary motor (n = 24), right posterior parietal (n = 28), left posterior parietal (n = 27), occipital (n = 20), mid/posterior cingulate (n = 24), caudate (n = 20), left temporal area (n = 17), and precuneus (Fig. 4, Table 2).

**Fig. 4:**
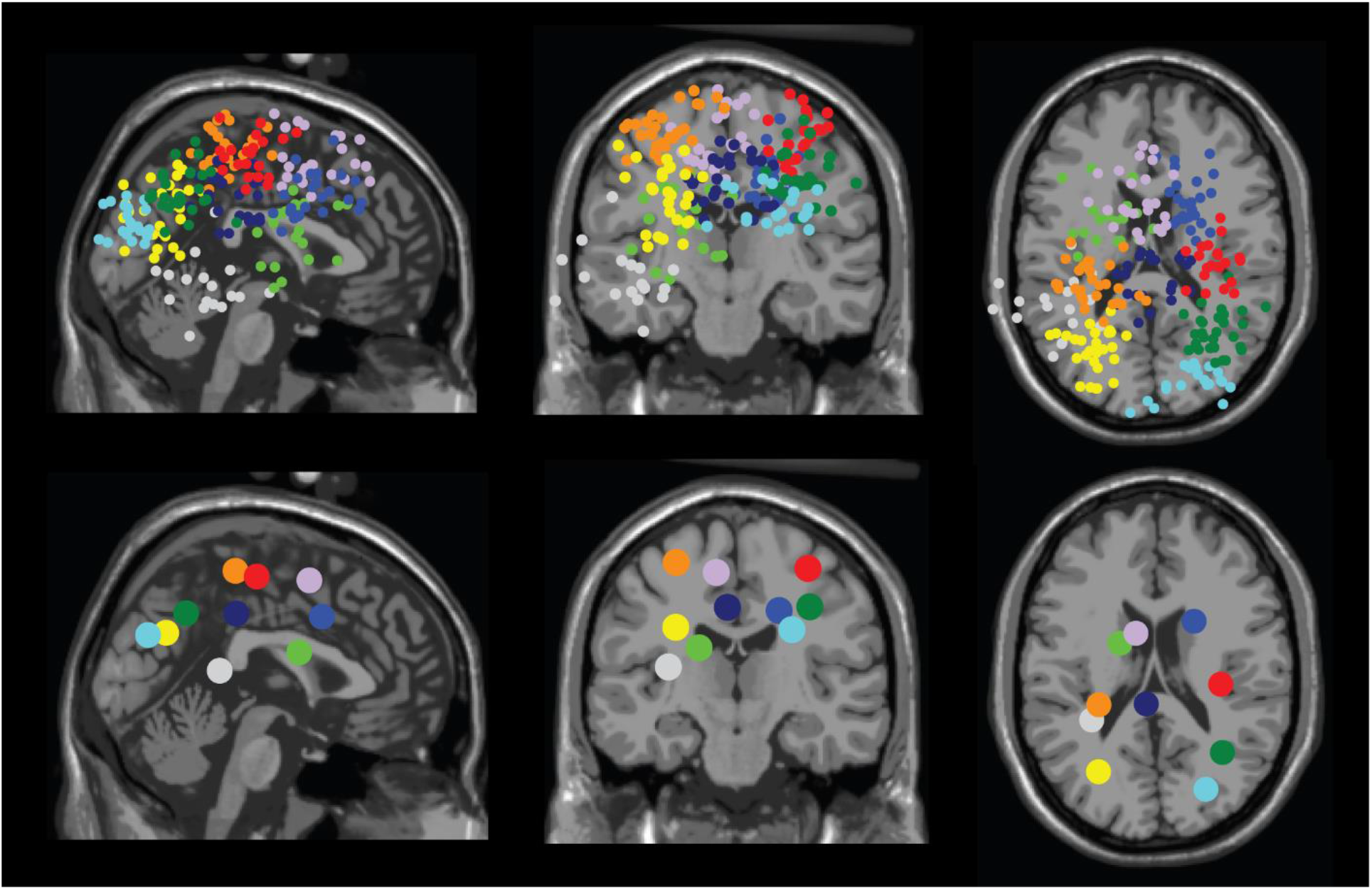
Dipole location for all participants (top row) and centroid of each cluster (bottom row) in axial (left), sagittal (middle), and coronal (right) planes. We identified clusters located at: right sensorimotor area (red), left sensorimotor area (orange), right premotor (medium blue), left pre-supplementary motor (purple), right posterior parietal (green), left posterior parietal (yellow), occipital area (cyan/light blue), mid/posterior cingulate (navy), caudate area (lime), and left temporal area (light gray).

**Table 2.**
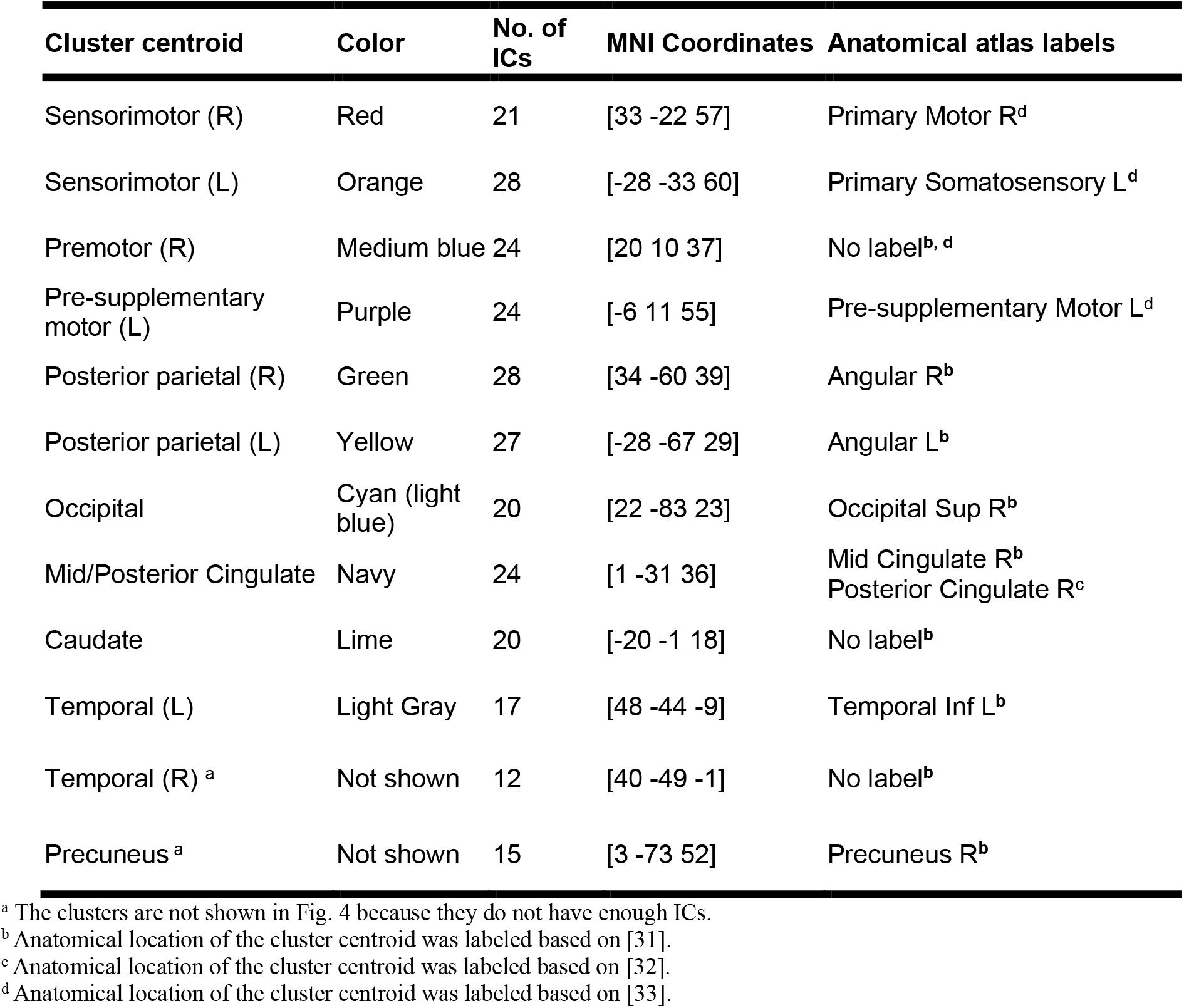
Montreal Neurological Institute (MNI) coordinates, and anatomical atlas labels for regions of interest (ROIs)

### Terrain unevenness on EEG power spectral density

We observed significant EEG power spectral modulation by terrain unevenness at multiple cortical areas (Fig.5 - 8). We decomposed the original power spectral densities (PSDs) into non- oscillatory components (modeled by the aperiodic fit) and oscillatory components shown as flattened PSDs using FOOOF toolbox (for example Fig. 5C, D; [34]). We then computed the average power within each band (theta (4 – 8 Hz), alpha (8 – 13 Hz), beta (13 – 30 Hz)) after flattening PSDs. We used a linear mixed effect model with Terrain as an independent variable to determine how terrain unevenness affected average EEG power within each band at each cluster. Post-hoc comparisons were referenced to the flat condition.

**Fig. 5:**
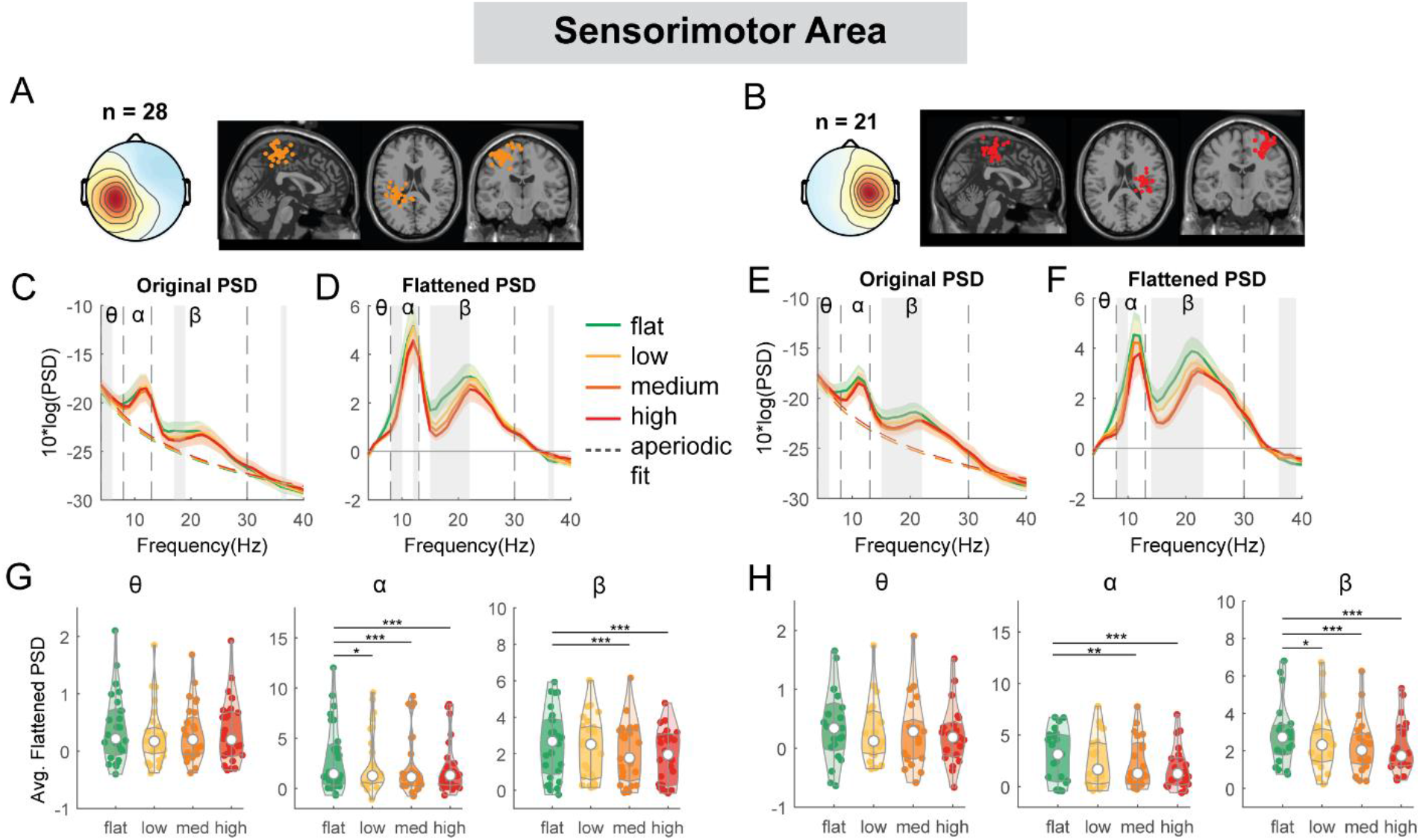
Power spectral density changes with terrain unevenness at the left and right sensorimotor area. (**A, B**) Average scalp topography within the cluster and dipole locations for each component plotted on the Montreal Neurological Institute template for the left and right sensorimotor cluster. Average original PSDs changed with terrain unevenness for the left (**C**) and right (**E**) sensorimotor cluster. Shaded colored areas indicated standard error of PSDs across components in the cluster. Dashed colored lines indicated average aperiodic fit. Gray shaded areas indicated a significant effect of terrain on PSDs. Vertical black dashed lines indicated main frequency bands of interest – theta (4 - 8Hz), alpha (8 - 13Hz), and beta (13 - 30Hz). Average flattened PSDs after removing the aperiodic fit for the left (**D**) and right (**F**) sensorimotor cluster. (**G - H**) Average power for theta, alpha, and beta band computed from flattened PSDs for all components within the cluster for the left and right clusters. Black bars and asterisks (*) indicate a significant difference in average power compared to the flat conditions (*p < 0.05, **p < 0.01, ***p < 0.001).

At both left and right sensorimotor clusters, we found a main effect of terrain on average alpha power (left: F(3,108) = 5.5, p = 0.001, Cohen’s f^2^ = 0.21; right: F(3, 80) = 6.7, p < 0.001, Cohen’s f^2^ = 0.35) and average beta power (left: F(3,108) = 11.8, p < 0.001, Cohen’s f^2^ = 0.44; right: F(3, 80) = 12.5, p < 0.001, Cohen’s f^2^ = 0.63), with both alpha and beta power decreasing with increasing terrain unevenness (Fig. 5). Compared to the flat condition, average alpha power was lower in the low (left: p = 0.02), medium (left: p < 0.001, right: p = 0.008), and high terrain conditions (left: p < 0.001, right: p < 0.001). Average beta power was also lower in the low (right: p = 0.012), medium (left: p < 0.001, right: p < 0.001), and high conditions (left: p < 0.001, right p < 0.001) compared to the flat condition.

At both left and right posterior parietal cluster, there was a main effect of terrain on average alpha power (left: F(3, 104) = 19.3, p < 0.001, Cohen’s f^2^ = 0.90; right: F(3, 108) = 24.0, p < 0.001, Cohen’s f^2^ = 1.02) and average beta power (left: F(3, 104) = 23.8, p < 0.001, Cohen’s f^2^ = 1.11; right: F(3, 108) = 24.5, p < 0.001, Cohen’s f^2^ = 1.02). Lower alpha and beta power were both associated with greater terrain unevenness (Fig. 6). Compared to the flat condition, average alpha power was lower for low, medium, and high conditions (all p < 0.001 for left and right).

**Fig. 6:**
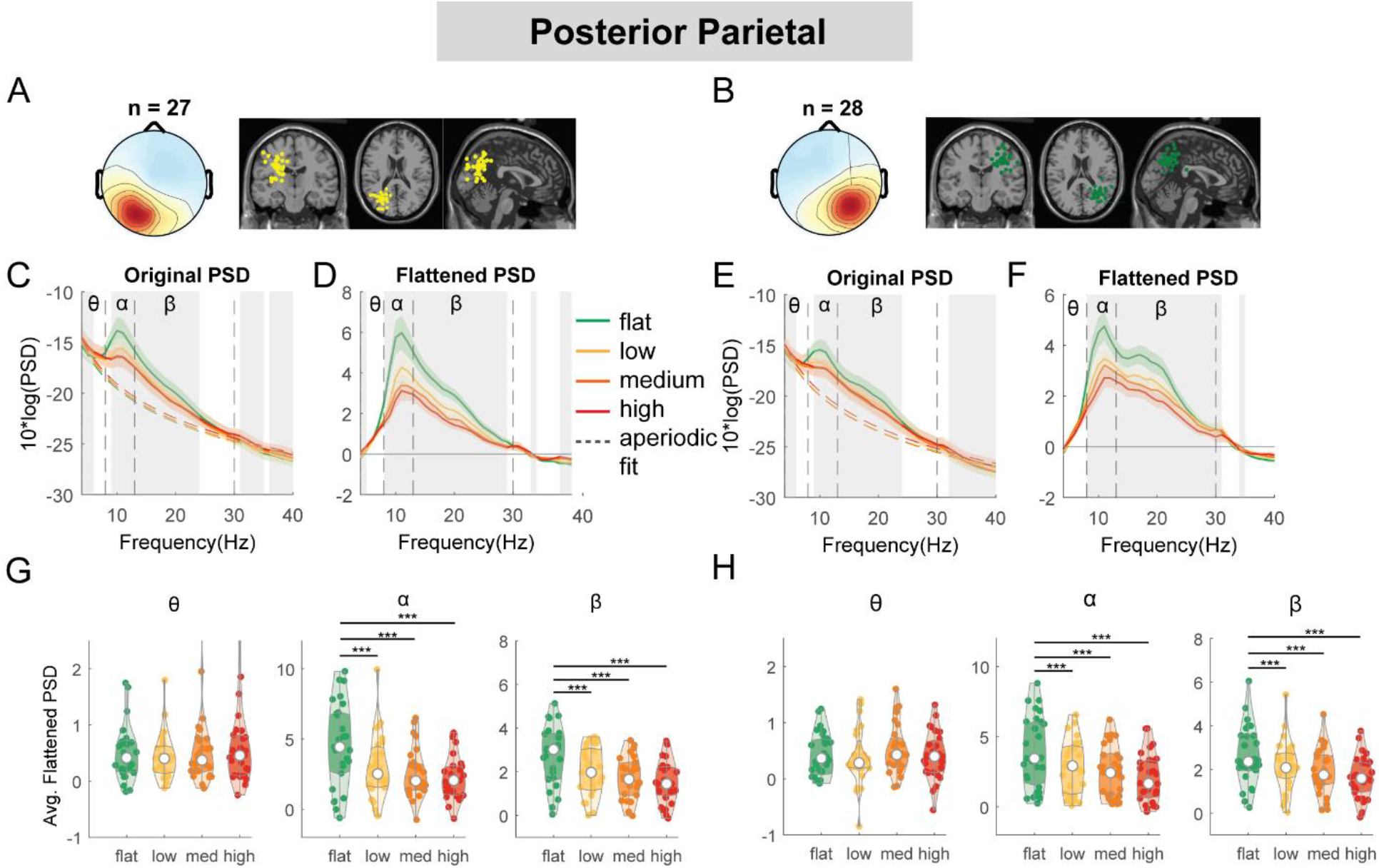
Power spectral density changes with terrain unevenness at the left and right posterior parietal area. (**A, B**) Average scalp topography within the cluster and dipole locations for each component plotted on the Montreal Neurological Institute template for the left and right posterior parietal cluster. Average original PSDs changed with terrain unevenness for the left (**C**) and right (**E**) posterior parietal cluster. Shaded colored areas indicated standard error of PSDs across components in the cluster. Dashed colored lines indicated average aperiodic fit. Gray shaded areas indicated a significant effect of terrain on PSDs. Vertical black dashed lines indicated main frequency bands of interest – theta (4 - 8Hz), alpha (8 - 13Hz), and beta (13 - 30Hz). Average flattened PSDs after removing the aperiodic fit for the left (**D**) and right (**F**) posterior parietal cluster. (**G - H**) Average power for theta, alpha, and beta band computed from flattened PSDs for all components within the cluster for the left and right clusters. Black bars and asterisks (*) indicate a significant difference in average power compared to the flat conditions (*p < 0.05, **p < 0.01, ***p < 0.001).

Average beta power was also lower in low, medium, and high conditions (all p < 0.001 for left and right) as compared to the flat condition.

The mid/posterior cingulate cluster demonstrated a main effect of terrain on average theta power (F(3, 92) = 3.5, p = 0.019, Cohen’s f^2^ = 0.16) and beta band (F(3, 92) = 16.2, p < 0.001, Cohen’s f^2^ = 0.75; Fig.7) but not on alpha power (F(3, 92) = 2.6, p = 0.055, Cohen’s f^2^ = 0.13). Theta power was greater for the high terrain condition (p = 0.01) compared to the flat terrain condition. Average beta power was lower in low (p < 0.001), medium (p < 0.001), and high conditions (p < 0.001) as compared to the flat condition.

**Fig. 7:**
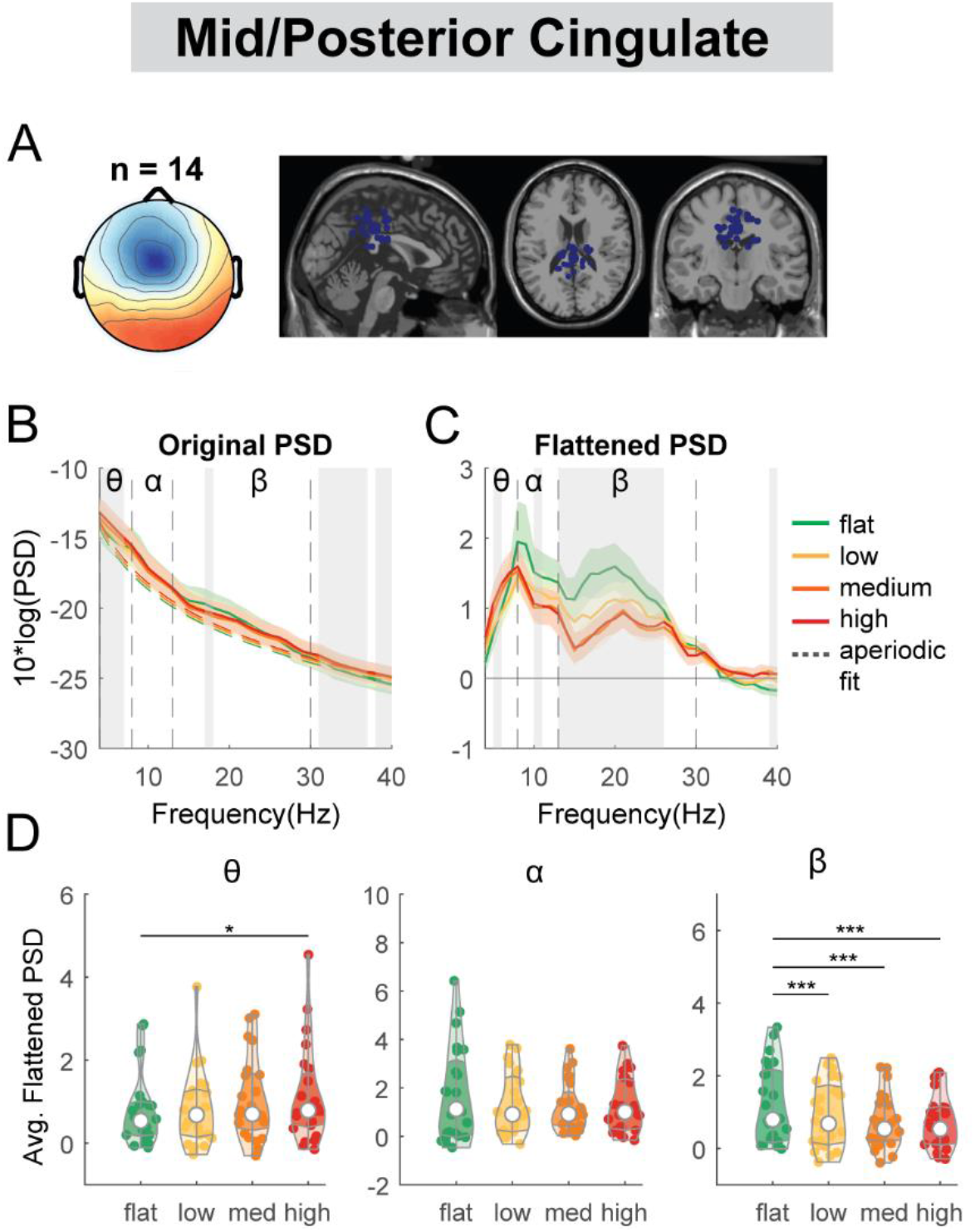
Power spectral density changes with terrain unevenness at the mid/posterior cingulate area. (**A**) Average scalp topography within the cluster and dipole locations for each component plotted on the Montreal Neurological Institute template for the mid/posterior cingulate cluster. (**B**) Average original PSDs changed with terrain unevenness for the cluster. Shaded colored areas indicated standard error of PSDs across components in the cluster. Dashed colored lines indicated average aperiodic fit. Gray shaded areas indicated a significant effect of terrain on PSDs. Vertical black dashed lines indicated main frequency bands of interest – theta (4 - 8 Hz), alpha (8 - 13 Hz), and beta (13 - 30 Hz). (**C**) Average flattened PSDs after removing the aperiodic fit. (**D**) Average power for theta, alpha, and beta band computed from flattened PSDs for all components within the cluster. Black bars and asterisks (*) indicate a significant difference in average power compared to the flat conditions (*p < 0.05, **p < 0.01, ***p < 0.001).

We also performed an exploratory analysis on the right premotor, left pre-supplementary motor, occipital, and caudate clusters. At the left pre-supplementary motor clusters, there was a main effect of terrain on average beta power (F(3, 92) = 8.2, p < 0.001, Cohen’s f^2^ = 0.37), with lower beta power associated with greater terrain unevenness (Fig. 8G). Compared to the flat condition, average beta power was lower in the low (p = 0.0070), medium (p < 0.001), and high conditions (p < 0.001). We also found an effect of terrain on average beta power (F(3, 92) = 5.3, p = 0.002, Cohen’s f^2^ = 0.24) at the right premotor cluster (Fig. 8H). Beta power was lower in the medium (p = 0.0061), and high (p < 0.001) terrain conditions.

**Fig. 8:**
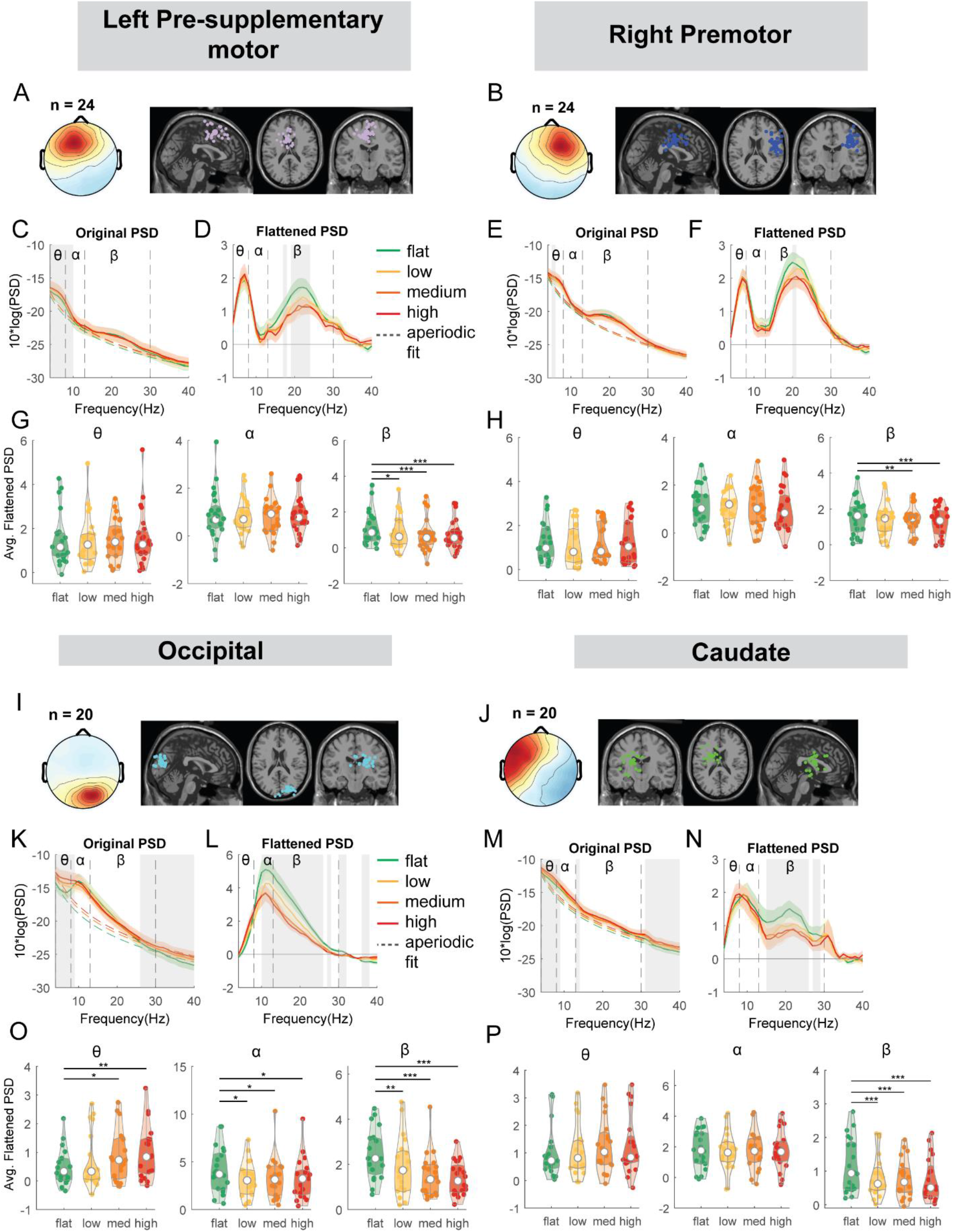
Power spectral density changes with terrain unevenness at the left pre-supplementary motor, right premotor area, occipital, and caudate area. (**A**) Average scalp topography within the cluster and dipole locations for each component plotted on the Montreal Neurological Institute template for the left pre-supplementary motor cluster. (**C**) Average original PSDs changed with terrain unevenness. Shaded colored areas indicated standard error of PSDs across components in the cluster. Dashed colored lines indicated average aperiodic fit. Gray shaded areas indicated a significant effect of terrain on PSDs. Vertical black dashed lines indicated main frequency bands of interest – theta (4-8Hz), alpha (8-13Hz), and beta (13-30Hz). (D) Average flattened PSDs after removing the aperiodic fit. (**G**) Average power for theta, alpha, and beta band computed from flattened PSDs for all components within the left pre-supplementary motor cluster. Black bars and asterisks (*) indicate a significant difference in average power compared to the flat conditions (*p < 0.05, **p < 0.01, ***p < 0.001). (**B**) Average scalp topography within the cluster and dipole locations for the right premotor cluster. (E) Average original PSDs changed with terrain unevenness. (**F**) Average flattened PSDs after removing the aperiodic fit. (**H**) Average power for theta, alpha, and beta band computed from flattened PSDs for all components within the right premotor cluster. (**I**) Average scalp topography and dipole locations for the occipital cluster. (**K**) Average original PSDs changed with terrain unevenness. (**L**) Average flattened PSDs after removing the aperiodic fit. (**O**) Average power for theta, alpha, and beta band computed from flattened PSDs for all components within the occipital cluster. (**J**) Average scalp topography and dipole locations for the caudate cluster. (**B**) Average original PSDs changed with terrain unevenness. (**N**) Average flattened PSDs after removing the aperiodic fit. (**P**) Average power for theta, alpha, and beta band computed from flattened PSDs for all components within the caudate cluster.

A main effect of terrain was found on average theta power (F(3, 76) = 4.2, p = 0.0083, Cohen’s f^2^ = 0.25), alpha power (F(3, 76) = 3.1, p = 0.02, Cohen’s f^2^ = 0.19), and beta power (F(3, 76) = 12.8, p < 0.001, Cohen’s f^2^ = 0.82) at the occipital cluster (Fig. 8O). Theta power was greater in the medium (p = 0.01) and high terrain (p = 0.0015) conditions compared to the flat condition.

Average alpha power was lower for low (p = 0.01), medium (p = 0.018), and high conditions (p = 0.016). Average beta power was lower for the low (p = 0.004), medium (p < 0.001), and high conditions (p < 0.001) as compared to the flat condition.

Lastly, we found a main effect of terrain on average beta power (F(3, 76) = 11.5, p < 0.001, Cohen’s f^2^ = 0.65) at the caudate cluster (Fig. 8P). Average beta power was lower for the low (p < 0.001), medium (p < 0.001), and high conditions (p < 0.001), compared to the flat condition.

### Gait-related spectral perturbation during uneven terrain walking

We first computed the event-related spectral perturbations (ERSPs) tied to gait events with respect to average power at each frequency across the gait cycle of the same condition [35]. We unmasked the significant deviations from the average spectrum of each condition with a bootstrap method with false discovery rate multiple comparison correction.

Alpha band and beta band ERSPs showed lateralization for left and right sensorimotor clusters (Fig. 9A, B). Gait-related alpha and beta power desynchronization occurred during the contralateral limb swing phase while alpha and beta synchronization occurred during the contralateral limb stance phase and push-off.

**Fig. 9.**
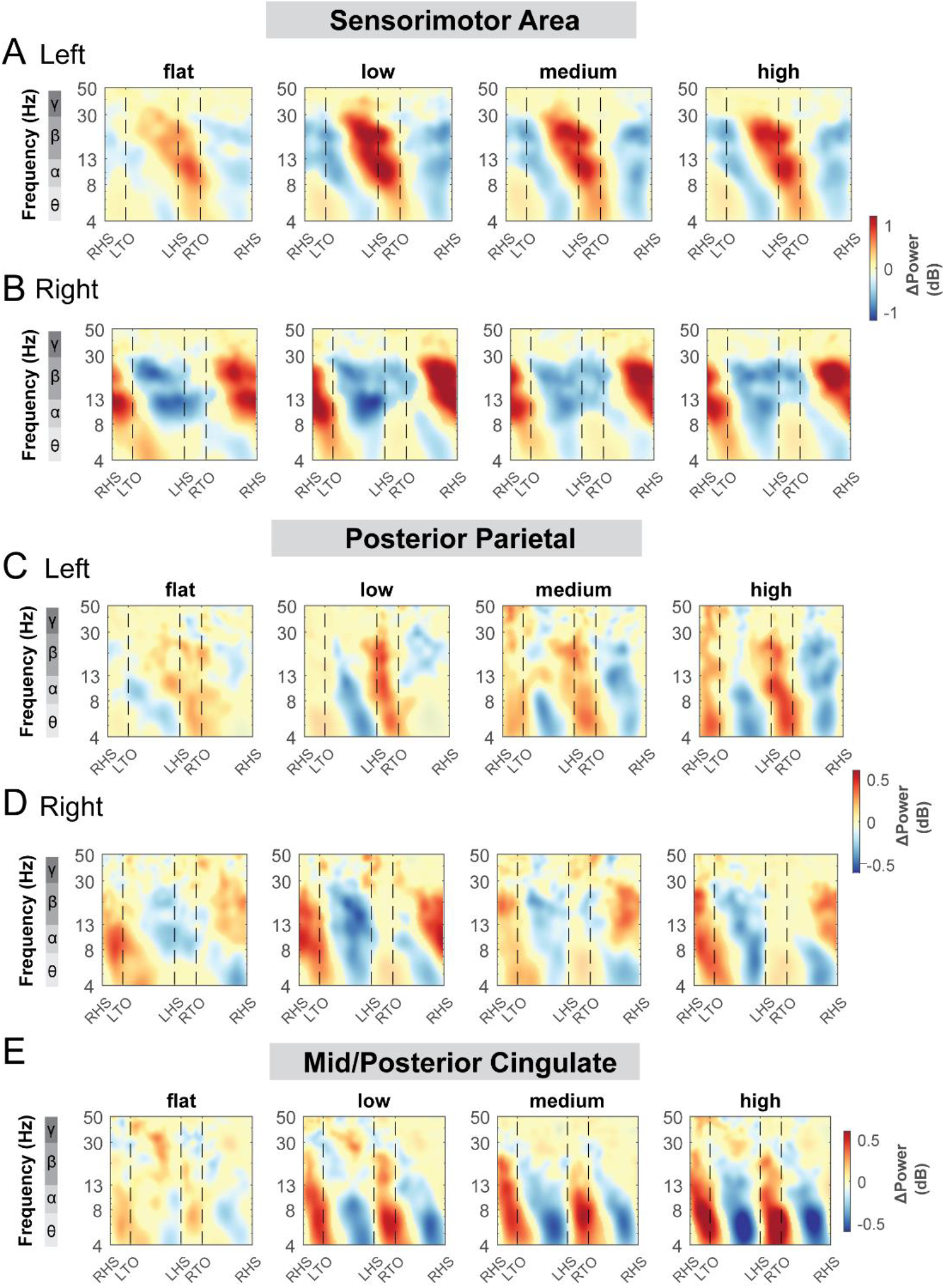
Event-related spectral perturbations at the sensorimotor (A-B), posterior parietal (C-D), and mid/posterior cingulate cluster (E) with respect to the average of each condition at different terrains. The x-axes of the ERSPs are time in gait cycle (RHS: right heel strike; LTO: left toe off; LHS: left heel strike; RTO: right toe off). All unmasked colors are statistically significant spectral power fluctuations relative to the mean power within the same condition. Colors indicate significant increases (red, synchronization) and decreases (blue, desynchronization) in spectral power from the average spectrum for all gait cycles to visualize intra-stride changes in the spectrograms. These data are significance masked (p < 0.05) through nonparametric bootstrapping with multiple comparison correction using false discovery rate.

There was also rhythmic ERSPs modulation across the gait cycle at both left and right posterior parietal clusters. For the left posterior parietal cluster, we observed theta and alpha power desynchronization during the ipsilateral swing phase and synchronization during the double support phase across all terrain conditions (Fig. 9C). As the terrain became more difficult, descriptively, we observed more broadband desynchronization during the contralateral swing phase. For the right posterior parietal cluster, we found broadband desynchronization during the ipsilateral swing phase in the low, medium, and high terrain conditions, as well as synchronization during the contralateral push-off phase (Fig. 9D).

ERSPs computed relative to the average power of the same condition at the mid/posterior cingulate areas visually increased with terrain unevenness (Fig. 9E). We found significant theta and alpha band synchronization during the double support phase and desynchronization following mid-swing when walking over uneven terrain (Fig. 9E).

We also performed similar analyses for the left pre-supplementary motor, right premotor, occipital, and caudate clusters (S1 Fig). There was synchronization during the double support phase and desynchronization around the mid-swing in theta and alpha band for the left pre-supplementary motor (S1A Fig) and right premotor area (S1B Fig). The fluctuation became more prominent during uneven terrain walking than walking on the flat surface. At the occipital area, we observed broadband (theta, alpha, and beta) synchronization during the double support phase and desynchronization around the mid-swing (S1C Fig). At the caudate area, we observed theta, alpha synchronization during the double support phase and desynchronization around the mid-swing during uneven terrain walking (S1D Fig).

**S1 Fig.**
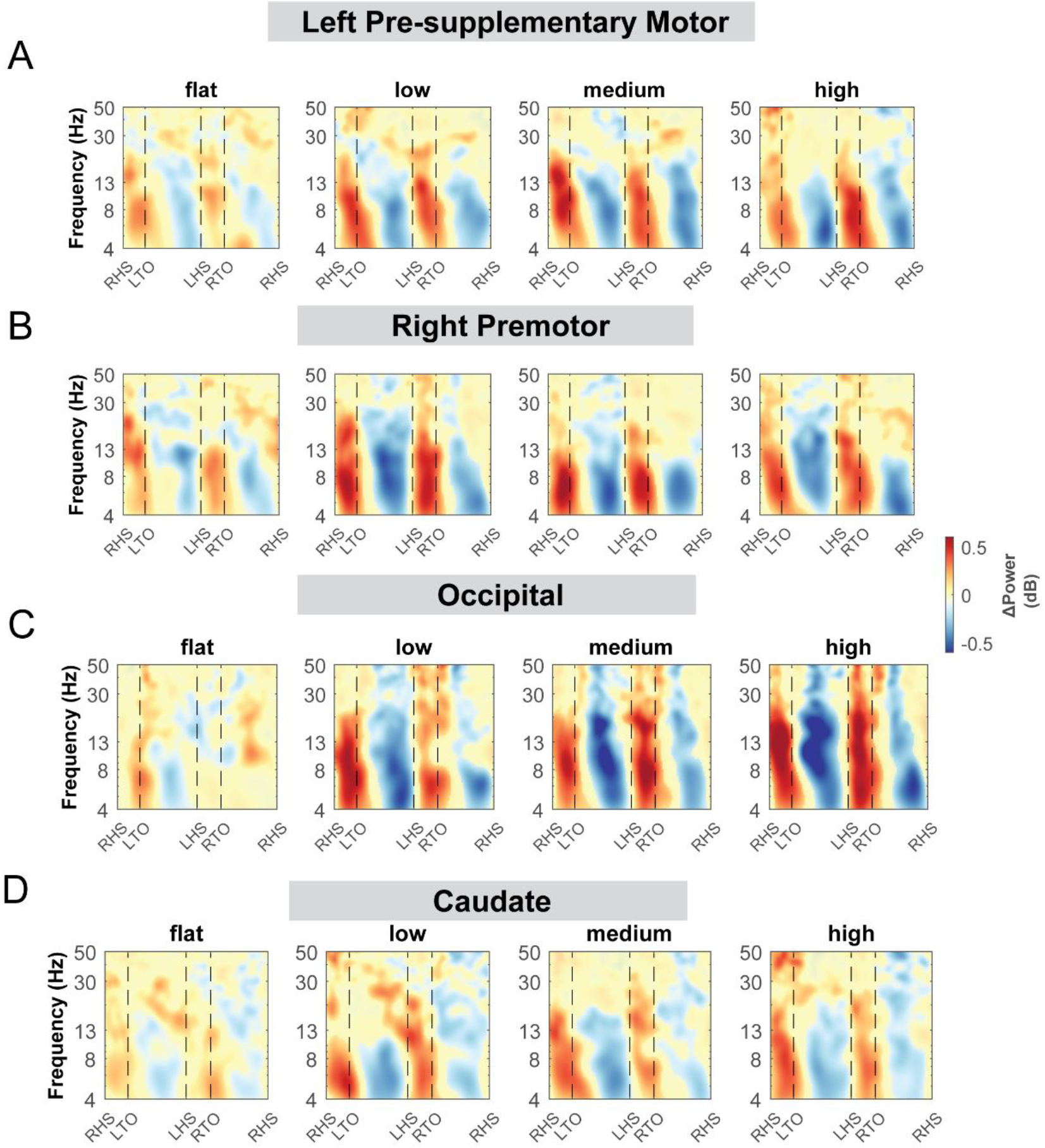
ERSPs at the left pre-supplementary motor (A), right premotor, occipital (C), and caudate clusters (D) with respect to the average of each condition at different terrains. The x-axes of the ERSPs are time in gait cycle (RHS: right heel strike; LTO: left toe off; LHS: left heel strike; RTO: right toe off). All unmasked colors are statistically significant spectral power fluctuations relative to the mean power within the same condition. Colors indicate significant increases (red, synchronization) and decreases (blue, desynchronization) in spectral power from the average spectrum for all gait cycles to visualize intra-stride changes in the spectrograms. These data are significance masked (p < 0.05) through nonparametric bootstrapping with multiple comparison correction using false discovery rate.

### Effects of terrain unevenness on event-related power perturbations

We also computed the ERSPs with respect to the grand average of all conditions to assess the effect of terrain unevenness on spectral power fluctuation tied to gait events. All clusters showed spectral power fluctuation in event-related spectral perturbation plots at various frequency bands during the gait cycle with red indicated synchronization and blue indicated desynchronization (for example, Fig. 10A). We used non-parametric permutation statistics with cluster-based multiple comparison correction to determine the time-frequency differences across terrain conditions with red color indicated significant differences across terrain conditions (p < 0.05; for example, Fig. 10B). To determine how spectral power changed relative to the flat condition, we computed the differences in ERSPs between each terrain condition relative to the flat condition (ERSP_terrain_ – ERSP_flat_) (for example, Fig. 10C). Regions that were not significantly different from flat condition have a semi-transparent mask by using permutation statistics with cluster-based multiple comparison correction (for example, Fig. 10C).

**Fig. 10:**
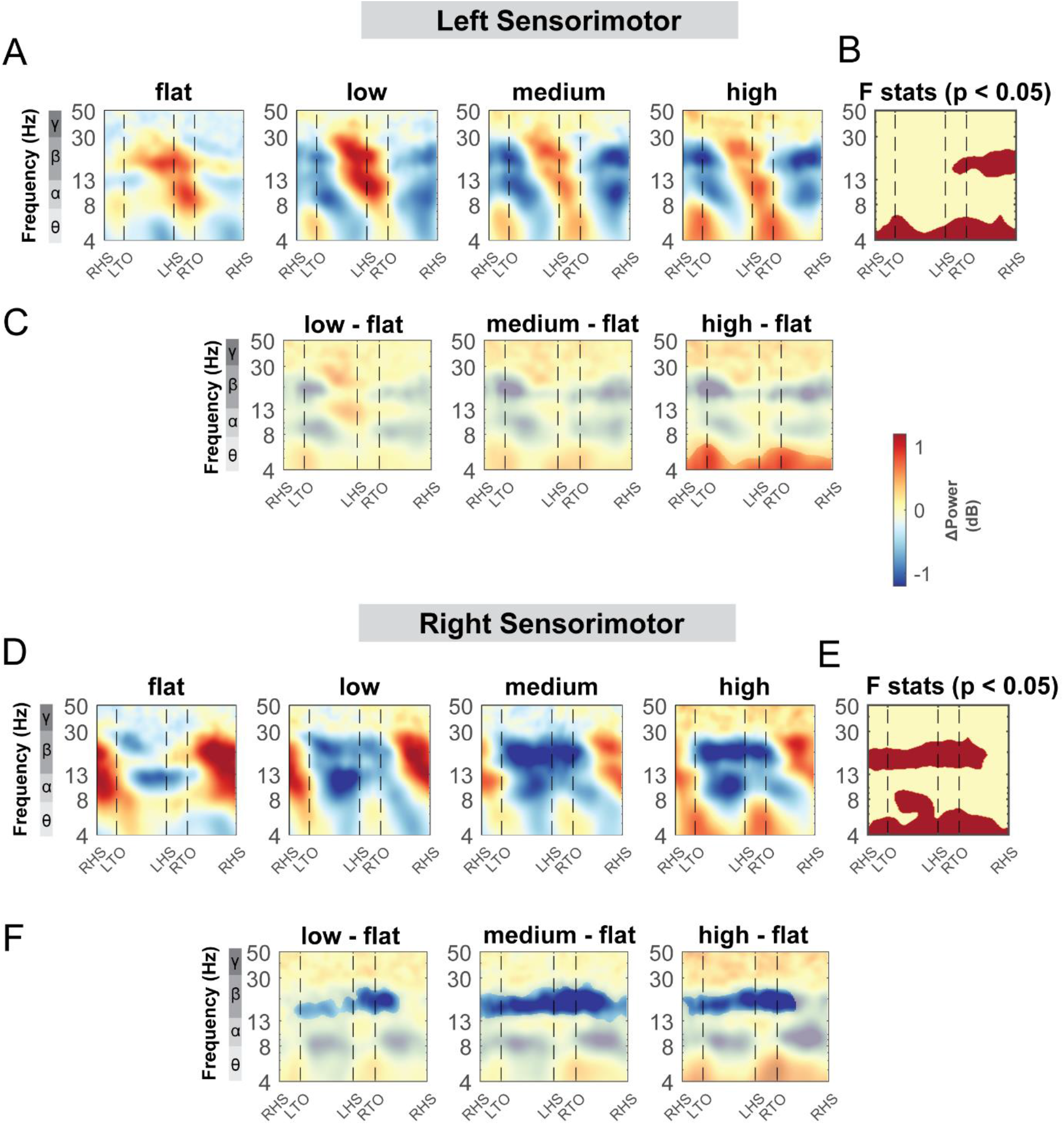
ERSPs at the left and right sensorimotor area with respect to the grand average of all conditions and with respect to the flat terrain condition. Averaged ERSP at different terrain at the left (A) and right sensorimotor cluster (D). Red indicated spectral power increased (neural synchronization) and blue indicated spectral power decreased (neural desynchronization) relative to the grand average of all conditions. Vertical dashed lines indicated gait events. RHS: right heel strike; LTO: left toe off; LHS: left heel strike; RTO: right toe off. (B, E) Significant effect of terrain on ERSPs across gait cycle with non-parametric statistics, with red indicating significance (p<0.05). ERSPs with respect to flat condition at the left (C) and right (F) sensorimotor cluster. Regions that are not significantly different from flat condition have a semi- transparent mask.

#### Sensorimotor clusters

Theta, alpha, and beta power changed with terrain unevenness during distinct gait phases for both the left and right sensorimotor clusters (Fig. 10). We found a significant effect of terrain in theta band across the gait cycle for both left and right clusters (Fig. 10B, D), with greater theta power in the high terrain condition especially during the double support phase as compared to that in the flat condition. At the left sensorimotor cluster, there was also an effect of terrain on beta power during the contralateral single-support stance phase (Fig. 10B) while we did not find any significant differences in beta power using pairwise comparison (Fig. 10C).

At the right sensorimotor cluster, we found a significant effect of terrain on the beta band during the double support phases and during the contralateral leg swing phase, with lower beta power in the low, medium, and high condition as compared to the flat condition (Fig. 10D - F). Lower beta power was observed following the contralateral foot strike to the ipsilateral swing in the low terrain condition as compared to the flat condition (Fig. 10F). Lower beta power was also observed during double support phases and the contralateral swing phase in the medium and high conditions. In addition, there was an effect of terrain on alpha band power from the contralateral mid-swing to the subsequent foot strike (Fig. 10E) although we did not find any significant pairwise difference with the flat terrain condition (Fig. 10F).

#### Posterior parietal clusters

Terrain modulated theta, alpha, and beta power across the gait cycle for both the left and right posterior parietal clusters (Fig. 11). There was a significant effect of terrain in theta band across the gait cycle for the left cluster (Fig. 11B), with greater theta power associated with greater terrain unevenness, especially at the double support phase. There was also an effect of terrain on alpha and beta power across the gait cycle at both left and right clusters, with lower alpha and beta power associated with greater terrain unevenness (Fig. 11 B, E). Strong desynchronization at alpha and beta band was observed in both left and right clusters during all levels of uneven terrain walking as compared to that during flat terrain (Fig. 11C, F).

**Fig. 11:**
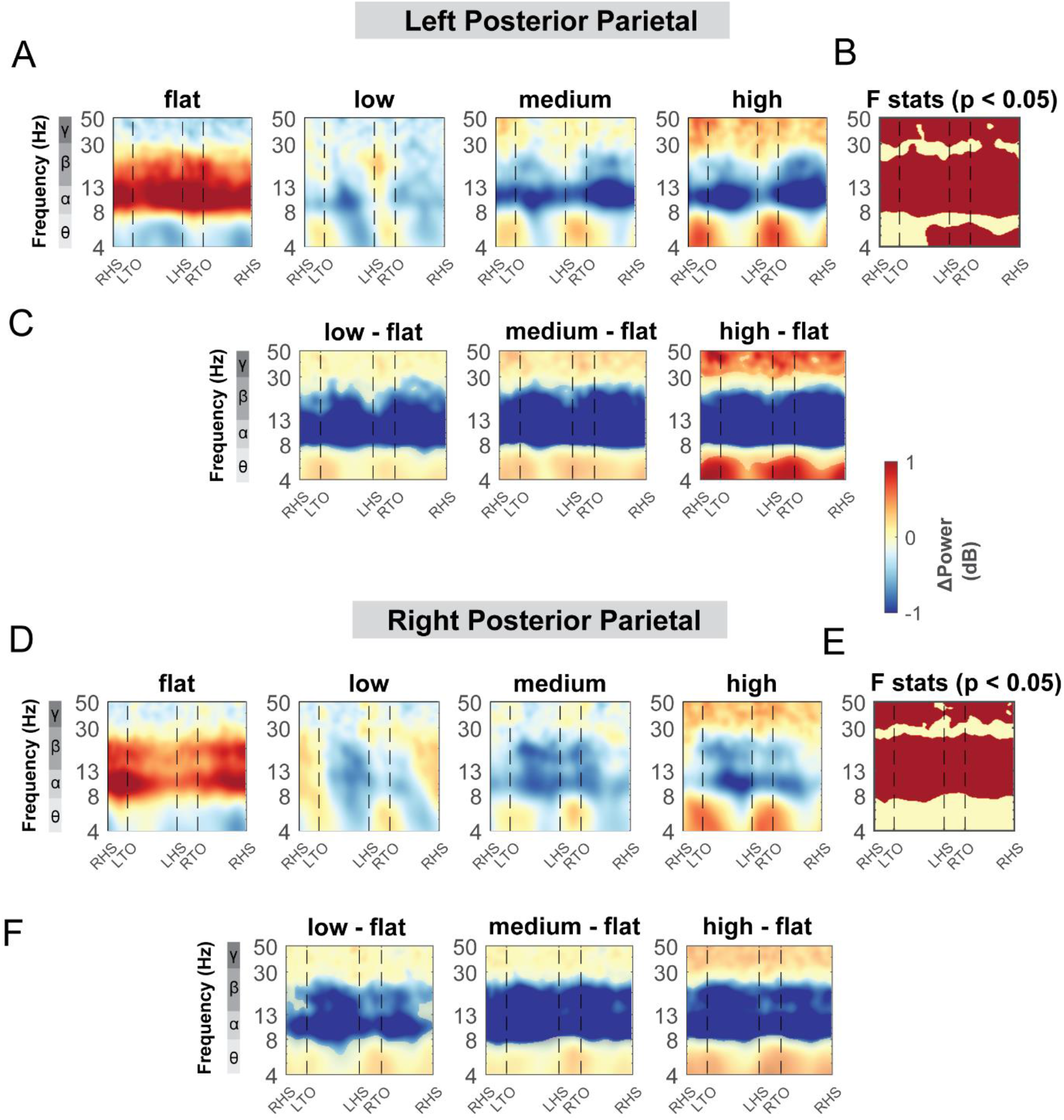
ERSPs at the left and right posterior parietal area with respect to the grand average of all conditions and with respect to the flat terrain condition. Averaged ERSP at different terrain at the left (**A**) and right posterior parietal cluster (**D**). Red indicated spectral power increased (neural synchronization) and blue indicated spectral power decreased (neural desynchronization) relative to the grand average of all conditions. Vertical dashed lines indicated gait events. RHS: right heel strike; LTO: left toe off; LHS: left heel strike; RTO: right toe off. (**B, E**) Significant effect of terrain on ERSPs across gait cycle with non-parametric statistics, with red indicating significance (p<0.05). ERSPs with respect to flat condition at the left (**C**) and right (**F**) posterior parietal cluster. Regions that are not significantly different from flat condition have a semi-transparent mask as determined by cluster-based permutation.

#### Mid/posterior cingulate cluster

The mid/posterior cingulate cluster also demonstrated changes in theta, alpha, and beta band power during uneven terrain walking (Fig. 12). There was an effect of terrain on theta power across the gait cycle, with greater theta power associated with greater terrain unevenness particularly at the double support phase (Fig. 12B). There was also an effect of terrain on beta power only during the left swing phase. We found lower beta power in the medium terrain condition compared to the flat terrain (Fig. 12C).

**Fig. 12:**
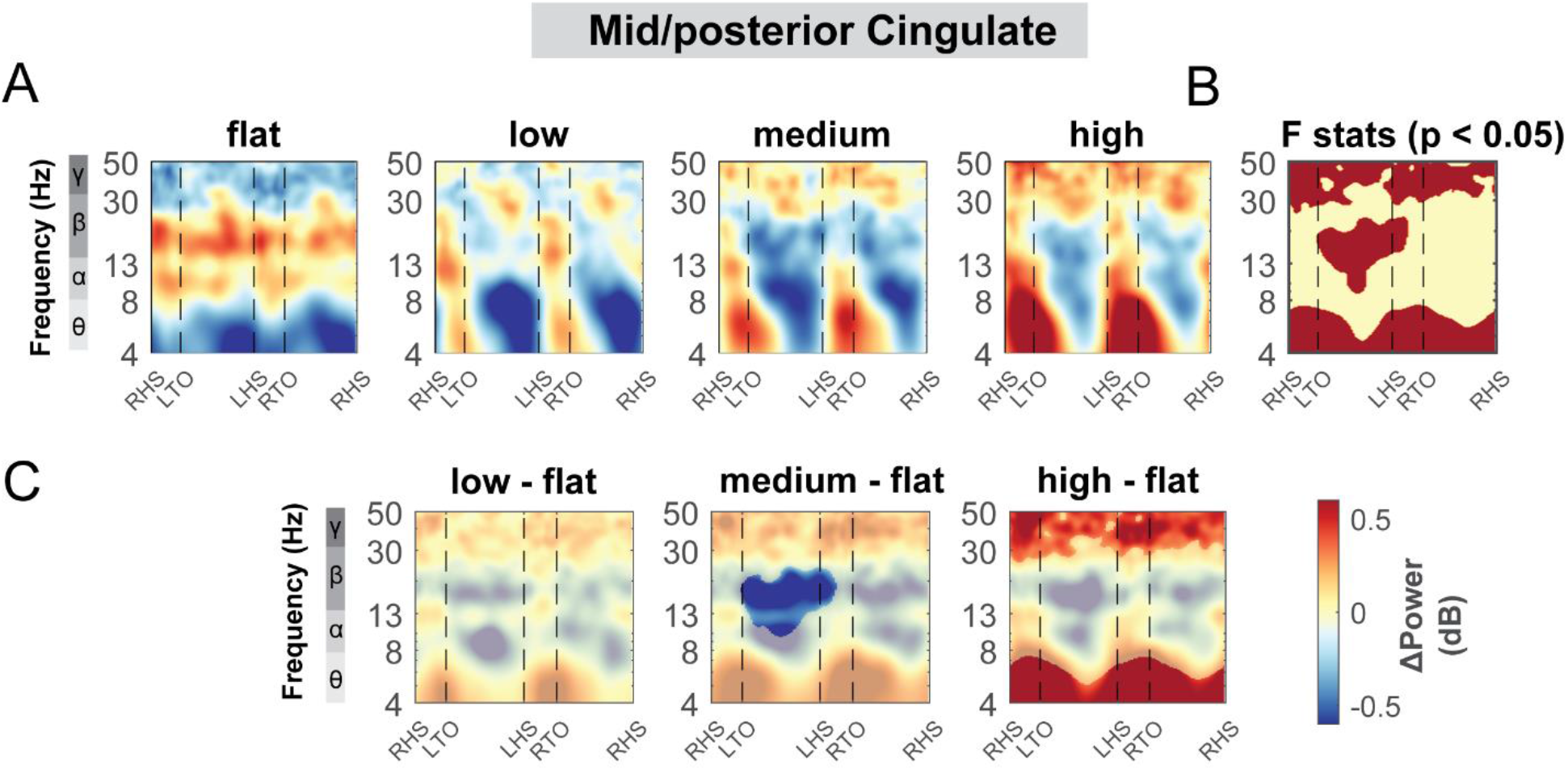
ERSPs at the mid/posterior cingulate area with respect to the grand average of all conditions and with respect to flat terrain condition. (**A**) Averaged ERSP at different terrain in the mid/posterior cingulate cluster. Red indicated spectral power increases (neural synchronization) and blue indicated spectral power decreases (neural desynchronization) relative to the grand average of all conditions. Vertical dashed lines indicated gait events. RHS: right heel strike; LTO: left toe off; LHS: left heel strike; RTO: right toe off. (**B**) Significant effect of terrain on ERSPs across gait cycle with non-parametric statistics, with red indicating significance (p<0.05). (**C**) ERSPs with respect to flat condition at the mid/posterior cingulate cluster. Regions that are not significantly different from flat condition have a semi-transparent mask as determined by cluster-based permutation.

#### Premotor, pre-supplementary motor, occipital, and caudate clusters

The left premotor, right pre-supplementary motor, occipital, and caudate clusters all showed power spectral fluctuation modulation by terrain (S2 - 5 Fig). There was an effect of terrain on theta and alpha power at the left premotor clusters (S2B Fig). Compared to the flat terrain, we found greater theta power in low, medium, and high terrain during the double support phase and the contralateral single-support stance phase at the left premotor cluster (S2C Fig). We also found greater alpha power in medium and high terrain conditions mainly during the double support phase (S2C Fig). At the right premotor cluster, there was an effect of terrain on theta power during the double support phase, with greater theta power observed in the high terrain condition compared to the flat terrain (S3B, C Fig). At the occipital cluster, we also found an effect of terrain on theta band power and gamma band power, with both greater theta power and gamma power in at low, medium, and high terrain compared to flat condition (S4 Fig).

Additionally, at the caudate cluster, we found an effect of terrain at theta, alpha, and gamma power (S5 Fig). Greater theta and alpha power were observed at double support phase in low and medium terrain condition and throughout the entire gait cycle at high terrain compared to the flat terrain.

**S2 Fig:**
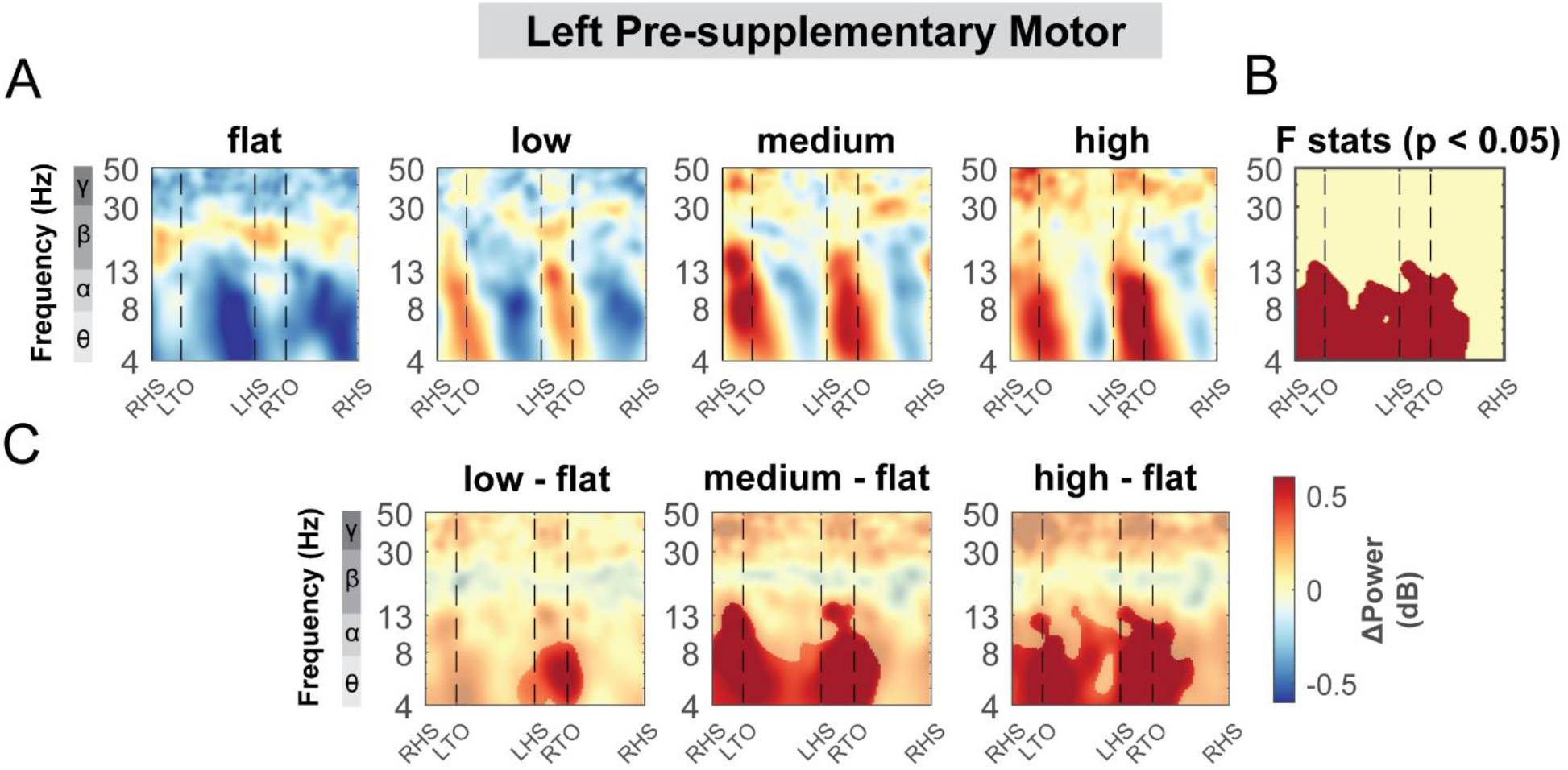
ERSPs at the left pre-supplementary motor clusters with respect to the grand average of all conditions and with respect to flat terrain condition. (**A**) Averaged ERSP at different terrain at the left pre-supplementary motor cluster. Red indicated spectral power increases (neural synchronization) and blue indicated spectral power decreases (neural desynchronization) relative to the grand average of all conditions. Vertical dashed lines indicated gait events. RHS: right heel strike; LTO: left toe off; LHS: left heel strike; RTO: right toe off. (**B**) Significant effect of terrain on ERSPs across gait cycle with non-parametric statistics, with red indicating significance (p<0.05). (**C**) ERSPs with respect to flat condition at the left pre-supplementary motor cluster. Regions that are not significantly different from flat condition have a semi-transparent mask as determined by cluster-based permutation.

**S3 Fig:**
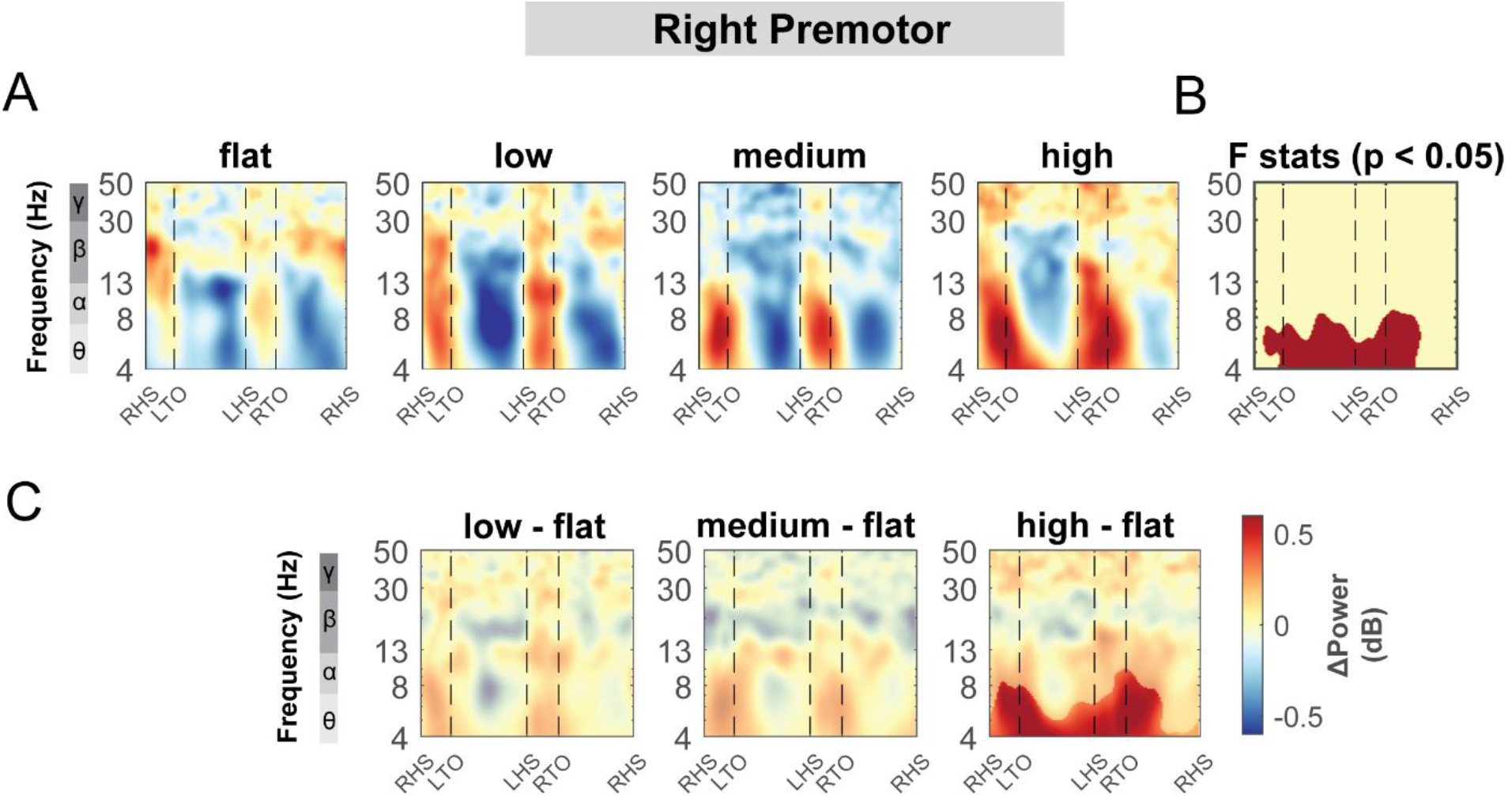
ERSPs at the right premotor cluster with respect to the grand average of all conditions and with respect to flat terrain condition. (**A**) Averaged ERSP at different terrain at the right premotor cluster. Red indicated spectral power increases (neural synchronization) and blue indicated spectral power decreases (neural desynchronization) relative to the grand average of all conditions. Vertical dashed lines indicated gait events. RHS: right heel strike; LTO: left toe off; LHS: left heel strike; RTO: right toe off. (**B**) Significant effect of terrain on ERSPs across gait cycle with non-parametric statistics, with red indicating significance (p<0.05). (**C**) ERSPs with respect to flat condition at the right premotor cluster. Regions that are not significantly different from flat condition have a semi-transparent mask as determined by cluster-based permutation.

**S4 Fig:**
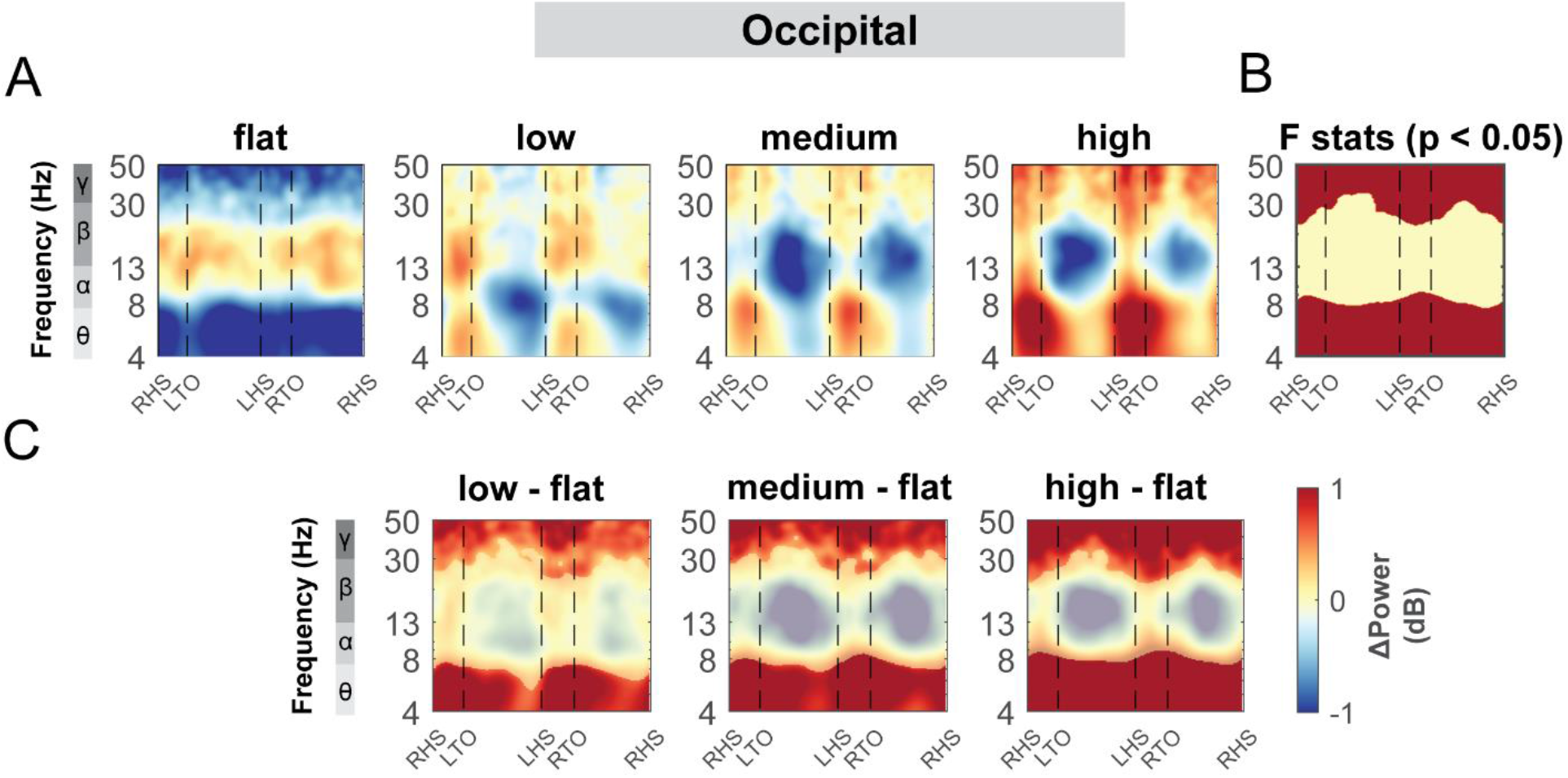
ERSPs at the occipital cluster with respect to the grand average of all conditions and with respect to flat terrain condition. (**A**) Averaged ERSP at different terrain at the occipital cluster. Red indicated spectral power increases (neural synchronization) and blue indicated spectral power decreases (neural desynchronization) relative to the grand average of all conditions. Vertical dashed lines indicated gait events. RHS: right heel strike; LTO: left toe off; LHS: left heel strike; RTO: right toe off. (**B**) Significant effect of terrain on ERSPs across gait cycle with non-parametric statistics, with red indicating significance (p < 0.05). (**C**) ERSPs with respect to flat condition at the occipital cluster. Regions that are not significantly different from flat condition have a semi-transparent mask as determined by cluster-based permutation.

**S5 Fig:**
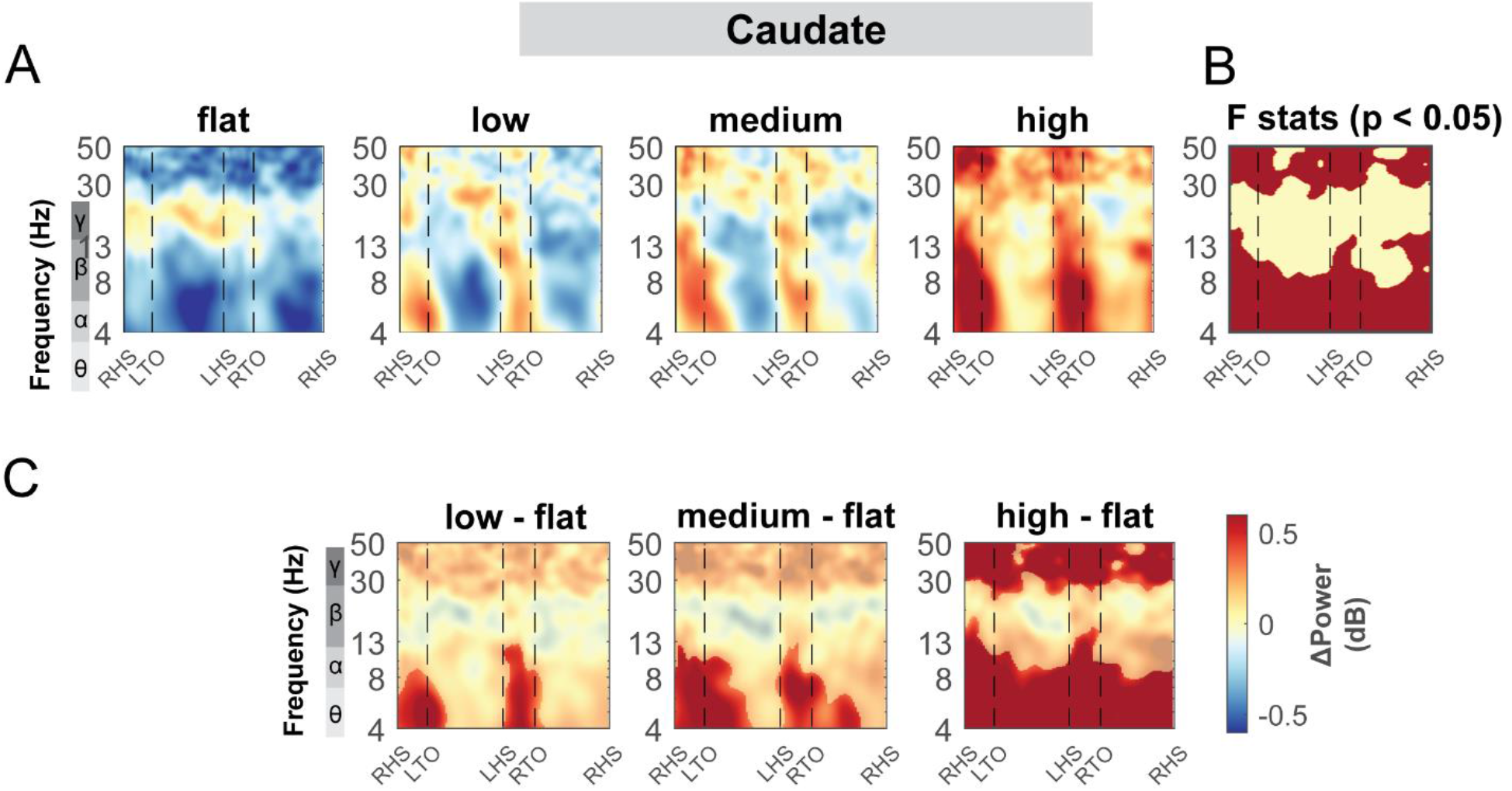
ERSPs at caudate cluster with respect to the grand average of all conditions and with respect to flat terrain condition. (**A**) Averaged ERSP at different terrain at the caudate cluster. Red indicated spectral power increases (neural synchronization) and blue indicated spectral power decreases (neural desynchronization) relative to the grand average of all conditions. Vertical dashed lines indicated gait events. RHS: right heel strike; LTO: left toe off; LHS: left heel strike; RTO: right toe off. (**B**) Significant effect of terrain on ERSPs across gait cycle with non-parametric statistics, with red indicating significance (p<0.05). (**C**) ERSPs with respect to flat condition at the caudate cluster. Regions that are not significantly different from flat condition have a semi-transparent mask as determined by cluster-based permutation.

### Speed modulation of EEG power spectral density

For the secondary objective of this study, we employed a linear mixed effect model for each outcome measure (average power for each band for each cluster) to assess the effect of walking speed on EEG power spectral at multiple cortical areas (Fig. 13 - 15). There was a main effect of speed on average theta power (F(1, 110) = 4.2, p = 0.04, Cohen’s f^2^ = 0.055) at the left sensorimotor area, with greater theta power associated with faster walking speed (Fig. 13E).

**Fig. 13:**
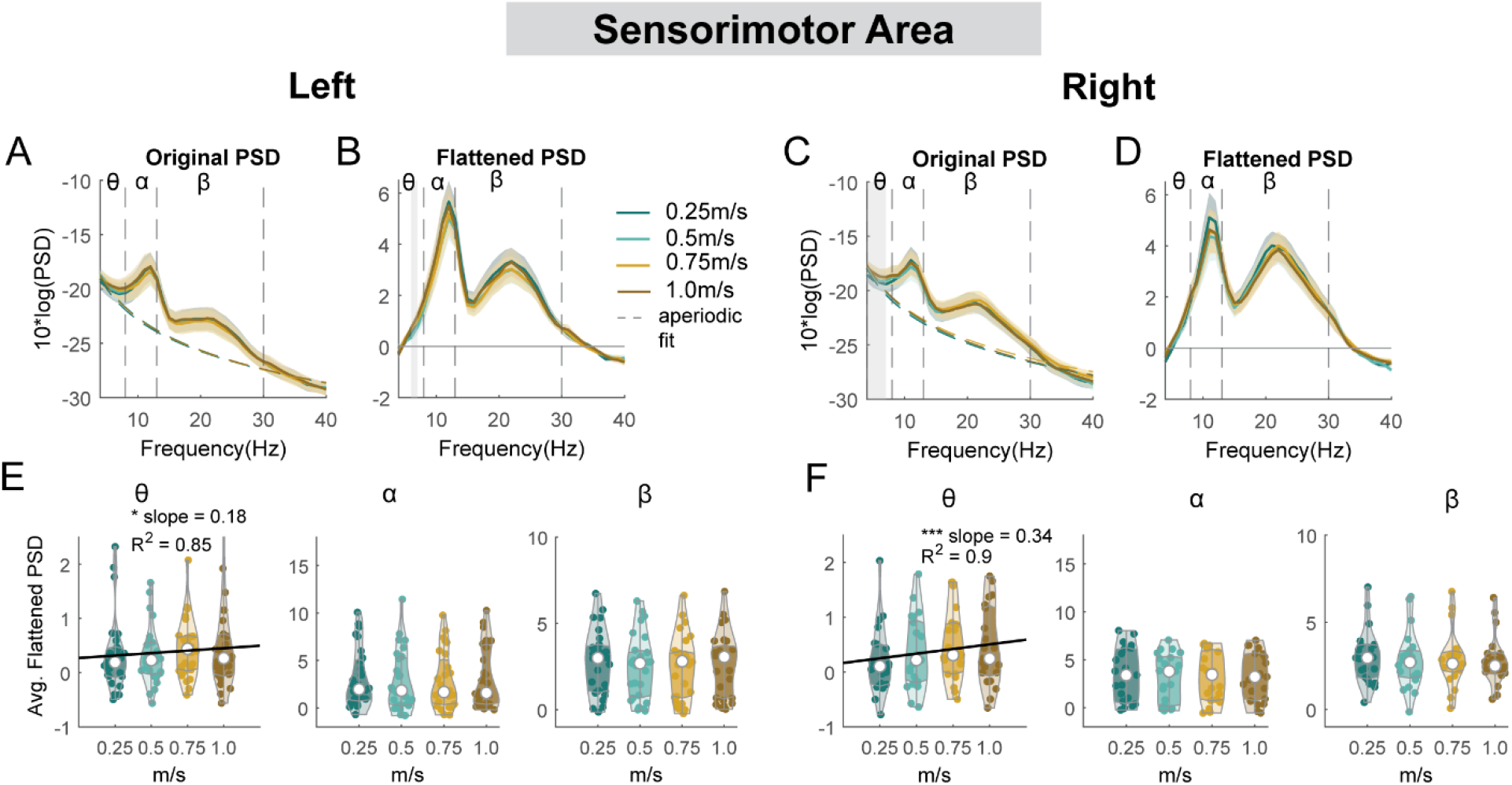
Power spectral density changes with walking speed at the left and right sensorimotor area. Average original PSDs changed with walking speed for the left (**A**) and right (**C**) sensorimotor cluster. Shaded colored areas indicated standard error of PSDs across components in the cluster. Dashed colored lines indicated average aperiodic fit. Gray shaded areas indicated a significant effect of terrain on PSDs. Vertical black dashed lines indicated main frequency bands of interest – theta (4 - 8 Hz), alpha (8 - 13 Hz), and beta (13 - 30 Hz). Average flattened PSDs after removing the aperiodic fit for the left (**B**) and right (**D**) sensorimotor cluster. (**E - F**) Average power for theta, alpha, and beta band computed from flattened PSDs for all components within the cluster for the left and right clusters. Black line indicates a significant correlation between average power and walking speed (***p < 0.001).

However, we did not find a significant main effect of speed on alpha power (F(1,110) = 0.5, p = 0.5, Cohen’s f^2^ = 0.006) or beta power (F(1, 110) = 1.6, p = 0.2, Cohen’s f^2^ = 0.0192).

Additionally, we observed a main effect of speed on average theta power at the right sensorimotor cluster (F(1, 82) = 14.9, p < 0.001, Cohen’s f^2^ = 0.26), with greater theta power at faster walking speed. There was no effect of speed on alpha power (F(1, 82) = 0.39, p = 0.5, Cohen’s f^2^ = 0.0064) or beta power (F(1, 82) = 2.1, p = 0.15, Cohen’s f^2^ = 0.035).

For the left posterior parietal cluster, there was a main effect of speed on average theta power (F(1, 106) = 5.3, p = 0.023, Cohen’s f^2^ = 0.076), with greater theta power associated with faster walking speed (Fig. 14C). However, we did not find a significant main effect of speed on alpha power (F(1, 106) = 0.04, p = 0.83, Cohen’s f^2^ = 0) or beta power (F(1, 106) = 0.22, p = 0.64, Cohen’s f^2^ = 0.003). For the right posterior parietal cluster, we found a main effect of speed on average theta power (F(1,110) = 4.2, p = 0.04, Cohen’s f^2^ = 0.055), with greater theta power associated with faster walking speed. We also found a main effect of speed on average beta power (F(1, 110) = 8.4, p = 0.005, Cohen’s f^2^ = 0.1) with lower beta power associated with faster walking speed. There was no effect of speed on alpha power (F(1, 110) = 3.85, p = 0.052, Cohen’s f^2^ = 0.048).

**Fig. 14:**
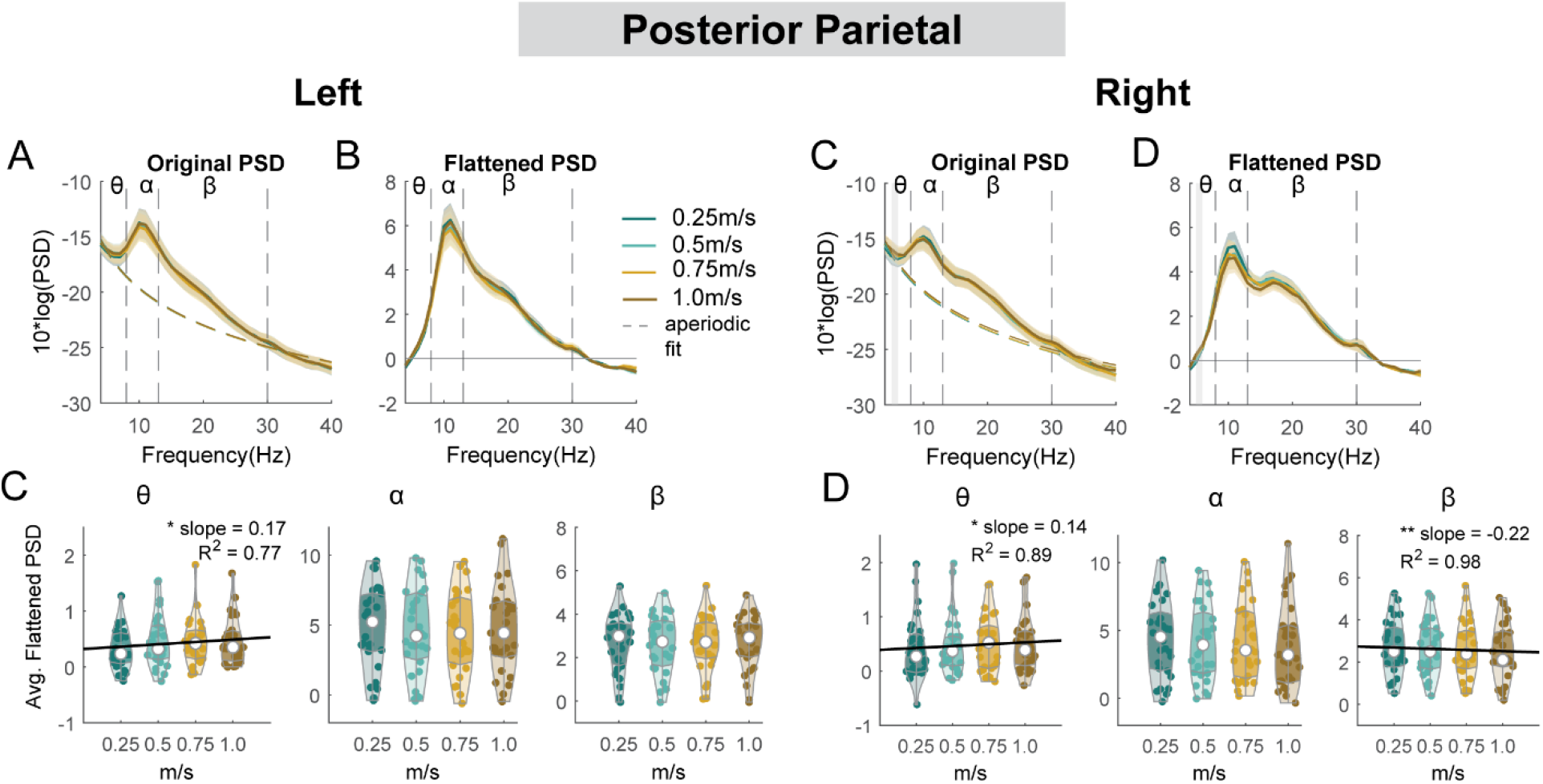
Power spectral density changes with walking speed at the left and right posterior parietal area. Average original PSDs changed with walking speed for the left (**A**) and right (**C**) posterior parietal cluster. Shaded colored areas indicated standard error of PSDs across components in the cluster. Dashed colored lines indicated average aperiodic fit. Gray shaded areas indicated a significant effect of terrain on PSDs. Vertical black dashed lines indicated main frequency bands of interest – theta (4 - 8 Hz), alpha (8 - 13 Hz), and beta (13 - 30 Hz). Average flattened PSDs after removing the aperiodic fit for the left (**B**) and right (**D**) posterior parietal cluster. (**E - F**) Average power for theta, alpha, and beta band computed from flattened PSDs for all components within the cluster for the left and right clusters. Black line indicates a significant correlation between average power and walking speed (*p < 0.05, **p < 0.01, ***p < 0.001).

There was a main effect of speed on the average beta power (F(1, 94) = 5.4, p = 0.023, Cohen’s f^2^ = 0.047) at the mid/posterior cingulate, with lower beta power associated with faster walking speed (Fig. 15). However, we did not find a significant main effect of speed on theta power (F(1, 94) = 3.2, p = 0.08, Cohen’s f^2^ = 0.01) or alpha power (F(1, 94) = 0.65, p = 0.42, Cohen’s f^2^ = 0.076).

**Fig. 15:**
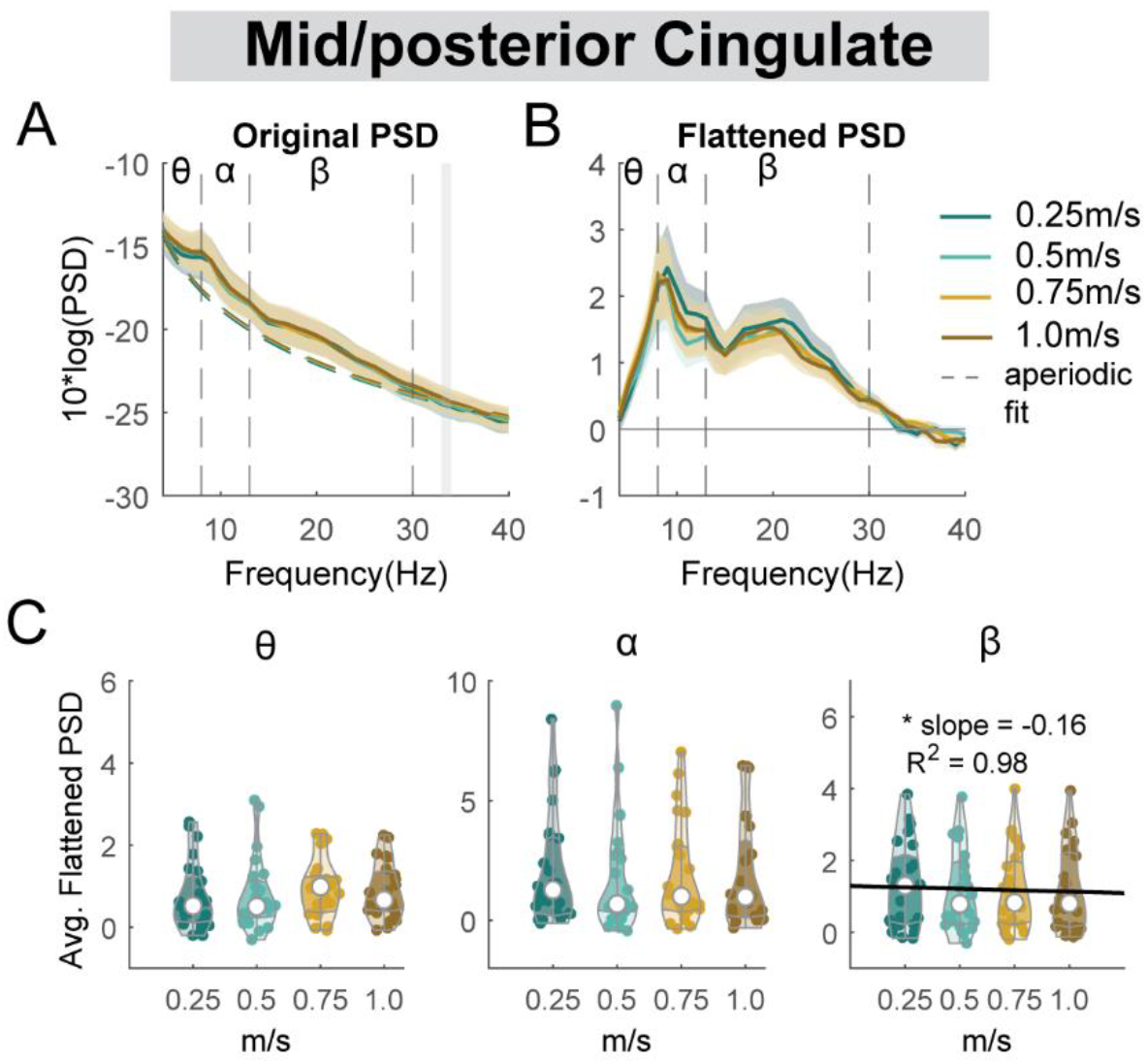
Power spectral density changes with walking speed at the mid/posterior cingulate area. (**A**) Average original PSDs changed with walking speed for the mid/posterior cingulate cluster. Shaded colored areas indicated standard error of PSDs across components in the cluster. Dashed colored lines indicated average aperiodic fit. Gray shaded areas indicated a significant effect of terrain on PSDs. Vertical black dashed lines indicated main frequency bands of interest – theta (4 - 8 Hz), alpha (8 - 13 Hz), and beta (13 - 30 Hz). (**B**) Average flattened PSDs after removing the aperiodic fit for the mid/posterior cingulate cluster. (**C**) Average power for theta, alpha, and beta band computed from flattened PSDs for all components within the cluster. Black line indicates a significant correlation between average power and walking speed (*p < 0.05, **p < 0.01, ***p < 0.001).

Similarly, we performed an exploratory analysis on the effect of walking speed on band powers at the left pre-supplementary motor, right premotor, occipital, and caudate clusters (Fig. 16).

**Fig. 16:**
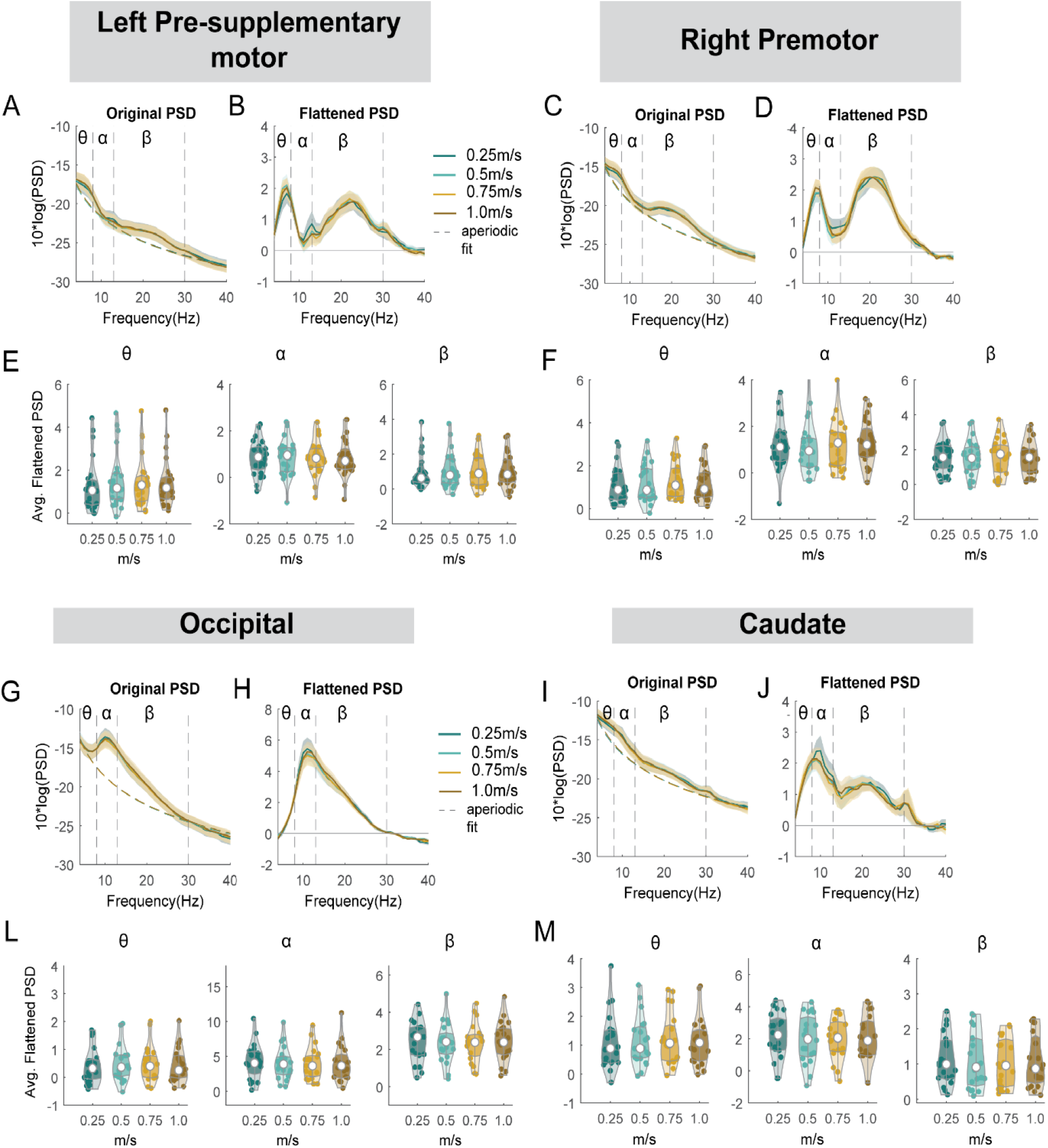
Power spectral density changes with walking speed at the left pre-supplementary motor, right premotor, occipital, and caudate area. (**A**) Average original PSDs changed with walking speed for the left pre-supplementary cluster. Shaded colored areas indicated standard error of PSDs across components in the cluster. Dashed colored lines indicated average aperiodic fit. Gray shaded areas indicated a significant effect of speed on PSDs. Vertical black dashed lines indicated main frequency bands of interest – theta (4 - 8 Hz), alpha (8 - 13 Hz), and beta (13 - 30 Hz). (**B**) Average flattened PSDs after removing the aperiodic fit for the left pre-supplementary motor cluster. (**E**) Average power for theta, alpha, and beta band computed from flattened PSDs for all components within the cluster for the left pre-supplementary clusters. Black line indicates a significant correlation between average power and walking speed (*p < 0.05, **p < 0.01, ***p < 0.001). (**C**) Average original PSDs changed with walking speed for the right premotor cluster. (**D**) Average flattened PSDs after removing the aperiodic fit for the right premotor cluster. (**F**) Average power for theta, alpha, and beta band computed from flattened PSDs for all components within the right premotor cluster. (**G**) Average original PSDs changed with walking speed for the occipital cluster. (**H**) Average flattened PSDs after removing the aperiodic fit for the occipital cluster. (**L**) Average power for theta, alpha, and beta band computed from flattened PSDs for all components within the occipital cluster. (**I**) Average original PSDs changed with walking speed for the caudate cluster. (**J**) Average flattened PSDs after removing the aperiodic fit for the caudate cluster. (**M**) Average power for theta, alpha, and beta band computed from flattened PSDs for all components within the caudate cluster.

There was no effect of speed on theta, alpha, or beta power at either left pre-supplementary motor, right premotor cluster, occipital, or caudate cluster (all p > 0.05).

### Gait-related spectral perturbations during walking at different speeds

We again computed the event-related power perturbations within the gait cycle at each walking speed with respect to the average power across the gait cycle of the same condition at each frequency [35]. We unmasked the significant deviations from the average spectrum of each condition with a bootstrap method with false discovery rate multiple comparison correction.

Similar to Fig. 9, alpha band and beta band activity showed lateralization for left and right sensorimotor clusters at different walking speeds (Fig. 17A, B). We observed alpha and beta desynchronization during the contralateral swing phase and synchronization during the ipsilateral swing phase at 0.25 m/s and 0.5 m/s. At higher walking speeds (0.75 m/s and 1.0 m/s), we observed additional alpha and beta synchronization during the double support phase from the ipsilateral foot strike until the contralateral foot off. Descriptively, power fluctuations decreased with walking speed at the alpha and beta band.

**Fig. 17.**
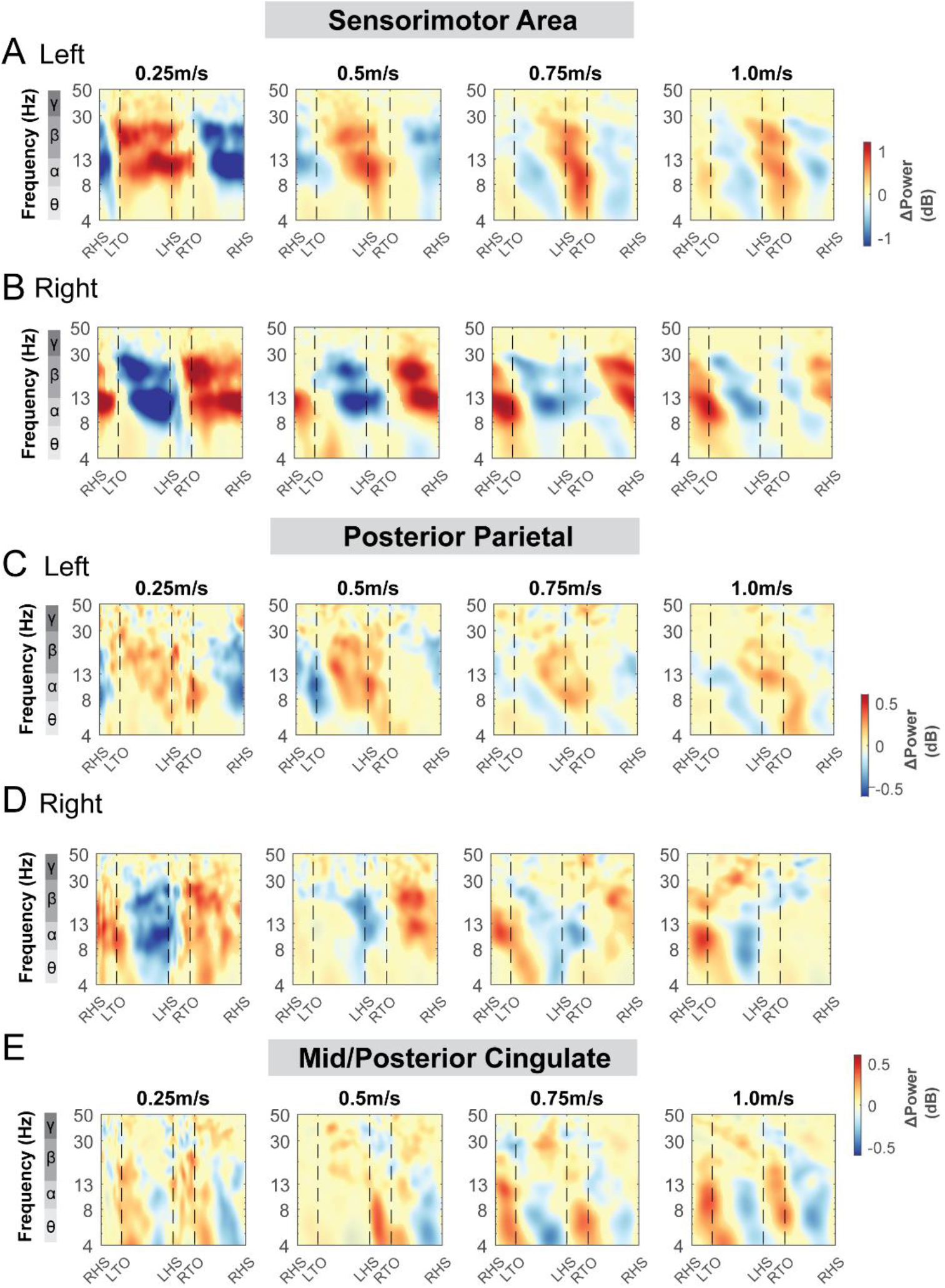
ERSPs at the sensorimotor (A-B), posterior parietal (C-D), and mid/posterior cingulate areas (E) with respect to the average of each condition at different speeds. The x- axes of the ERSPs are time in gait cycle (RHS: right heel strike; LTO: left toe off; LHS: left heel strike; RTO: right toe off). All unmasked colors are statistically significant spectral power fluctuations relative to the mean power within the same condition. Colors indicate significant increases (red, synchronization) and decreases (blue, desynchronization) in spectral power from the average spectrum for all gait cycles to visualize intra-stride changes in the spectrograms. These data are significance masked (p<0.05) through nonparametric bootstrapping with multiple comparison correction using false discovery rate.

At the posterior parietal area, the spectral power fluctuations across the gait cycle were not consistent across the speeds (Fig. 17C, D). We observed alpha and beta desynchronization during the contralateral swing phase and synchronization during the contralateral swing phase at slower walking speeds (0.25 m/s and 0.5 m/s). At the mid/posterior cingulate cluster, we observed theta and alpha synchronization during both double support phase and desynchronization during swing phase (Fig. 17E). At the supplementary motor clusters, occipital cluster, and caudate clusters, power spectral fluctuation within each condition was not different across speeds (S5 Fig). Only at higher speeds (0.75 m/s and 1.0 m/s) did we observe prominent theta and alpha band synchronization during the double support phase and desynchronization at mid-swing phase at the supplementary motor, occipital, and caudate clusters.

**S6 Fig.**
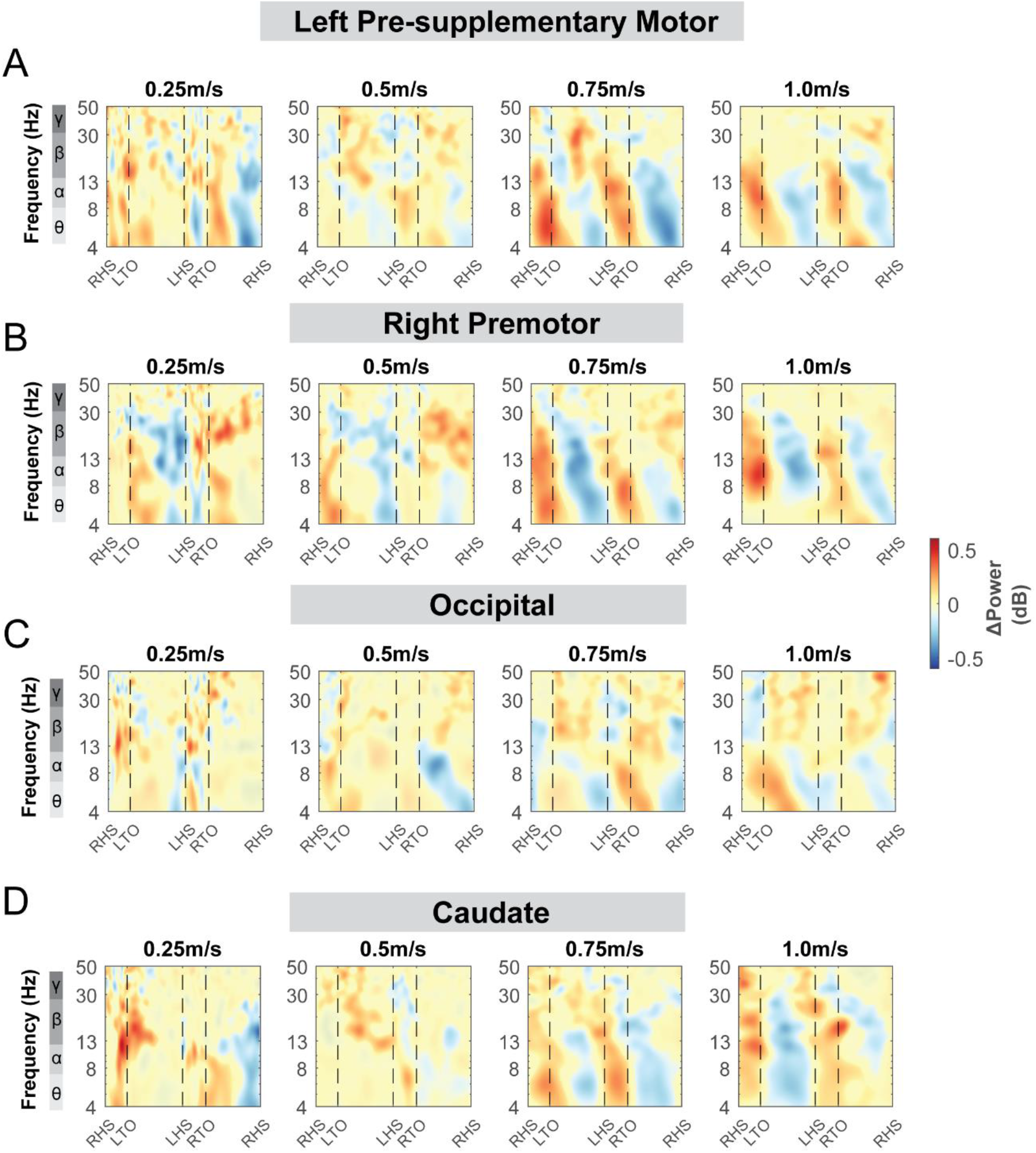
ERSPs at the left pre-supplementary motor (A), right premotor (B), occipital (C), and caudate areas (D) with respect to the average of each condition at different speeds. The x-axes of the ERSPs are time in gait cycle (RHS: right heel strike; LTO: left toe off; LHS: left heel strike; RTO: right toe off). All unmasked colors are statistically significant spectral power fluctuations relative to the mean power within the same condition. Colors indicate significant increases (red, synchronization) and decreases (blue, desynchronization) in spectral power from the average spectrum for all gait cycles to visualize intra-stride changes in the spectrograms. These data are significance masked (p<0.05) through nonparametric bootstrapping with multiple comparison correction using false discovery rate.

### Effects of walking speed on event-related power perturbations

We then computed the ERSPs with respect to the grand average of all conditions to assess the effect of walking speed on spectral power fluctuation tied to gait events. All clusters showed spectral power fluctuation in event-related spectral perturbation plots at various frequency bands during the gait cycle with red indicated synchronization and blue indicated desynchronization (for example, Fig. 18A). We used non-parametric permutation statistics with cluster-based multiple comparison correction to determine the time-frequency differences across speed conditions with red color indicated significant differences across speed conditions (p < 0.05; for example, Fig. 18B). To determine how spectral power changed relative to the 1.0 m/s condition, we computed the differences in ERSPs between each speed condition relative to the 1.0 m/s condition (ERSP_terrain_ – ERSP_1.0m/s_) (for example, Fig. 18C). Regions that were not significantly different from 1.0 m/s condition have a semi-transparent mask using permutation statistics with cluster-based multiple comparison correction (for example, Fig. 18C).

**Fig. 18:**
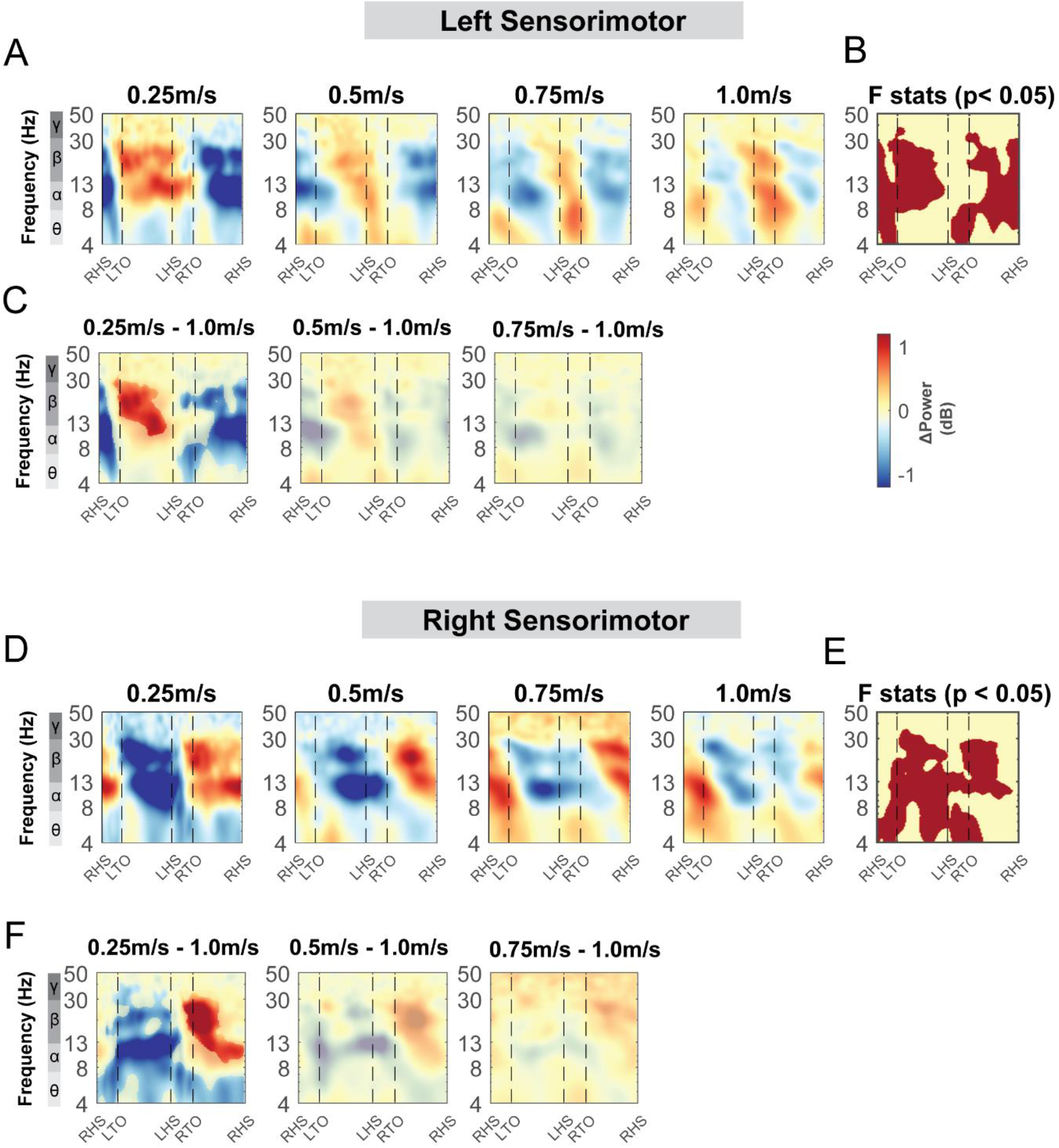
ERSPs at the sensorimotor area with respect to the grand average of all conditions and with respect to the 1.0 m/s speed condition. Averaged ERSP at different speeds at the left (**A**) and right sensorimotor cluster (**D**). Red indicated spectral power increases (neural synchronization) and blue indicated spectral power decreases (neural desynchronization) relative to the grand average of all conditions. Vertical dashed lines indicated gait events. RHS: right heel strike; LTO: left toe off; LHS: left heel strike; RTO: right toe off. (**B, E**) Significant effect of terrain on ERSPs across gait cycle with non-parametric statistics, with red indicating significance (p < 0.05). ERSPs with respect to 1.0m/s speed condition at the left (**C**) and right (**F**) sensorimotor cluster. Regions that are not significantly different from 1.0 m/s condition have a semi-transparent mask.

#### Sensorimotor clusters

Theta, alpha, and beta power fluctuations changed with walking speed for both the left and right sensorimotor clusters (Fig. 18A, D). We found a significant effect of speed in the theta band during double support phases for both left and right clusters (Fig. 18B, E). Theta power was lower during the double support phase when walking at 0.25 m/s compared to 1.0 m/s. There was also an effect of speed on theta power during the contralateral swing phase in both the left and right sensorimotor clusters. We observed a lower theta power when walking at 0.25 m/s compared to 1.0 m/s (Fig. 18C, F). Additionally, we observed an effect of speed on alpha power during the swing phase at both clusters. We found a lower alpha power at 0.25 m/s during the contralateral swing phase and a greater alpha power during the contralateral stance phase at 0.25 m/s compared to 1.0 m/s. Lastly, we found an effect of walking speed on the beta power during both swing phases (Fig. 18B, E). Beta power was greater during the contralateral swing phase and lower during the contralateral stance phase when walking at 0.25 m/s compared to 1.0 m/s.

#### Posterior parietal clusters, mid/posterior cingulate clusters, and other clusters

We did not find any effect of speed on ERSPs at the posterior parietal clusters, except for beta power during the contralateral swing phase at the right posterior parietal clusters (Fig. 19B, E). We found a lower beta power during the contralateral swing phase at the right posterior parietal clusters when walking at 0.25m/s versus 1.0m/s (Fig. 19E, F). We also did not find any effect of speed on ERSPs at the mid/posterior cingulate cluster (p > 0.05; Fig. 20).

**Fig. 19:**
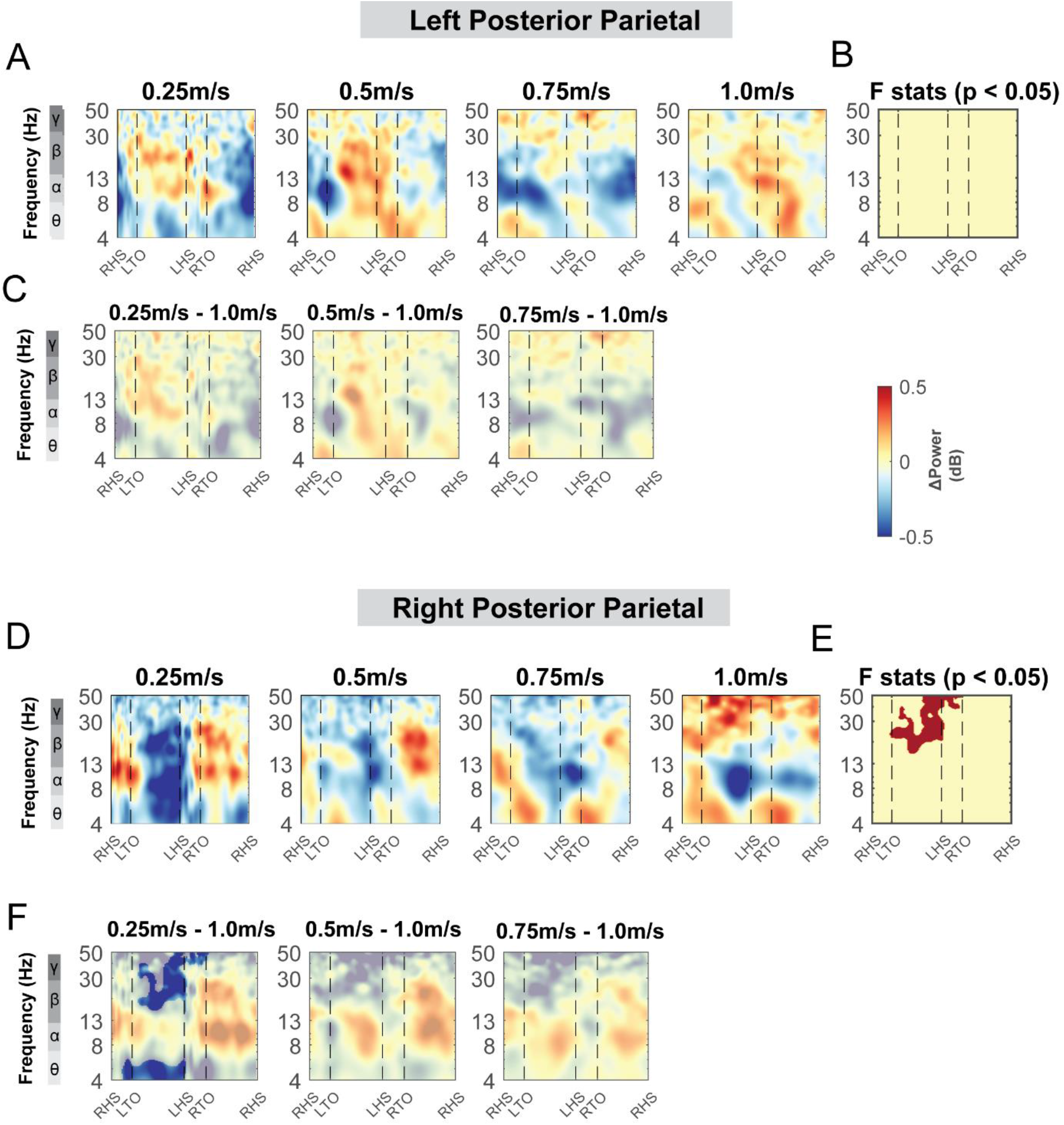
ERSPs at the posterior parietal area with respect to the grand average of all conditions and with respect to the 1.0 m/s speed condition. Averaged ERSP at different speeds at the left (**A**) and right posterior parietal cluster (**D**). Red indicated spectral power increases (neural synchronization) and blue indicated spectral power decreases (neural desynchronization) relative to the grand average of all conditions. Vertical dashed lines indicated gait events. RHS: right foot strike; LTO: left foot off; LHS: left foot strike; RTO: right foot off. (**B, E**) Significant effect of terrain on ERSPs across gait cycle with non-parametric statistics, with red indicating significance (p < 0.05). ERSPs with respect to 1.0m/s speed condition at the left (**C**) and right (**F**) posterior parietal cluster. Regions that are not significantly different from 1.0 m/s condition have a semi-transparent mask.

**Fig. 20:**
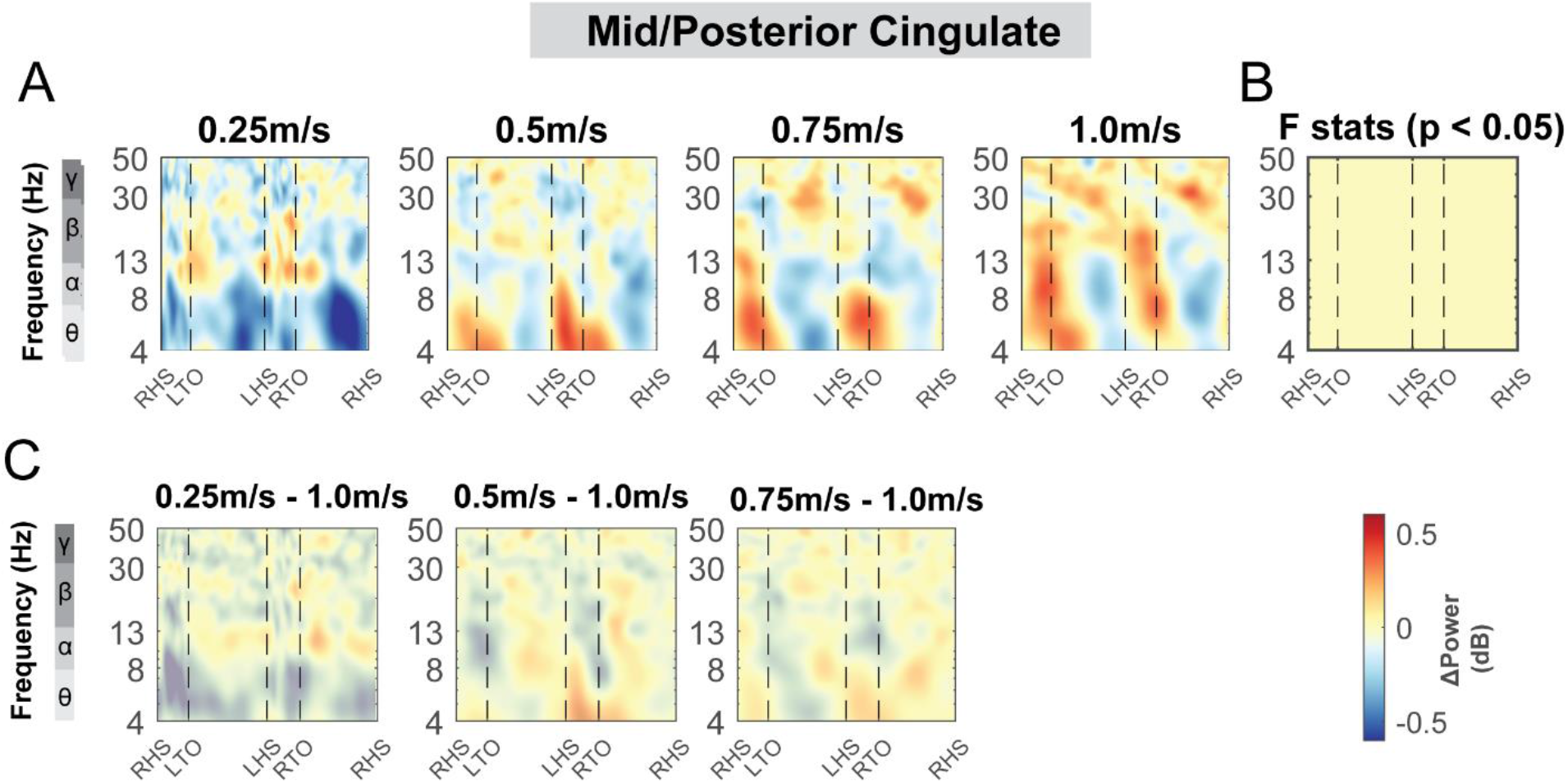
ERSPs at the mid/posterior cingulate area with respect to the grand average of all conditions and with respect to the 1.0 m/s speed condition. Averaged ERSP at different speeds at mid/posterior cingulate cluster (**A**). Red indicated spectral power increases (neural synchronization) and blue indicated spectral power decreases (neural desynchronization) relative to the grand average of all conditions. Vertical dashed lines indicated gait events. RHS: right heel strike; LTO: left toe off; LHS: left heel strike; RTO: right toe off. (**B**) Significant effect of terrain on ERSPs across gait cycle with non-parametric statistics, with red indicating significance (p<0.05). (**C**) ERSPs with respect to 1.0 m/s speed condition. Regions that are not significantly different from 1.0 m/s condition have a semi-transparent mask.

There was an effect of speed on ERSPs in theta band during the double support phase only for the left pre-supplementary motor cluster (p < 0.05) while we did not find any differences in ERSPs using pairwise comparison referenced to the 1.0 m/s condition. For other clusters including the right premotor, occipital, and caudate clusters, we did not find any significant effect of speed on ERSPs (all p’s >0.05) (S6-8 Fig).

**S7 Fig:**
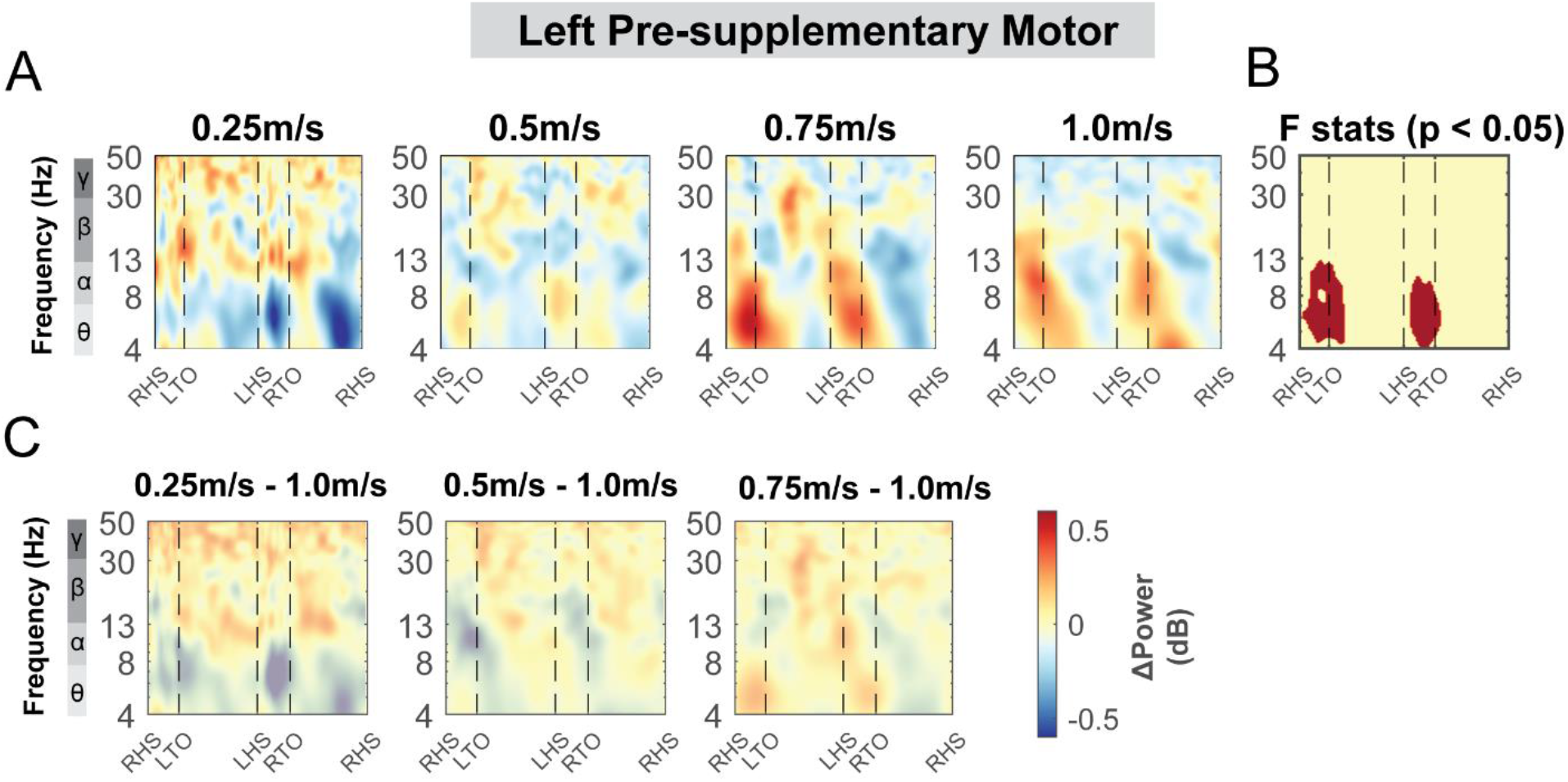
ERSPs at the left pre-supplementary motor area with respect to the grand average of all conditions and with respect to the 1.0 m/s speed condition. (**A**) Averaged ERSP at different speeds at the left pre-supplementary cluster. Red indicated spectral power increases (neural synchronization) and blue indicated spectral power decreases (neural desynchronization) relative to the grand average of all conditions. Vertical dashed lines indicated gait events. RHS: right heel strike; LTO: left toe off; LHS: left heel strike; RTO: right toe off. (**B**) Significant effect of terrain on ERSPs across gait cycle with non-parametric statistics, with red indicating significance (p<0.05). (**C**) ERSPs with respect to 1.0 m/s speed condition. Regions that are not significantly different from 1.0 m/s condition have a semi-transparent mask.

**S8 Fig:**
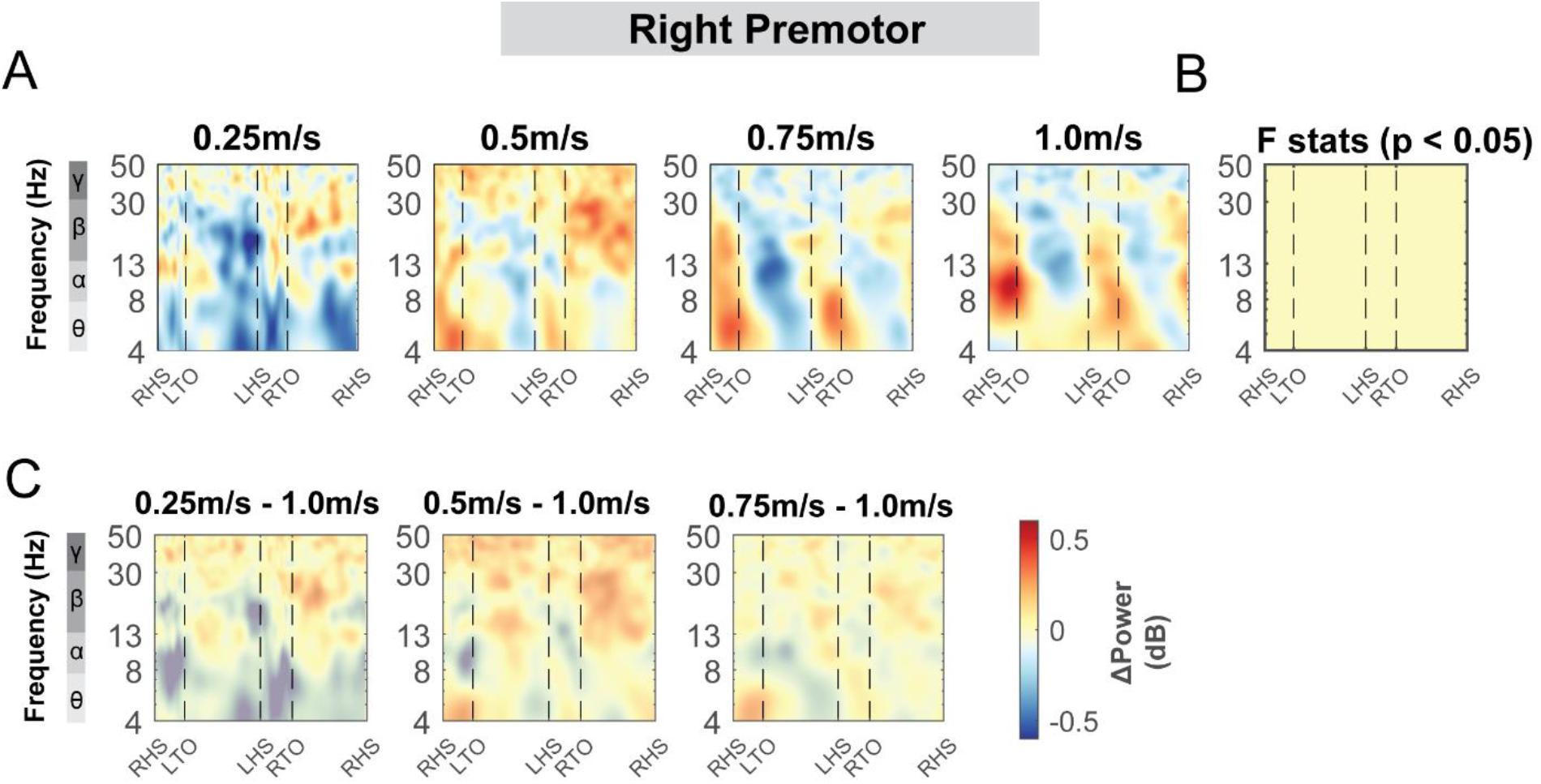
ERSPs at the right premotor cluster with respect to the grand average of all conditions and with respect to the 1.0m/s speed condition. (**A**) Averaged ERSP at different speeds at the right premotor cluster. Red indicated spectral power increases (neural synchronization) and blue indicated spectral power decreases (neural desynchronization) relative to the grand average of all conditions. Vertical dashed lines indicated gait events. RHS: right heel strike; LTO: left toe off; LHS: left heel strike; RTO: right toe off. (**B**) Significant effect of terrain on ERSPs across gait cycle with non-parametric statistics, with red indicating significance (p<0.05). (**C**) ERSPs with respect to 1.0 m/s speed condition. Regions that are not significantly different from 1.0 m/s condition have a semi-transparent mask.

**S9 Fig:**
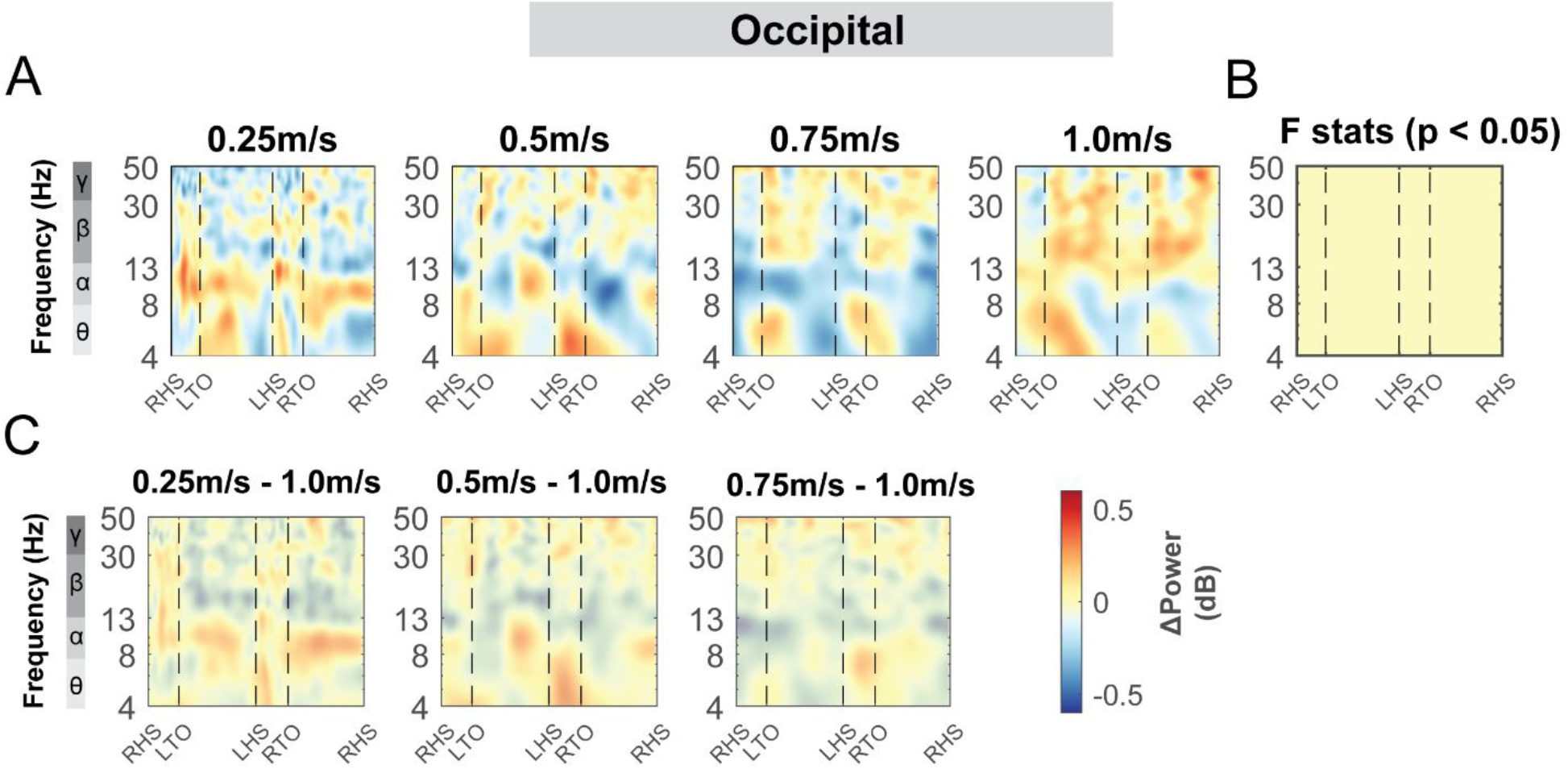
ERSPs at the occipital cluster with respect to the grand average of all conditions and with respect to the 1.0m/s speed condition. (**A**) Averaged ERSP at different speeds at the occipital cluster. Red indicated spectral power increases (neural synchronization) and blue indicated spectral power decreases (neural desynchronization) relative to the grand average of all conditions. Vertical dashed lines indicated gait events. RHS: right heel strike; LTO: left toe off; LHS: left heel strike; RTO: right toe off. (**B**) Significant effect of terrain on ERSPs across gait cycle with non-parametric statistics, with red indicating significance (p<0.05). (**C**) ERSPs with respect to 1.0 m/s speed condition. Regions that are not significantly different from 1.0 m/s condition have a semi-transparent mask.

**S10 Fig.**
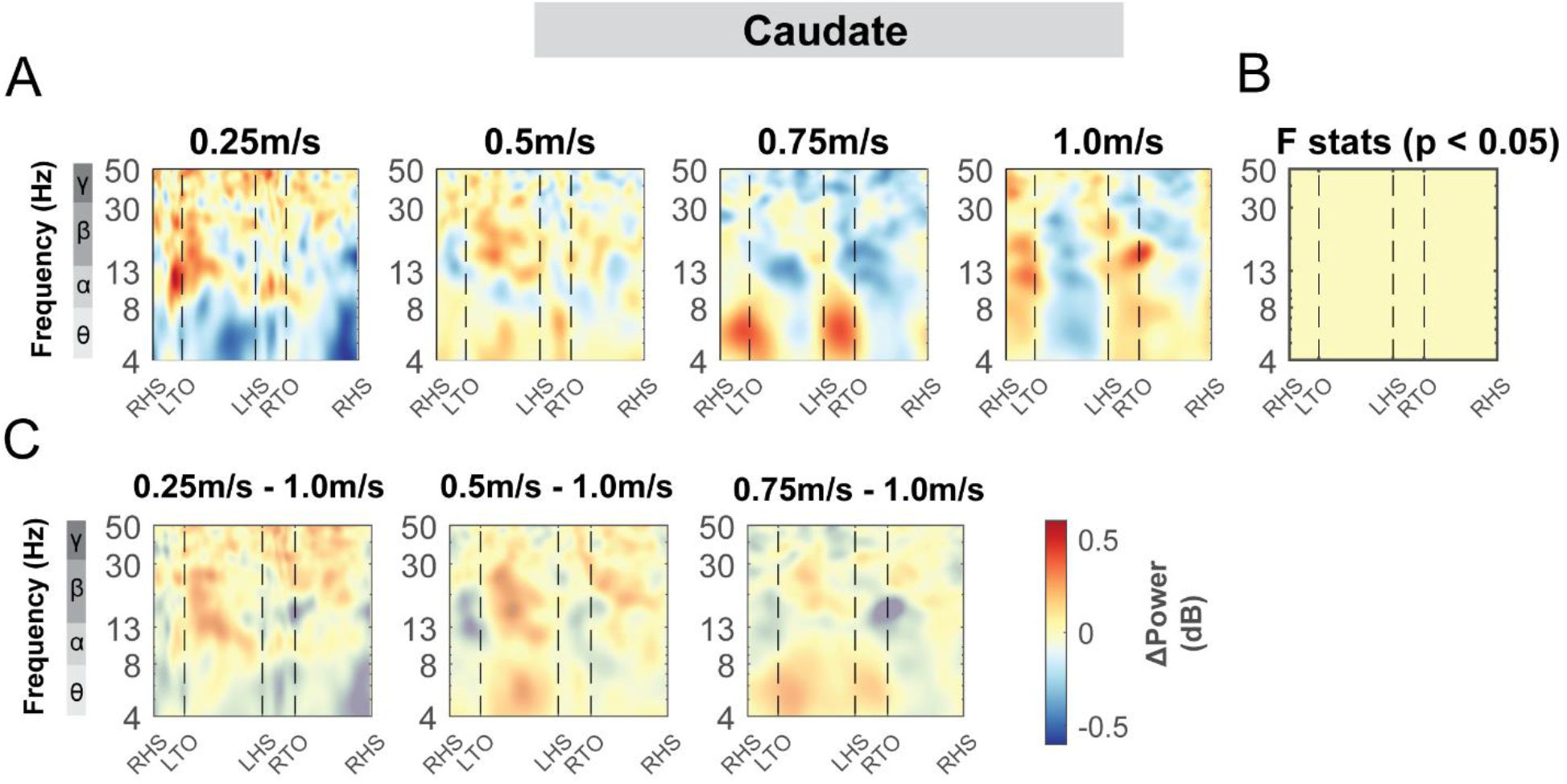
: ERSPs at the caudate cluster with respect to the grand average of all conditions and with respect to the 1.0m/s speed condition. (**A**) Averaged ERSP at different speeds at caudate cluster. Red indicated spectral power increases (neural synchronization) and blue indicated spectral power decreases (neural desynchronization) relative to the grand average of all conditions. Vertical dashed lines indicated gait events. RHS: right heel strike; LTO: left toe off; LHS: left heel strike; RTO: right toe off. (**B**) Significant effect of terrain on ERSPs across gait cycle with non-parametric statistics, with red indicating significance (p<0.05). (**C**) ERSPs with respect to 1.0 m/s speed condition. Regions that are not significantly different from 1.0 m/s condition have a semi-transparent mask.

## Discussion

Our study’s primary objective was to determine how electrocortical activity measured by EEG changed with parametric variations in terrain unevenness for neurotypical young adults. We identified multiple brain regions associated with uneven terrain walking. We found that alpha and beta spectral power were lower with greater terrain unevenness at the sensorimotor and posterior parietal areas while theta spectral power was greater in the mid/posterior cingulate area with greater terrain unevenness. We also observed that gait-related spectral power fluctuations changed with terrain unevenness in all identified brain clusters. Our secondary goal was to determine how electrocortical activity changed with walking speed. Contrary to our speed hypothesis, we found that alpha and beta average spectral power did not change with walking speed in the sensorimotor areas. We only observed an effect of speed on gait-related spectral power fluctuations in the sensorimotor area but not much in other brain areas. These results suggest that distinct cortical processes may be recruited for walking over uneven terrain versus flat terrain at different speeds. This also confirms that the observed cortical changes during uneven terrain walking were not related to the variability in walking speed between participants. Slower gait speed and difficulty in traversing uneven terrain are both characteristics of mobility impairment as humans get older as well as in other conditions known to impact mobility (e.g., chronic pain, post-stroke hemiparesis, lower limb amputation)[2,37,38]. Providing this baseline data of electrocortical dynamics in neurotypical young participants would help better understand cortical deficiencies that occur across the lifespan and lead to a better understanding of neural compensation for future studies [8,36]. Our results could also help reinforce the importance of cortical involvement in the control of human walking, often characterized as primarily dependent on reflex activation and spinal neural networks.

### Alpha and beta power decrease with terrain unevenness

Consistent with our hypothesis, alpha and beta spectral power across the gait cycle were lower with greater terrain unevenness at the sensorimotor area. Alpha oscillations are considered to reflect an ‘idling’ state of the brain [39] and lower alpha band power indicates active cortical processing. In the sensorimotor region, alpha and beta band power decreased following movement initiation is attributed to an increased cortical neuron activity for motor planning and execution [40,41]. Our results suggest that cortical involvement was greater during uneven terrain walking versus walking on flat surfaces. Beta power reduction during uneven terrain walking was prominent during the contralateral limb swing phase before foot placement in our study; however, Jacobsen et al. found beta power reduction only following foot strike [15].

Several factors may contribute to the discrepancy. The terrain was more challenging in our study compared to that in Jacobsen et al. as they compared paved overground concrete terrain with unpaved grassy terrain. It seems probable that walking on our uneven terrain treadmill required more cortical processing regarding movement adjustments during swing compared to walking on the unpaved grassy terrain. We studied the same walking speed on the treadmill for all terrain conditions while participants walked more slowly on the unpaved grassy terrain than the paved concrete terrain in the overground study by Jacobsen et al.[15]. Lastly, we used a clustering approach to group the brain components based on each source’s location before computing the average spectral power across participants. In contrast, Jacobsen et al. computed the spectral power modulation in the channel-space at the Cz electrode, which limits their interpretation of the location of the sources. As a result, beta power reduction at Cz electrode following right foot strike may have contributions from other non-sensorimotor areas.

The posterior parietal area demonstrated sustained alpha and beta power desynchronization across the gait cycle during uneven terrain walking compared to walking on a flat surface (Fig. 6). The posterior parietal area is associated with multisensory integration and estimation of an obstacle’s location relative to the body’s current state to appropriately modify the gait pattern [42,43]. Lower alpha and beta power may reflect cortical processing of multi-sensory modalities (i.e. vision, vestibular, and proprioception) to maintain balance when walking on an uneven terrain. In addition, a sustained decrease in alpha power across the gait cycle in the posterior parietal area could be attributed to greater attention to balance control and greater alertness to threat perception during uneven terrain walking compared to walking on an even surface [44].

Greater attention can help prioritize task-relevant sensory processing during gait to filter task-irrelevant stimuli or noise [44]. Therefore, alpha power change at the posterior parietal area can potentially be used as a cortical marker of sensorimotor attention and alertness during gait for future studies that investigate sustained attention and its relationship with mobility deficits.

### Theta band power increase with terrain unevenness

Inconsistent with our hypothesis, we did not find a cluster at the anterior cingulate area but rather at the mid/posterior cingulate area, which plays a role in somatosensory processing [45] and orientation of the body in space to sensory stimuli [46]. Multiple other EEG studies have reported mid/posterior cingulate involvement during locomotor tasks that required balance control. Sipp et al. identified a posterior cingulate cluster during a narrow beam walking task in which theta band power significantly increased following loss of balance [17]. In a different study, participants walked in a split-belt environment where one belt speed moved faster than the other. Both anterior and posterior cingulate clusters showed strong theta synchronization during early adaptation (when balance was challenged) versus pre-adaptation (when participants walked with tied-belt speed) [47]. It is likely that mid/posterior cingulate area receives multi-sensory input from sensorimotor cortex and parietal cortex to guide body orientation and movements during balance challenging tasks [46,48]. Future studies should investigate the effective connectivity between the mid/posterior cingulate and sensorimotor area and posterior parietal to better determine the role of the mid/posterior cingulate in maintaining balance during gait.

On more uneven terrain, greater theta band power was slightly associated with greater terrain unevenness in the mid/posterior cingulate cluster (Fig. 7). Additionally, we found theta synchronization across the gait cycle during the most challenging terrain condition compared to the flat condition in the left sensorimotor, left posterior parietal, mid/posterior cingulate, left pre-supplementary motor, right premotor, and occipital cluster. These results indicated that theta synchronization is widely distributed across the brain, and a higher level cognitive control is needed during the more complex locomotor task [48,49]. Additionally, theta power may facilitate multisensory integration during movement. For example, theta oscillation at the temporal lobe occurred more often in a participant who is congenitally blind compared to other normally sighted participants when walking in a room, indicating that theta power was associated with somatosensory processing during movement [50]. Together, greater theta power when walking on a more uneven surface may be attributed to a greater need for sensory processing to maintain balance during uneven terrain walking [49].

### Gait-related spectral power fluctuations

Patterns of event-related spectral power fluctuations in the sensorimotor area are in line with previous literature using both non-invasive and invasive recordings [29,51–54]. The power spectral fluctuations computed with respect to the average spectral power within each condition showed significant alpha and beta desynchronization during contralateral swing phase and synchronization during contralateral limb stance phase and push-off (Fig. 9). The spectral power fluctuation profile at the sensorimotor area found in this study is consistent with the neural activation profiles classically recorded in rats, rabbits, cats, and nonhuman primates [55–57].

Neuron firing rates peaked during the gait phase transition and swing phase in rats, and likewise, cortical motor neurons in cats and primates also demonstrated increased firing rates toward push-off phase and swing phase [44–46]. These results suggested increased cognitive processing for movement planning during the swing phase.

We also observed gait-related spectral power fluctuations at the non-sensorimotor areas when participants walked on both flat surfaces and uneven terrain. For instance, there was rhythmic modulation of power spectral fluctuations at the left and right posterior parietal clusters during walking on a flat surface (Fig. 9). Such modulation is similar to that observed in cats [58,59] where neural population peak activity occurred during the swing phase of the contralateral forelimb at the area 5 of the posterior parietal cluster [58,59]. These results suggested that other brain areas may receive movement-related information from the sensorimotor area during locomotion. However, we also observed some differences between gait-related spectral fluctuations between the non-sensorimotor areas and sensorimotor areas. One difference is that we did not observe strong lateralization at the non-sensorimotor areas. This is likely because fewer limb-dependent cells exist at higher-level brain centers compared to the motor cortex [58]. A group of limb-independent cells was only found in the posterior parietal cortex of cats that discharged related to the lead limb but not related to the side of the limb. Also, gait-related power fluctuations in the posterior parietal area were not as robust as in the sensorimotor area.

One potential explanation is that a smaller portion of the local neural population may be rhythmically modulated during locomotion and only engaged when more precise control of whole-body movement is needed [58].

### Use of visual information during uneven terrain walking

Visual information is critical for people to plan their movement when walking over uneven terrain. This is evidenced by a greater theta, and lower alpha and beta band spectral power at the occipital area during uneven terrain walking compared to flat terrain (Fig. 8). In addition, we also observed gait-related theta synchronization and gamma synchronization in all levels of uneven terrain versus flat terrain (S4 Fig). A visual stimulus, particularly a moving stimulus, leads to changes in gamma band activity in the occipital area [60,61]. We did not instruct participants where they should look while walking, but the rigid, colored pucks were at least in their peripheral vision and may have induced changes in gamma band activity. EEG signals in the gamma band during walking are often contaminated by muscle artifacts such as neck muscle activity. Still, the gamma activity cannot be fully attributed to muscle artifacts because gait-related spectral power fluctuations in the gamma band within the occipital area differ substantially from neck muscle activity spectral power fluctuations across the gait cycle [29].

### Speed modulation of electrocortical dynamics

Contrary to our hypothesis, we only observed small effect of walking speed on average theta band power but not on alpha and beta band power at the sensorimotor clusters. For intra-stride spectral power fluctuations, there was significantly greater alpha and beta desynchronization during the contralateral swing phase only at 0.25 m/s versus 1.0 m/s. These findings suggest that maintaining a very slow speed (0.25 m/s) on the treadmill may require substantially more cortical processing and attentional resources for movement planning and execution in younger adults. This finding may be inconsistent with previous studies that found that fast gait speeds reduced sensorimotor alpha and beta power substantially [29]. Several reasons may explain the apparent discrepancy. First, our range of speeds was from 0.25 m/s to 1.0 m/s while previous studies focused on speeds that were higher than 0.5 m/s. For example, Nordin et al. studied the range of 0.5 m/s to 2.0 m/s [29]. The speeds we used in this study, particularly 0.25 m/s, were much slower than normal self-selected speeds in young adults (∼1.3 m/s) and thus may require added cortical processing to maintain the very slow speed. Second, our study had a much larger sample size (n = 32) than previous mobile EEG studies (n = ∼10). We used a similar processing pipeline, but included individual-specific head models that improved source localization [62].

This enabled us to have a better estimation of sensorimotor source locations than the previous study [29]. We speculate that there might be a nonlinearity in the speed modulation of electrocortical dynamics such that only extremely slow or fast walking speeds may substantially affect electrocortical activities. This remains to be tested.

We only observed substantial intra-stride gait-related spectral power modulations with speed at the sensorimotor area but no other brain areas. Although average beta band spectral power was negatively associated with faster gait speed at the right posterior parietal cluster and mid/posterior cingulate cluster, the effect size was small. We did not find any changes in intra-stride gait-related spectral power fluctuations in the alpha or beta band at the posterior parietal, mid/posterior cingulate, premotor, and pre-supplementary motor areas as we observed during uneven terrain walking. It does not necessarily mean that other brain areas were not involved in gait speed modulation. For example, prefrontal area activation recorded with functional near-infrared spectroscopy increased in older adults during fast walking [63]. However, prefrontal activity can be difficult to obtain with mobile EEG due to ocular artifacts and facial muscle activities. Better signal processing or new hardware design may be needed to identify prefrontal activity with mobile EEG. Together, our results suggest that there might be distinct neural processes that assist balance control during uneven terrain walking and adjusting gait speeds.

### Limitations

There are several limitations to our study. The range of walking speeds selected for the speed condition was slower than the typical self-selected treadmill walking speed in young adults [22,64]. We did not collect walking trials with speeds over 1.0 m/s or with self-selected treadmill speeds for young adults because this study was part of a larger study that also aimed to recruit many older adults (>70yrs old) who were less likely to be able to walk more than 1.0 m/s [64].

To allow for comparison between young and older adults, the walking speeds were set from 0.25 m/s to 1.0 m/s. It is also important to note that we focused on cortical areas. Electrical activity at subcortical areas and at the cerebellum is rarely to be identified with scalp EEG during human walking. We identified a cluster in the caudate area with the help of the individual-specific head model, but our focus was on cortical spectral power fluctuations. Thus, the current paper only reported results in the caudate cluster rather than have a hypothesis about how activities at caudate would change with terrain unevenness and speed. Also, future studies may investigate gaze control during uneven terrain walking with brain dynamics. Visual control strategies can change based on terrain. A recent study found that natural uneven terrain led participants to look two steps forward while maintaining a constant ahead-looking window to gather information about their surroundings and adjust their gait pattern to maintain balance [13]. Treadmill locomotion places a limit on the forward distance available for gazing which may affect motor planning for foot placement.

## Materials and Methods

### Participant

We recruited a total of 35 healthy young individuals (19 females, mean age 24 +/- 4 yrs, walking speed on uneven terrain = 0.7 +/- 0.2 m/s) with no musculoskeletal, severe cardiovascular, orthopedic, or psychiatric diagnosis as part of a larger parent study (i.e., the Mind in Motion study (NCT03737760). Full inclusion and exclusion criteria were reported by Clark et al. [32].

Three participants reported being left-handed. A recent paper using EEG for mobile brain imaging found a Cohen’s d of 1.22 for comparing electrocortical power fluctuations between walking on a paved concrete surface and a grassy unpaved surface [15]. Based on that effect size and their data, we aimed for 30 participants so that we would have a minimum of 15 participants’ data in each independent component cluster for EEG analysis. We recruited 35 participants considering a ∼10% - 15% drop-out rate due to gelling or noise artifacts based on previous studies in our lab [47,65]. We removed one female participant from the analysis due to difficulty with electrode gelling during data collection. We also removed two female participants due to technical issues with the data. All participants provided informed consent before participating in the experiment. The study was conducted in accordance with the Declaration of Helsinki and approved by the Institutional Review Board of the University of Florida (IRB 201802227).

### Design of uneven terrain treadmill

Details of the terrain design have been described previously [2]. We modified the terrain unevenness using rigid foam disks that were attached to the slat belt treadmill (PPS 70 Bari-Mill, Woodway, Waukesha, WI, USA; 70 cm x 173 cm walking surface; Fig. 1). The rigid disks were made from polyurethane using a circular free-rise mold with a diameter of 12.7 cm. Hoop-and-loop fasteners attached the disks to the treadmill surface so that we could easily switch the level of terrain unevenness. The spatial configuration of the disks was the same for each uneven terrain and consistent across participants. There were no large gaps between disks, so participants stepped on at least one disk with almost all footfalls.

We parametrically varied the terrain unevenness for low, medium, and high levels by altering the height of the rigid disks on the treadmill. The low terrain included yellow 1.3 cm high disks. The medium terrain included orange disks of two different heights: 50% of them were 1.3 cm high disks and 50% were 2.5 cm high. The high terrain consisted of red disks of three different heights: 20% were 1.3 cm tall, 30% were 2.5 cm tall, and 50% were 3.8 cm tall. There were no disks for the flat terrain, but there were painted green circles on the treadmill in the same configuration as the other terrains.

### Experimental protocol

The experimental protocol is a subset of the Mind in Motion larger study. Details of the full protocol were provided in [36]. We included one session of EEG and one session of MRI scans for this experiment protocol. The EEG and MRI sessions were performed on separate days within approximately 30 days or less of each other.

The EEG visit included treadmill walking trials on four different levels of uneven terrain (flat, low, medium, and high), treadmill walking at four different speeds (0.25 m/s, 0.5 m/s, 0.75 m/s, 1.0 m/s) on the flat surface, and one seated resting trial (Fig. 1A). Before the EEG visit, participants walked on an overground version of the Flat, Low, Medium, and High Terrain on a 3.5 m mat three times. We instructed the participants to walk at a normal, comfortable pace. The overground speed for each terrain was computed as the average speed to walk through the middle 3-meter portion. We set the treadmill walking speed across all terrains to 75% of the slowest overground speed (slowest terrain) because the treadmill walking speed was approximately 10-15% slower than the overground walking speed and due to safety concerns for uneven terrain walking [66,67]. Participants wore a harness to prevent falling to the ground, but the harness did not provide any body weight support unless they fell. We did not record any falls in this dataset. Participants performed a block of two treadmill walking trials per condition. Each walking trial was 3 minutes. We pseudorandomized the conditions with 8 unique orders of uneven terrain conditions and speed conditions, respectively. The total amount of data collected including both walking trials and the resting trial was approximately 50 minutes.

### Data Acquisition

During EEG visit, participants wore a custom-made dual-layer EEG cap (ActiCAP snap sensors; Brain Products GmbH, Germany), including 120 scalp electrodes and 120 mechanically coupled noise electrodes. The scalp electrodes followed a 10-05 electrode system. We inverted and mechanically coupled noise electrodes to the scalp electrodes [68,69]. We used a conductive fabric as an artificial skin circuit and bridged the noise electrodes. We re-purposed eight of the original 128 scalp electrodes (TP9, P9, PO9, O9, O10, PO10, P10, and TP10) to measure muscle activity of the sternocleidomastoid and trapezius on the left and right sides. We aimed to keep all scalp electrode impedance values below 15 Kohm during the setup. Ground and reference electrodes were kept below 5 Kohm. We digitized the electrode locations using a structural scanner (ST01, Occipital Inc., San Francisco, CA). We used four LiveAmp 64 amplifiers and logged EEG data at 500 Hz. The online reference and ground electrodes were at CPz and Fpz, respectively.

In addition, we recorded the ground reaction force of each foot with capacitive shoe insole sensors (loadsol 1- 184 sensor, Novel Electronics Inc., St. Paul, MN, USA) at 200Hz and sacrum kinematics with an inertial measurement unit (IMU, Opal APDM Inc., Portland, OR, USA) at 128 Hz. We synchronized data from the IMU to the EEG LiveAmp offline via a pulse at the beginning and the end of the trial. We also synchronized the insole sensor force data with EEG LiveAmp offline via pulses occurring every five seconds. More details about the data synchronization were provided in our previous paper [2].

### MRI Acquisition

On a separate day, we collected structural MRI scans for all participants. We acquired the anatomical brain structure from a T1-weighted sequence. The parameters for this anatomical image were: repetition time (TR) = 2000ms, echo time (TE) = 2.99ms, flip angle = 8°, voxel resolution = 0.8mm^3^, field of view = 256 × 256 × 167 mm^2^ (4:22 minutes of scan time), using a 64-channel coil array on a 3T Siemens MAGNETOM Prisma MR scanner.

### Data Processing

#### Behavioral analysis

We post-processed the kinematic and kinetic data in Matlab 2020b (Mathworks, USA) to compute the variables of interest. We defined foot strike as the point when ground reaction forces became greater than 20N and foot off as the point when ground reaction forces became less than 20N [2]. We defined the step duration as the time between consecutive foot strikes. We also computed the peak-to-peak excursion of the sacrum in the anteroposterior and mediolateral direction using the IMU data. Details of the calculations were reported in our recent paper [2].

We removed outliers that were +/-2.5 standard deviations away from the mean [2]. We calculated the variability of each of these measures as the coefficient of variation (standard deviation over mean).

#### EEG data pre-processing

We processed all EEG data using custom Matlab scripts (R2020b) and EEGLAB (v 2021.0; Fig. 2) [30]. We first applied a 1Hz high-pass filter (-6dB at 0.5Hz) with *eegfiltnew* on all scalp, noise, and muscle channels to remove drift for each trial. We also applied a 20Hz low-pass filter with *eegfiltnew* on muscle channels. We used the CleanLine plugin in EEGLAB to remove line noise at 60Hz and 120Hz. We rejected bad channels that were 3 standard deviations away from the mean of EEG and noise channels, respectively. We performed average reference for scalp, noise, and muscle channels respectively.

We then used a novel algorithm iCanClean [70,71] to remove artifacts that were highly correlated with noise reference electrodes (R^2^ = 0.65 with a four-second moving window) and muscle reference electrodes (R^2^ = 0.4 with four-second moving window). This algorithm has been previously validated to improve mobile EEG data quality during human experiments [69,71]. Then, we used *clean_artifacts* in EEGLAB to remove bad channels and noisy time frames using default parameters except for the following parameters (chan_crit1 = 0.5, win_crit1=0.4, winTol = [-Inf, 10]). These parameters were selected in a preliminary analysis of a subset of the data, which aimed to minimize the number of channels and time frames rejected while maximizing a good number of brain components by ICLabel [62,72]. We retained 110 ± 6 channels and rejected a maximum of 5% of time frames (mean = 1%). Scalp EEG data were re-referenced again. We performed adaptive mixture independent analysis (AMICA) on the preprocessed data [73] to decompose the preprocessed EEG data into statistically independent components. We later used the independent components to perform source localization.

### Individual-specific volume conduction model

We processed the T_1_-weighted MRI using Fieldtrip (v.20210910) for each participant [62]. The images were resliced to be isotropic (1 mm^3^). We digitized the fiducial locations (left/right preauricular, nasion) on the MRI. We used the *headreco* from SimNIBS toolbox (v 3.2) to perform tissue segmentation [74]. We segmented individual MRIs into six tissue layers (scalp, skull, air, cerebrospinal fluid, gray matter, and white matter). Hexahedral meshes were generated with recommended node-shift parameters using *prepare_mesh_hexahedral.* We co-registered the digitized electrode locations to the individual-specific head model by aligning the fiducial locations digitized in the MRIs to those in the structural scan. We calculated the leadfield matrix for each individual-specific head model using the SIMBIO toolbox in Fieldtrip. We distributed source positions in the gray matter 5 mm apart.

### Source localization

We performed EEG source localization using an equivalent dipole fitting approach using *ft_dipolefitting* function in the Fieldtrip toolbox with the individual-specific volume conduction head models. We then converted the dipole locations to the Montreal Neurological Institute (MNI) template. We retained brain components using the following criteria [62]: 1) ICLabel [72] (version: lite) classified the brain probability of greater or equal to 50%, 2) negative slope of the power density spectrum for 2 - 40 Hz to remove muscle components, 3) residual variance of dipole fitting <15%, 4) minimal high-frequency power coupling using PowPowCAT toolbox to further remove muscle components [75], and lastly, we visually inspected all the components and removed non-brain components based on previous criteria. We retained 14 ± 5 brain components per participant.

### K-means clustering of brain components

We clustered the brain components by dipole location using k-means in EEGLAB. We determined the optimal number of clusters (k = 12) using Silhoutte and DaviesBoukin criteria in *evalcluster*. We retained the clusters with more than half of the participants. Components that are further than three standard deviations away from any of the clusters were identified as outliers. In the case that multiple components per subject existed in a cluster, we selected the component with the lowest residual variance to prevent artificially inflating the sample size [65].

### Computing power spectral density and event-related spectral perturbations

We then performed frequency and time-frequency analyses for each cluster. For the walking trials, data were segmented into epochs of 5s (from 0.5s before to 4.5s after the right foot strike). The epoch length was chosen to accommodate participants with a long step duration of 0.25 m/s. We rejected epochs that were three standard deviations from the mean gait event time and 10% of epochs that had the highest voltage maximal, resulting in 205 ± 30 epochs for flat, 202 ± 31 epochs for low, 205 ± 30 epochs for medium, 208 ± 37 epochs for high terrain, and 108 ± 18 epochs for 0.25 m/s, 170 ± 17 epochs for 0.5 m/s, 210 ± 23 epochs for 0.75 m/s, and 249 ± 18 for 1.0 m/s. We found fewer epochs at slower speeds as the number of steps reduced for a trial with fixed duration.

For the frequency analysis, we computed the log power spectral density (PSD) for independent components in each cluster and normalized by subtracting each individual’s mean log spectral power density from all conditions. We used the FOOOF toolbox[34] to separate the aperiodic and periodic components of each power spectra from 3 to 40Hz (peak width limits: [1 8], minimum peak height: 0.05, maximum number of peaks: 2). We computed the flattened power spectral density by subtracting the aperiodic component from each of the original power spectral density. Lastly, we computed average power for each band using the flattened power spectral density.

We then assessed the electrocortical activity tied to gait events using event-related spectral perturbations (ERSPs). We computed the single trial spectrograms with *newtimef* (Morlet wavelets cycles: [3 0.8]). ERSPs were time-warped to the gait cycle from right foot-strike to the subsequent right foot-strike. ERSPs were normalized to 1) average spectral power across all gait cycles within conditions and 2) average spectral power across gait cycles for all conditions (common baseline removal). We averaged ERSPs for each participant and then averaged across all participants for each cluster and for each condition.

### Statistical analysis

All statistical analyses were performed in Matlab 2020b (Mathworks). We first assessed whether there were significant differences in any behavioral measures across terrain using a linear mixed effect model for each outcome measure. The dependent variables included step duration, step duration coefficient of variation, and sacral excursion coefficient of variation in mediolateral and anteroposterior directions. The independent variables included Terrain (flat, low, medium, high). Walking speed was included as a covariate for all mixed effect models. We included a random intercept to account for unmodeled sources of between-subject variability. We calculated Cohen’s f^2^ as a measure of effect size for the main effects and used the following definitions for the effect sizes: 0.02 = small, 0.15 = medium, 0.35 = large effect size [76]. We analyzed the residual normality using the Lilifores Test. Pairwise comparison was corrected for multiple comparison using false discovery rate [77]. Significance level was kept α < 0.05.

We then assessed if power spectral density for each cluster differed across terrain and speed, respectively. We performed non-parametric permutation statistics for both the original and flattened power spectral density using Fieldtrip in EEGLAB (α = 0.05, 2000 iterations) and corrected for multiple comparisons using false discovery rate. We also analyzed if average power for theta, alpha, and beta band for each cluster after removing the aperiodic component differed across terrain and speed using a linear mixed model for each dependent variable. The independent variables included Terrain or Speed. Terrain was modeled as a categorical variable while Speed was a continuous variable. We included a random intercept for each model. The reference level was set to flat condition or 1.0 m/s condition. We used 1.0 m/s as the reference level because this speed was closer to the self-selected walking speed in young adults [22,64].

For ERSPs, we first assessed the statistically significant time-frequency change within each condition. The spectral baseline was the average spectral power across all gait cycles within the conditions. For each condition in each cluster, we bootstrapped ERSPs (α = 0.05, 2000 iterations) and corrected for multiple comparison using false discovery rate. We then assessed the differences in ERSPs across terrain and speed, respectively. We performed non-parametric permutation statistics to determine the time-frequency differences across terrain conditions and relative to flat condition (ERSP_terrain_ – ERSP_flat_). Similarly, we performed permutation statistics to determine time-frequency differences across speed conditions and relative to 1.0 m/s condition (ERSP_speed_ – ERSP_1.0m/s_). All these analyses were corrected with cluster-based multiple comparison using Fieldtrip through EEGLAB (α = 0.05, 2000 iterations).

## Data availability

Data is available via OpenNeuro: https://openneuro.org/crn/reviewer/eyJhbGciOiJIUzI1NiIsInR5cCI6IkpXVCJ9.eyJzdWIiOiI3ODVlMGUyNC1lOTQ3LTRkZDEtYmQxZi1mNDJhMTVlYjI1OGQiLCJlbWFpbCI6InJldmlld2VyQG9wZW5uZXVyby5vcmciLCJwcm92aWRlciI6Ik9wZW5OZXVybyIsIm5hbWUiOiJBbm9ueW1vdXMgUmV2aWV3ZXIiLCJhZG1pbiI6ZmFsc2UsInNjb3BlcyI6WyJkYXRhc2V0OnJldmlld2VyIl0sImRhdGFzZXQiOiJkczAwNDYyNSIsImlhdCI6MTY4ODI2NjgzNCwiZXhwIjoxNzE5ODAyODM0fQ.r9jdMAJJzGPo6tQ0JahaJbFRqym4ywdp8U1dY73oX7M

## Acknowledgement

We would like to thank HNL lab members for their help with data collection: Ryland Swearinger, Madison Tenerowicz, Quinlan Degnan, Sydney Irwin and HNL members for the insightful discussion that helped improve the paper. We would also like to thank our study coordinators who made a huge effort recruiting participants, particularly during the Covid-19 pandemic.

## Supplementary Information

S1 Fig: ERSPs at the left pre-supplementary motor (A), right premotor (B), occipital (C), and caudate clusters (D) with respect to the average of each condition at different terrains.

S2 Fig: ERSPs at the left pre-supplementary motor clusters with respect to the grand average of all conditions and with respect to flat terrain condition.

S3 Fig: ERSPs at the right premotor clusters with respect to the grand average of all conditions and with respect to flat terrain condition.

S4 Fig: ERSPs at the occipital cluster with respect to the grand average of all conditions and with respect to flat terrain condition.

S5 Fig: ERSPs at the caudate cluster with respect to the grand average of all conditions and with respect to flat terrain condition.

S6 Fig: ERSPs at the left pre-supplementary motor (A), right premotor (B), occipital (C), and caudate areas (D) with respect to the average of each condition at different speeds.

S7 Fig: ERSPs at the left pre-supplementary motor area with respect to the grand average of all conditions and with respect to 1.0m/s speed condition.

S8 Fig: ERSPs at the right premotor area with respect to the grand average of all conditions and with respect to 1.0m/s speed condition.

S9 Fig: ERSPs at the occipital cluster with respect to the grand average of all conditions and with respect to 1.0m/s speed condition.

S10 Fig: ERSPs at the caudate cluster with respect to the grand average of all conditions and with respect to 1.0m/s speed condition.

## Reference

1. Coleman TD, Lawrence HJ, Childers WL. Standardizing Methodology for Research with Uneven Terrains Focused on Dynamic Balance During Gait. Journal of Applied Biomechanics. 2016;32: 599–602. doi:10.1123/jab.2016-0014

2. Downey RJ, Richer N, Gupta R, Liu C, Pliner EM, Roy A, et al. Uneven terrain treadmill walking in younger and older adults. PLOS ONE. 2022;17: e0278646. doi:10.1371/journal.pone.0278646

3. Kent JA, Sommerfeld JH, Mukherjee M, Takahashi KZ, Stergiou N. Locomotor patterns change over time during walking on an uneven surface. Journal of Experimental Biology. 2019;222: jeb202093. doi:10.1242/jeb.202093

4. Thomas NDA, Gardiner JD, Crompton RH, Lawson R. Physical and perceptual measures of walking surface complexity strongly predict gait and gaze behaviour. Human Movement Science. 2020;71: 102615. doi:10.1016/j.humov.2020.102615

5. Voloshina AS, Ferris DP. Biomechanics and energetics of running on uneven terrain. The Journal of experimental biology. 2015;216: 711–719. doi:10.1242/jeb.106518

6. Clark DJ. Automaticity of walking: functional significance, mechanisms, measurement and rehabilitation strategies. Frontiers in Human Neuroscience. 2015;9. Available: https://www.frontiersin.org/articles/10.3389/fnhum.2015.00246

7. Lau TM, Gwin JT, Ferris DP. Walking reduces sensorimotor network connectivity compared to standing. J NeuroEngineering Rehabil. 2014;11: 14. doi:10.1186/1743-0003-11-14

8. Fettrow T, Hupfeld K, Tays G, Clark DJ, Reuter-Lorenz PA, Seidler RD. Brain activity during walking in older adults: Implications for compensatory versus dysfunctional accounts. Neurobiol Aging. 2021;105: 349–364. doi:10.1016/j.neurobiolaging.2021.05.015

9. Clark DJ, Christou EA, Ring SA, Williamson JB, Doty L. Enhanced Somatosensory Feedback Reduces Prefrontal Cortical Activity During Walking in Older Adults. J Gerontol A Biol Sci Med Sci. 2014;69: 1422–1428. doi:10.1093/gerona/glu125

10. Clark DJ, Rose DK, Ring SA, Porges EC. Utilization of central nervous system resources for preparation and performance of complex walking tasks in older adults. Frontiers in Aging Neuroscience. 2014;6. Available: https://www.frontiersin.org/articles/10.3389/fnagi.2014.00217

11. Hawkins KA, Clark DJ, Balasubramanian CK, Fox EJ. Walking on uneven terrain in healthy adults and the implications for people after stroke. NeuroRehabilitation. 2017;41: 765. doi:10.3233/NRE-172154

12. Darici O, Kuo AD. Humans optimally anticipate and compensate for an uneven step during walking. Elife. 2022;11: e65402. doi:10.7554/eLife.65402

13. Matthis JS, Yates JL, Hayhoe MM. Gaze and the Control of Foot Placement When Walking in Natural Terrain. Current Biology. 2018;28: 1224–1233.e5. doi:10.1016/j.cub.2018.03.008

14. Bruijn SM, Van Dieën JH, Daffertshofer A. Beta activity in the premotor cortex is increased during stabilized as compared to normal walking. Front Hum Neurosci. 2015;9: 593. doi:10.3389/fnhum.2015.00593

15. Jacobsen NSJ, Blum S, Scanlon JEM, Witt K, Debener S. Mobile electroencephalography captures differences of walking over even and uneven terrain but not of single and dual-task gait. Front Sports Act Living. 2022;4: 945341. doi:10.3389/fspor.2022.945341

16. Luu TP, Brantley JA, Nakagome S, Zhu F, Contreras-Vidal JL. Electrocortical correlates of human level-ground, slope, and stair walking. PLOS ONE. 2017;12: e0188500. doi:10.1371/journal.pone.0188500

17. Sipp AR, Gwin JT, Makeig S, Ferris DP. Loss of balance during balance beam walking elicits a multifocal theta band electrocortical response. J Neurophysiol. 2013;110: 2050– 2060. doi:10.1152/jn.00744.2012

18. Solis-Escalante T, Stokkermans M, Cohen MX, Weerdesteyn V. Cortical responses to whole-body balance perturbations index perturbation magnitude and predict reactive stepping behavior. European Journal of Neuroscience. 2021;54: 8120–8138. doi:10.1111/ejn.14972

19. Varghese JP, Marlin A, B. Beyer K, Staines WR, Mochizuki G, McIlroy WE. Frequency characteristics of cortical activity associated with perturbations to upright stability. Neuroscience Letters. 2014;578: 33–38. doi:10.1016/j.neulet.2014.06.017

20. Peterson SM, Ferris DP. Differentiation in Theta and Beta Electrocortical Activity between Visual and Physical Perturbations to Walking and Standing Balance. eNeuro. 2018;5: ENEURO.0207-18.2018. doi:10.1523/ENEURO.0207-18.2018

21. Cevallos C, Zarka D, Hoellinger T, Leroy A, Dan B, Cheron G. Oscillations in the human brain during walking execution, imagination and observation. Neuropsychologia. 2015;79: 223–232. doi:10.1016/j.neuropsychologia.2015.06.039

22. Liu C, Park S, Finley J. The choice of reference point for computing sagittal plane angular momentum affects inferences about dynamic balance. PeerJ. 2022;10: e13371. doi:10.7717/peerj.13371

23. Franz JR, Kram R. Advanced age affects the individual leg mechanics of level, uphill, and downhill walking. Journal of Biomechanics. 2013;46: 535–540. doi:10.1016/j.jbiomech.2012.09.032

24. Neptune RR, Sasaki K, Kautz SA. The effect of walking speed on muscle function and mechanical energetics. Gait Posture. 2008;28: 135–143. doi:10.1016/j.gaitpost.2007.11.004

25. Hak L, Houdijk H, Beek PJ, Van Dieë JH. Steps to take to enhance gait stability: The effect of stride frequency, stride length, and walking speed on local dynamic stability and margins of stability. PLoS ONE. 2013;8. doi:10.1371/journal.pone.0082842

26. Kang HG, Dingwell JB. Effects of walking speed, strength and range of motion on gait stability in healthy older adults. Journal of Biomechanics. 2008;41: 2899–2905. doi:10.1016/j.jbiomech.2008.08.002

27. Li L, Haddad JM, Hamill J. Stability and variability may respond differently to changes in walking speed. Human Movement Science. 2005;24: 257–267. doi:10.1016/j.humov.2005.03.003

28. Bulea TC, Kim J, Damiano DL, Stanley CJ, Park H-S. Prefrontal, posterior parietal and sensorimotor network activity underlying speed control during walking. Frontiers in Human Neuroscience. 2015;9. Available: https://www.frontiersin.org/articles/10.3389/fnhum.2015.00247

29. Nordin AD, Hairston WD, Ferris DP. Faster Gait Speeds Reduce Alpha and Beta EEG Spectral Power From Human Sensorimotor Cortex. IEEE Transactions on Biomedical Engineering. 2020;67: 842–853. doi:10.1109/TBME.2019.2921766

30. Delorme A, Makeig S. EEGLAB: an open source toolbox for analysis of single-trial EEG dynamics including independent component analysis. J Neurosci Methods. 2004;134: 9–21. doi:10.1016/j.jneumeth.2003.10.009

31. Tzourio-Mazoyer N, Landeau B, Papathanassiou D, Crivello F, Etard O, Delcroix N, et al. Automated Anatomical Labeling of Activations in SPM Using a Macroscopic Anatomical Parcellation of the MNI MRI Single-Subject Brain. NeuroImage. 2002;15: 273–289. doi:10.1006/nimg.2001.0978

32. Papademetris X, Jackowski MP, Rajeevan N, DiStasio M, Okuda H, Constable RT, et al. BioImage Suite: An integrated medical image analysis suite: An update. Insight J. 2006;2006: 209.

33. Mayka MA, Corcos DM, Leurgans SE, Vaillancourt DE. Three-dimensional locations and boundaries of motor and premotor cortices as defined by functional brain imaging: a meta- analysis. Neuroimage. 2006;31: 1453–1474. doi:10.1016/j.neuroimage.2006.02.004

34. Donoghue T, Haller M, Peterson EJ, Varma P, Sebastian P, Gao R, et al. Parameterizing neural power spectra into periodic and aperiodic components. Nat Neurosci. 2020;23: 1655–1665. doi:10.1038/s41593-020-00744-x

35. Gwin JT, Gramann K, Makeig S, Ferris DP. Electrocortical activity is coupled to gait cycle phase during treadmill walking. NeuroImage. 2011;54: 1289–1296. doi:10.1016/j.neuroimage.2010.08.066

36. Clark DJ, Manini TM, Ferris DP, Hass CJ, Brumback BA, Cruz-Almeida Y, et al. Multimodal Imaging of Brain Activity to Investigate Walking and Mobility Decline in Older Adults (Mind in Motion Study): Hypothesis, Theory, and Methods. Frontiers in Aging Neuroscience. 2020;11. Available: https://www.frontiersin.org/article/10.3389/fnagi.2019.00358

37. Ogawa EF, Shi L, Bean JF, Hausdorff JM, Dong Z, Manor B, et al. Chronic Pain Characteristics and Gait in Older Adults: The MOBILIZE Boston Study II. Arch Phys Med Rehabil. 2020;101: 418–425. doi:10.1016/j.apmr.2019.09.010

38. Von Schroeder HP, Coutts RD, Lyden PD, Billings E, Nickel VL. Gait parameters following stroke: A practical assessment. Journal of Rehabilitation Research and Development. 1995;32: 25–31.

39. Mulholland T. Human EEG, behavioral stillness and biofeedback. Int J Psychophysiol. 1995;19: 263–279. doi:10.1016/0167-8760(95)00019-o

40. Deiber M-P, Sallard E, Ludwig C, Ghezzi C, Barral J, Ibanez V. EEG alpha activity reflects motor preparation rather than the mode of action selection. Frontiers in Integrative Neuroscience. 2012;6. Available: https://www.frontiersin.org/articles/10.3389/fnint.2012.00059

41. Pfurtscheller G, Lopes da Silva FH. Event-related EEG/MEG synchronization and desynchronization: basic principles. Clinical Neurophysiology. 1999;110: 1842–1857. doi:10.1016/S1388-2457(99)00141-8

42. Drew T, Marigold DS. Taking the next step: cortical contributions to the control of locomotion. Current Opinion in Neurobiology. 2015;33: 25–33. doi:10.1016/j.conb.2015.01.011

43. Marigold DS, Drew T. Posterior parietal cortex estimates the relationship between object and body location during locomotion. eLife. 2017;6: 1–24. doi:10.7554/eLife.28143

44. Sarter M, Givens B, Bruno JP. The cognitive neuroscience of sustained attention: where top-down meets bottom-up. Brain Res Brain Res Rev. 2001;35: 146–160. doi:10.1016/s0165-0173(01)00044-3

45. Seitz RJ, Roland PE. Vibratory stimulation increases and decreases the regional cerebral blood flow and oxidative metabolism: a positron emission tomography (PET) study. Acta Neurol Scand. 1992;86: 60–67. doi:10.1111/j.1600-0404.1992.tb08055.x

46. Vogt BA. Midcingulate cortex: Structure, connections, homologies, functions and diseases. J Chem Neuroanat. 2016;74: 28–46. doi:10.1016/j.jchemneu.2016.01.010

47. Jacobsen NA, Ferris DP. Electrocortical Activity Correlated with Locomotor Adaptation during Split-belt Treadmill Walking. Journal of Physiology. 2023.

48. Cavanagh JF, Frank MJ. Frontal theta as a mechanism for cognitive control. Trends Cogn Sci. 2014;18: 414–421. doi:10.1016/j.tics.2014.04.012

49. Raghavachari S, Lisman JE, Tully M, Madsen JR, Bromfield EB, Kahana MJ. Theta Oscillations in Human Cortex During a Working-Memory Task: Evidence for Local Generators. Journal of Neurophysiology. 2006;95: 1630–1638. doi:10.1152/jn.00409.2005

50. M. Aghajan Z, Schuette P, Fields TA, Tran ME, Siddiqui SM, Hasulak NR, et al. Theta Oscillations in the Human Medial Temporal Lobe during Real-World Ambulatory Movement. Current Biology. 2017;27: 3743–3751.e3. doi:10.1016/j.cub.2017.10.062

51. Bradford JC, Lukos JR, Ferris DP. Electrocortical activity distinguishes between uphill and level walking in humans. J Neurophysiol. 2016;115: 958–966. doi:10.1152/jn.00089.2015

52. McCrimmon CM, Wang PT, Heydari P, Nguyen A, Shaw SJ, Gong H, et al. Electrocorticographic Encoding of Human Gait in the Leg Primary Motor Cortex. Cerebral Cortex. 2018;28: 2752–2762. doi:10.1093/cercor/bhx155

53. Oliveira AS, Schlink BR, Hairston WD, König P, Ferris DP. Restricted vision increases sensorimotor cortex involvement in human walking. J Neurophysiol. 2017;118: 1943–1951. doi:10.1152/jn.00926.2016

54. Zhao M, Bonassi G, Samogin J, Taberna GA, Pelosin E, Nieuwboer A, et al. Frequency- dependent modulation of neural oscillations across the gait cycle. Human Brain Mapping. 2022;43: 3404–3415. doi:10.1002/hbm.25856

55. Armstrong DM, Drew T. Locomotor-related neuronal discharges in cat motor cortex compared with peripheral receptive fields and evoked movements. The Journal of Physiology. 1984;346: 497–517. doi:10.1113/jphysiol.1984.sp015037

56. Beloozerova IN, Sirota MG. The role of the motor cortex in the control of accuracy of locomotor movements in the cat. J Physiol. 1993;461: 1–25.

57. DiGiovanna J, Dominici N, Friedli L, Rigosa J, Duis S, Kreider J, et al. Engagement of the Rat Hindlimb Motor Cortex across Natural Locomotor Behaviors. J Neurosci. 2016;36: 10440–10455. doi:10.1523/JNEUROSCI.4343-15.2016

58. Andujar J-É, Lajoie K, Drew T. A Contribution of Area 5 of the Posterior Parietal Cortex to the Planning of Visually Guided Locomotion: Limb-Specific and Limb-Independent Effects. Journal of Neurophysiology. 2010;103: 986–1006. doi:10.1152/jn.00912.2009

59. Beloozerova IN, Sirota MG. Integration of Motor and Visual Information in the Parietal Area 5 During Locomotion. Journal of Neurophysiology. 2003;90: 961–971. doi:10.1152/jn.01147.2002

60. Fan J, Byrne J, Worden MS, Guise KG, McCandliss BD, Fossella J, et al. The Relation of Brain Oscillations to Attentional Networks. J Neurosci. 2007;27: 6197–6206. doi:10.1523/JNEUROSCI.1833-07.2007

61. Muthukumaraswamy SD, Singh KD. Visual gamma oscillations: the effects of stimulus type, visual field coverage and stimulus motion on MEG and EEG recordings. Neuroimage. 2013;69: 223–230. doi:10.1016/j.neuroimage.2012.12.038

62. Liu C, Downey RJ, Mu Y, Richer N, Hwang J, Shah VA, et al. Comparison of EEG Source Localization Using Simplified and Anatomically Accurate Head Models in Younger and Older Adults. IEEE Transactions on Neural Systems and Rehabilitation Engineering. 2023;31: 2591–2602. doi:10.1109/TNSRE.2023.3281356

63. Belli V de, Orcioli-Silva D, Beretta VS, Vitório R, Zampier VC, Nóbrega-Sousa P, et al. Prefrontal Cortical Activity During Preferred and Fast Walking in Young and Older Adults: An fNIRS Study. Neuroscience. 2021;473: 81–89. doi:10.1016/j.neuroscience.2021.08.019

64. Nagano H, Begg RK, Sparrow WA, Taylor S. A comparison of treadmill and overground walking effects on step cycle asymmetry in young and older individuals. Journal of Applied Biomechanics. 2013;29: 188–193. doi:10.1123/jab.29.2.188

65. Studnicki A, Ferris DP. Parieto-Occipital Electrocortical Dynamics during Real-World Table Tennis. eNeuro. 2023;10: ENEURO.0463-22.2023. doi:10.1523/ENEURO.0463-22.2023

66. Dal U, Erdogan T, Resitoglu B, Beydagi H. Determination of preferred walking speed on treadmill may lead to high oxygen cost on treadmill walking. Gait and Posture. 2010;31: 366–369. doi:10.1016/j.gaitpost.2010.01.006

67. Malatesta D, Canepa M, Menendez Fernandez A. The effect of treadmill and overground walking on preferred walking speed and gait kinematics in healthy, physically active older adults. Eur J Appl Physiol. 2017;117: 1833–1843. doi:10.1007/s00421-017-3672-3

68. Nordin AD, Hairston WD, Ferris DP. Dual-electrode motion artifact cancellation for mobile electroencephalography. J Neural Eng. 2018;15: 056024. doi:10.1088/1741-2552/aad7d7

69. Studnicki A, Downey RJ, Ferris DP. Characterizing and Removing Artifacts Using Dual- Layer EEG during Table Tennis. Sensors (Basel). 2022;22: 5867. doi:10.3390/s22155867

70. Downey RJ, Ferris DP. The iCanClean Algorithm: How to Remove Artifacts using Reference Noise Recordings. arXiv. 2022; 4. doi:10.48550/ARXIV.2201.11798

71. Gonsisko CB, Ferris DP, Downey RJ. iCanClean Improves Independent Component Analysis of Mobile Brain Imaging with EEG. Sensors. 2023;23: 928. doi:10.3390/s23020928

72. Pion-Tonachini L, Kreutz-Delgado K, Makeig S. ICLabel: An automated electroencephalographic independent component classifier, dataset, and website. Neuroimage. 2019;198: 181–197. doi:10.1016/j.neuroimage.2019.05.026

73. Palmer J, Kreutz-Delgado K, Makeig S. AMICA : An Adaptive Mixture of Independent Component Analyzers with Shared Components. 2011. Available: https://www.semanticscholar.org/paper/AMICA-%3A-An-Adaptive-Mixture-of-Independent-with-Palmer-Kreutz-Delgado/5774e96ad450c228400dc311f16caf1f20967c10

74. Nielsen JD, Madsen KH, Puonti O, Siebner HR, Bauer C, Madsen CG, et al. Automatic skull segmentation from MR images for realistic volume conductor models of the head: Assessment of the state-of-the-art. NeuroImage. 2018;174: 587–598. doi:10.1016/j.neuroimage.2018.03.001

75. Thammasan N, Miyakoshi M. Cross-Frequency Power-Power Coupling Analysis: A Useful Cross-Frequency Measure to Classify ICA-Decomposed EEG. Sensors. 2020;20: 7040. doi:10.3390/s20247040

76. Cohen J. Statistical Power Analysis for the Behavioral Sciences. Academic Press; 2013.

77. Benjamini Y, Hochberg Y. Controlling the False Discovery Rate: A Practical and Powerful Approach to Multiple Testing. Journal of the Royal Statistical Society: Series B (Methodological). 1995;57: 289–300. doi:10.1111/j.2517-6161.1995.tb02031.x

